# A network pharmacology-based approach and molecular docking study to explore the therapeutic potential of a nutraceutical formula (Vernolac) in the treatment of cancer

**DOI:** 10.1101/2025.09.06.674609

**Authors:** Sandani De Vass Gunawardane, Matheen Muhammadh Milhan, Poorni Chanuka Rathnayaka, Prabudhi S. Garusinghe, Kavishka S. Gunaratne, Anusha Kanagasundaram, T.M.D. Darshanamala, Duvinika Chalani Senevirathne, Shalini Kaushalya Wijerathne, R.P.C.D. Perera, Kanishka Sithira Senathilake, Umapriyathrashini Rajagopalan, Kamani Hemamala Tennakoon, Sameera Ranganath Samarakoon

## Abstract

Vernolac is a commercially available polyherbal nutraceutical capsule comprised of *Vernonia zeylanica* aerial parts, *Nigella sativa* seeds, *Hemidesmus indicus* roots, *Leucas zeylanica* aerial parts, and *Smilax glabra* rhizome. Different herbal formulations, organic extracts, and many isolated phytochemicals of the above plants have been reported to exhibit anticancer properties. However, the anticancer mechanisms of action of Vernolac, as a polyherbal formulation, remain unexplored. This study employed an integrative network pharmacology-based approach, complemented by *in vitro* experiments, to investigate the anticancer potential of Vernolac. Phytochemicals in Vernolac were retrieved from databases, screened for drug-likeness and oral bioavailability using SwissADME, yielding 155 drug-like phytochemicals, and their protein targets were predicted via SwissTargetPrediction. The intersection of targets of phytochemicals and cancer-related targets from GeneCards yielded 137 common targets. Protein-protein interaction analysis in STRING and Cytoscape identified key hub nodes, including AKT1, BCL2, CASP3, CTNNB1, EGFR, ESR1, GAPDH, HSP90AA1, HSP90AB1, IL6, JUN, SRC, STAT3, and TNF. Clustering, topology, and formula-herb-compound-target-disease and target-pathway networks highlighted key phytochemicals, including vernolactone, thymoquinone, quercetin, nigellidine, α-hederin, and carvacrol. Gene ontology (GO) and Kyoto Encyclopedia of Genes and Genomes (KEGG) enrichment revealed that the identified targets are significantly enriched in multiple cancer pathways. Molecular docking and dynamics simulations identified novel target-ligand interactions. Overall, network analysis suggests that Vernolac may exert anticancer effects through apoptosis induction, immune modulation, antioxidant, anti-inflammation, antiproliferative, and chemoradiosensitizing mechanisms. Moreover, Vernolac may exhibit chemoradioprotective potential by alleviating therapy-induced toxicity, supporting its promise as a potential adjunct to conventional cancer treatments. The Sulforhodamine B assay demonstrated selective antiproliferative activity of Vernolac against cancerous cells MCF-7 (IC_50_ = 54.01 ± 0.02 μg/mL), Caco-2 (IC_50_ = 85.52 ± 0.13 μg/mL), NTERA-2 cl.D1 (IC_50_ = 42.41 ± 0.06 μg/mL), and non-cancerous MCF-10A (IC_50_ = 803.5 ± 0.03 μg/mL). Novel target-ligand interactions identified via molecular docking and dynamics simulations, and the underlying mechanisms of Vernolac predicted in this study, require further validation through *in vitro* and *in vivo* experiments.

## Introduction

Cancer remains one of the leading causes of death worldwide. Despite significant development in early detection and treatment strategies, the global cancer burden continues to escalate [1,2]. While conventional treatments like surgery, chemotherapy, radiation, and targeted therapies remain standard approaches, they are frequently associated with unpleasant side effects. [3]. Furthermore, the emergence of therapy resistance further complicates treatment outcomes, leading to treatment failure [4]. These challenges underscore the urgent need for novel therapeutic strategies that are both effective and well-tolerated.

Natural products have gained significant attention in cancer therapy due to their broad-spectrum therapeutic possibilities [5]. Many natural products derived from plants, marine organisms, fungi, and microbes exhibit anticancer effects by modulating key cellular processes, including apoptosis, proliferation, angiogenesis, and immune regulation [6]. In this context, nutraceuticals have emerged as promising adjuncts to conventional cancer therapies [7]. These products are particularly attractive for their potential to exert multi-targeted effects while protecting healthy tissues, making them promising candidates for long-term use in cancer management [8].

Vernolac is a polyherbal nutraceutical currently in the Sri Lankan market for the management of cancer. This formulation comprised five medicinal plants with ethnomedical relevance, namely *Vernonia zeylanica* aerial parts, *Nigella sativa* seeds, *Smilax glabra* rhizome, *Leucas zeylanica* aerial parts, and *Hemidesmus indicus* roots [9]. Although the anticancer potential of most of the phytochemicals, several herbal formulations, and organic extracts of the aforementioned plants has been reported in previous studies [10–17], the anticancer mechanisms of Vernolac, as a polyherbal formulation, remain unexplored.

Network pharmacology is an emerging discipline in pharmacological research, offering a powerful platform to decode the complex interactions between compounds, targets, and disease-related pathways [18]. Unlike the traditional “one drug-one target” approach, network pharmacology embraces the multi-component, multi-target, and multi-pathway nature [18,19], particularly of polyherbal formulations like Vernolac.

In the present study, a systematic network pharmacology-based approach was employed to investigate the anticancer mechanisms of action of Vernolac. By integrating target prediction, protein-protein interaction analysis, and pathway analysis, this study aimed to explore the key targets and pathways modulated by Vernolac. Moreover, molecular docking and dynamics simulation studies were employed to further validate interactions between phytochemicals in Vernolac and key cancer targets. Additionally, we evaluated the cytotoxicity of Vernolac in both cancer cells and normal cells. The workflow of the network pharmacology approach is shown in Fig 1.

**Fig 1.**
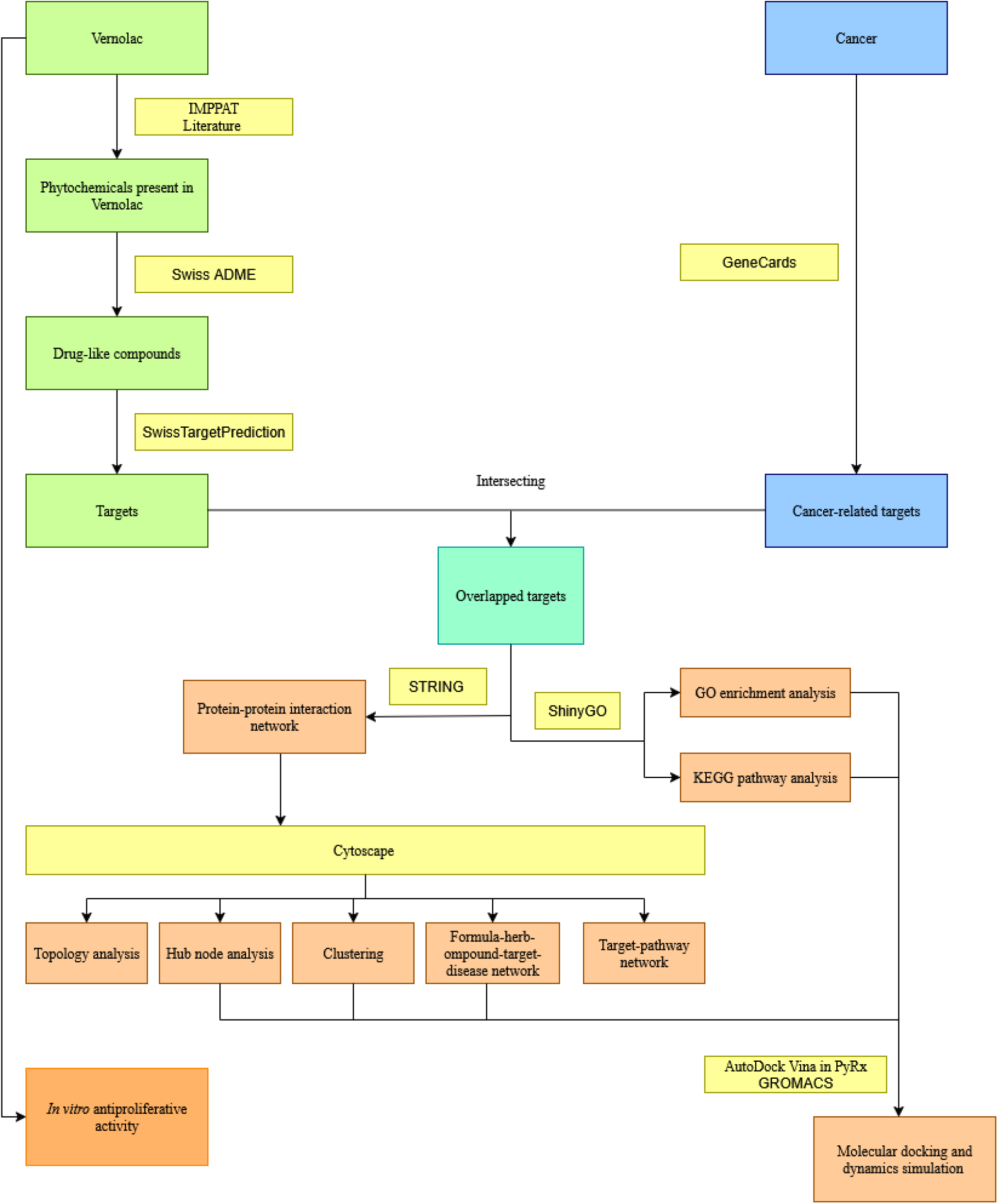
Workflow for investigation of the anticancer mechanisms of action of Vernolac in the treatment of cancer.

## Materials and methods

### Acquisition of phytochemicals in Vernolac

Vernolac is composed of *N. sativa*, *S. glabra*, *L. zeylanica*, *V. zeylanica*, and *H. indicus*. Compounds of *N. sativa*, *S. glabra and H. indicus* retrieved from the Indian Medicinal Plants, Phytochemistry and Therapeutics 2.0 (IMPPAT 2.0) database (https://cb.imsc.res.in/imppat/).

Literature mining also supplemented the information from the database. Since there is no information about *L. zeylanica* and *V. zeylanica* in the IMPPAT 2.0, their compounds were retrieved through a literature search. After removing duplicates, the chemical structure and canonical SMILES of each compound were obtained from the PubChem database (https://pubchem.ncbi.nlm.nih.gov/). The chemical structures of compounds for those that do not have information in the PubChem database were drawn by MarvinSketch ChemAxon (https://chemaxon.com/marvin).

### Prediction of drug-likeness of Vernolac compounds

Despite remarkable *in vitro* results, some phytochemicals have been shown to have minimal or negligible *in vivo* activities, which leads to poor absorption, consequently, low bioavailability [18]. Drug discovery widely uses absorption, distribution, metabolism, and excretion (ADME) studies to optimize the balance of properties required to convert leads into effective medications [20]. Therefore, ADME characteristics of phytochemicals must be considered. In the present study, drug likeness and bioavailability of compounds were predicted using SwissADME, a free web tool to evaluate pharmacokinetics, drug-likeness and medicinal chemistry friendliness of small molecules.

The chemical structure or canonical SMILES of each compound was imported into SwissADME (http://www.swissadme.ch/) to obtain drug-likeness status and bioavailability score. The active compounds were filtered by the criteria of the Lipinski rules for drug-likeness status: ‘Yes’ and the oral bioavailability score of ≥ 0.30. Compounds that align with the above criteria were selected for target prediction.

### Target prediction of compounds and cancer

The structure or SMILES of each phytochemical was imported into SwissTargetPrediction (http://www.swisstargetprediction.ch/), a free web tool that predicts the most probable protein targets of small molecules. The species was set to “*Homo sapiens*,” and the target data of each compound was exported in the CSV format, and the targets with a probability greater than zero were extracted for further analyses. The list of targets was imported to the UniProt database (https://www.uniprot.org/) to ensure standardization and consistency of protein names. Cancer-related genes were obtained by searching the GeneCards database (https://www.genecards.org/) with the keyword ‘Cancer’.

### Acquisition of common targets

The targets of Vernolac and cancer-related targets were uploaded to the Venny 2.1.0 online tool (https://bioinfogp.cnb.csic.es/tools/venny/) to obtain targets overlapped with cancer and Vernolac.

### Selection of core targets for further analyses

As the number of common targets was high, to identify core targets in the large protein-protein interaction (PPI) network, a stepwise filtering strategy was implemented based on network topology parameters, focusing on the degree centrality to prioritize highly connected and biologically significant nodes.

### Construction of the preliminary protein-protein interaction (PPI) network

The UniProt IDs of common targets of cancer and Vernolac were uploaded into the STRING database (https://string-db.org/, version 12.0) for initial PPI network construction, selecting the ‘multiple protein’ option on the database, and the species was selected as “*Homo sapiens*”. The confidence score was set to 0.9 to ensure high-confidence interactions. This threshold yielded a network consisting of 679 nodes (proteins) and 4934 edges (interactions).

### Network analysis and filtering

Cytoscape is an open-source software platform for analyzing and visualizing complex networks (https://cytoscape.org/). Cytoscape version 3.10.3, along with its various plugins, was utilized in this study.

The STRING PPI network was imported into Cytoscape for topology analysis. To identify core targets, the network topology was analyzed using the CytoNCA plugin (https://apps.cytoscape.org/apps/cytonca), a Cytoscape plugin integrating calculation, evaluation and visualization analysis for multiple centrality measures [21]. Weighted analysis was performed by incorporating interaction scores from the STRING database as edge weights to reflect the reliability and strength of protein-protein interactions.

Degree centrality (DC), a measure of the number of direct connections (edges) a node has with other nodes in the network, was used to identify core targets within the network. The DC highlights hub proteins that are likely to be biologically significant due to their extensive connectivity within the PPI network, often participating in multiple pathways and regulating essential biological processes.

Nodes with a DC ≥ 20 were selected to ensure the retention of core proteins with significant connectivity. This cutoff was selected based on the distribution of DC values in the network and its ability to balance the retention of biologically important targets while simplifying the network. After filtering, the network was reduced to 137 nodes representing the core targets with high connectivity. The downstream analyses were conducted on the refined network that contains 137 nodes and their interactions.

### Construction and network analyses of the core target PPI network

The core target PPI network was constructed using the STRING database. The core target PPI network was imported into Cytoscape, and network parameters were analyzed. Topology parameters, including betweenness centrality (BC), closeness centrality (CC), and degree centrality (DC), were analyzed with weight using the CytoNCA plugin.

### Identification of hub nodes

The CytoHubba plugin (https://apps.cytoscape.org/apps/cytohubba) was used to identify top hub proteins. In this study, five different methods of the CytoHubba plugin were employed, including two global rank methods (betweenness and closeness) and three local rank methods (degree, maximal clique centrality (MCC), and maximum neighborhood component (MNC)), as described in a previous study [19]. While global rank methods consider the relationship between each node and the entire network, local rank methods calculate the score of a node within a network based on the proximity of the node to its direct neighbors [22]. The top 20 hub nodes were obtained using each method, and the common hub proteins were identified by combining the five CytoHubba methods.

### Identification of clusters of the PPI network

In complex PPI networks, some proteins are densely connected, forming clusters. Proteins in a cluster have the same or similar functions and frequently interact with each other, creating a functional module within the network. The clustering of the PPI network was performed using the Molecular Complex Detection (MCODE) Cytoscape plugin (https://apps.cytoscape.org/apps/mcode). MCODE is an algorithm that effectively finds densely connected groups within a molecular interaction network, based on interaction data.

The PPI network was imported to Cytoscape for clustering using the MCODE. Parameters, including degree cutoff (2), node score cutoff (0.2), k-core cutoff (2), and max depth (100), were adjusted, and the haircut filter was enabled. The clusters were visualized, and the topology parameters of each cluster were calculated using MCODE and the Analyze Network Cytoscape plugin.

### Construction and topology analysis of Formula-Herb-Compound-Target-Disease network

The formula-herb-compound-target-disease network was constructed using Cytoscape 3.10.3 software to visualize the complex interactions between compounds and cancer-related protein targets. Separate data files were prepared for the formula-herb, herb-compound, compound-target, target-target, and target-disease relationships using Microsoft Excel. The data files were then imported into Cytoscape to merge networks, constructing the interaction network of Vernolac in cancer treatment. In this graphical network, the formula, herbs, compounds, targets, and disease were denoted as nodes, and the edges represent formula-herb-compound-target-disease interactions. The Analyze Network plug-in in Cytoscape was used to assess the topology parameters (degree) of the network.

### GO and KEGG pathway enrichment analysis

Enrichment analysis on the core target proteins was carried out using the ShinyGO (version 0.81) web-based application (https://bioinformatics.sdstate.edu/go/). The ShinyGO is a large annotation and pathway database for Gene Ontology (GO) functional enrichment analysis and Kyoto Encyclopedia of Genes and Genomes (KEGG) pathway enrichment analysis [23].

The three major GO terms analyzed were the biological processes (BP), cellular components (CC), and molecular functions (MF). KEGG pathway enrichment analysis was used to identify molecular biological pathways enriched by the core targets of Vernolac and cancer. The false discovery rate (FDR) cutoff was set to 0.05, and the top 20 BP, CC, and MF GO terms and the top 20 KEGG pathways were selected based on the fold enrichment. The bubble charts with fold enrichment, FDR and gene counts were used for graphical representation of the top 20 GO terms and KEGG pathways.

### Construction of the target-pathway network

To effectively illustrate the relationship between protein targets and the highly enriched pathways in which they are involved, the top 20 KEGG pathways were selected to construct a target-pathway network. Target-target and target-pathway data sheets were prepared in Microsoft Excel. Then, Cytoscape was used to merge the imported files and generate a comprehensive target-pathway network that highlights key biological connections.

### Molecular docking

The X-ray crystal structures of the most significant targets identified through network analysis were obtained from the RCSB Protein Data Bank (https://www.rcsb.org/). UCSF Chimera 1.18 software was used to preprocess receptor proteins by removing ligands and water molecules. Subsequent preparation steps included hydrogenation, charge calculation and energy minimization. The energy minimization process targeted the nearest local minima using the Amber ff4SB force field with 100 steepest descent steps, resulting in the finalized PDB structures for virtual screening.

For the phytochemicals (ligands), the majority of their 3D structures were obtained from the PubChem database (https://pubchem.ncbi.nlm.nih.gov/) and the ChEMBL database (https://www.ebi.ac.uk/chembl/) in SDF format. For phytochemicals not available in these databases, 3D structures were generated using OpenBabel software from 2D structures drawn with Marvin JS software. The structures were subjected to geometry optimization using the mmff94 force field in OpenBabel (PyRx).

Docking studies were conducted with AutoDock Vina in PyRx, utilizing a Lamarckian genetic algorithm-based scoring function. Docking parameters were set to include the active sites of the target proteins where the compounds bind. Additionally, binding interaction within the active site pocket and bond analysis were thoroughly examined.

Post-docking studies, such as molecular dynamics simulations, were conducted to further substantiate the docking results. All docking scores for the target proteins adhered to a cutoff of -7.0 kcal/mol. In addition to this cutoff, consideration was also given to thymoquinone, a previously reported major anticancer compound.

### Molecular dynamics (MD) simulation

Molecular dynamics simulations were conducted using GROMACS 2022.4 software to analyze the interaction and stability of target proteins with phytochemicals. The docked complexes with the best pose based on the docking scores were subjected to MD simulations. The CHARMM27 force field was applied to model the receptor proteins and compounds, and the ligand topologies were generated using the SwissParam web server (https://old.swissparam.ch/).

A triclinic box was configured for all systems with a minimum distance of 1.0 nm between the protein complex and the box edge. Systems were solvated using the TIP3P water model and neutralized with appropriate sodium and chloride ions. Energy minimization was performed using the steepest descent algorithm. Equilibration phases were carried out under the Canonical ensemble (NVT) and isothermal-isobaric ensemble (NPT) conditions at 310 K and 1 bar for 100 ps.

A production run of 100 ns was performed for each system. All trajectories obtained from the simulations were analyzed by root mean square deviation (RMSD) and root mean square fluctuation (RMSF).

#### Cell Culture

All cell lines were cultured and maintained according to the American Type Culture Collection (ATCC) guidelines. The cancer cell lines MCF-7 (ATCC HTB-22), Caco-2 (ATCC HTB-37), and NTERA 2 cl.D1 (ATCC CRL-1973), cultured in Dulbecco’s Modified Eagle Medium (DMEM) and MCF-10A (ATCC CRL-10317) cells were cultured in Mammary Epithelial Cell Basal Medium (MEBM). Fetal bovine serum (FBS) 10%, 50 IU/mL penicillin, and 50 µg/mL streptomycin antibiotic mixtures were added to the culture medium according to ATCC guidelines. All cell lines were maintained in a 95% air and 5% CO_2_ atmosphere with 95% humidity at 37 °C. The cells were maintained by sub-culturing in 25 cm^2^ (T25) cell culture flasks, and cells growing in the exponential phase were used for the SRB assay.

### Preparation of the sample

The polyherbal formulation comprised of aforementioned plant materials was prepared using the supercritical carbon dioxide (CO_2_) method at 42 °C, under 30 MPa pressure at a flow rate of 4 mL/min for 1 h. The collected extract was dissolved in 0.08% DMSO and serially diluted with media at concentrations ranging from 6.25 to 400 µg/mL.

### Sulforhodamine B (SRB) assay

The antiproliferative effect of Vernolac was evaluated using the Sulforhodamine B (SRB) assay, following the protocol described by Rajagopalan et al. [24]. Cells were seeded into 96-well plates (10^4^ cells/well in 100 µL of media) and allowed to attach for 24 h at 37 °C in a humidified atmosphere containing 5% CO_2_. Cells were treated with different concentrations of Vernolac and incubated for 24 h and 48 h. Cells in the control group received only media containing 0.08% DMSO. Following treatment, cells were washed twice with phosphate-buffered saline (PBS), fixed by adding 50% ice-cold trichloroacetic acid (TCA), and incubated at 4 °C for 1 h. The plates were washed with tap water five times and air-dried. Subsequently, 50 µL of 0.4% (w/v) SRB dye prepared in 1% acetic acid was added to each well and incubated in the dark at room temperature for 15 min. Excess dye was removed by washing the wells with 1% acetic acid. The plates were air-dried, and the protein-bound SRB dye was solubilized by adding 100 µL of 10 mM unbuffered Tris base to each well. The plates were then placed on an orbital shaker for 1 h at room temperature. The absorbance was recorded at 540 nm using a Synergy HT microplate reader, BioTek Instruments, USA, and the percentage cell viability was calculated using the following formula: Percentage cell viability = [(A_s_ – A_b_) / (A_c_ – A_b_)] × 100, where A_s_ = absorbance of the sample, A_c_ = absorbance of the negative control, and A_b_ = absorbance of the blank. The IC_50_ values were calculated by non-linear curve fit analysis using GraphPad Prism 8.0.1 (GraphPad Software Inc., San Diego, CA, USA) with R^2^ >0.9 and p >0.5. All data were presented as mean ± SD, obtained from three independent replicates (n = 3) for each cell line.

## Results

### Screening of phytochemicals of Vernolac

After an extensive search of compounds of Vernolac through the IMPAAT database and literature mining, 403 compounds were retrieved, including 238, 28, 49, 01, and 87 compounds in *N. sativa, S. glabra, L. zeylanica, V. zeylanica, and H. indicus*, respectively. Some herbs shared similar compounds (Fig 2). The overlapping regions indicate the presence of shared phytochemicals in all five herbs, suggesting possible synergistic effects that may contribute to the formulation’s overall anticancer potential.

**Fig 2.**
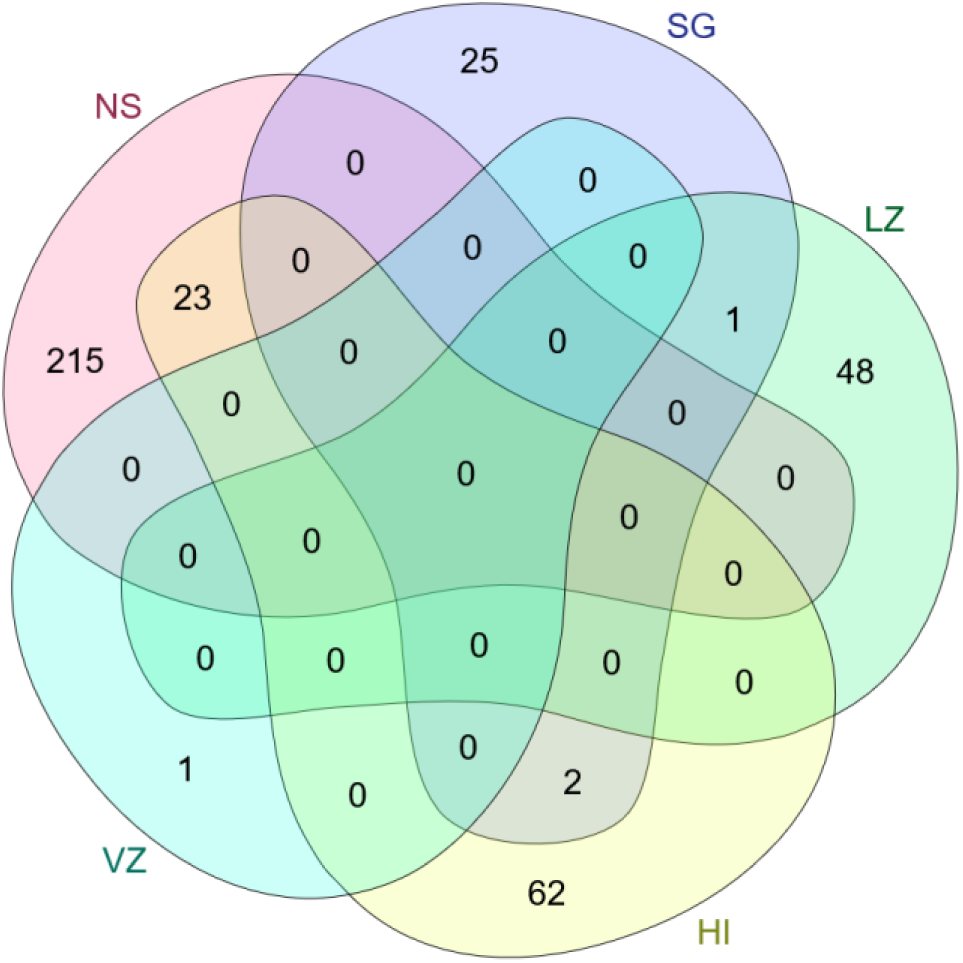
Venn diagram representing the distribution of phytochemicals among the five herbs in Vernolac. NS- *Nigella sativa*, SG- *Smilax glabra*, LZ- *Leucas zeylanica*, HI- *Hemidesmus indicus*, VZ- *Vernonia zeylanica*

### Drug likeness and oral bioavailability of Vernolac compounds

A total of 489 compounds in Vernolac obtained through literature and database mining were screened for drug-likeness and oral bioavailability using SwissADME. Based on Lipinski’s rule of five (≤2 violations) and a bioavailability score >0.30, 155 compounds were identified as drug-like and prioritized for further analyses.

### Target prediction of compounds and cancer

Target prediction for drug-like compounds selected from SwissADME predictions was performed, obtaining 500, 427, 629, 422, and 29 targets for *N. sativa*, *S. glabra*, *L. zeylanica*, *H. indicus*, and *V. zeylanica*, respectively. After removing duplicates, a total of 927 targets were obtained. To identify cancer targets, a search was conducted in the GeneCards database, obtaining 18,803 cancer-related protein-coding genes.

### Acquisition of common targets

Using Venny 2.1.0, potential Vernolac targets relevant to cancer were identified by intersecting them with cancer-related targets. A total of 876 targets associated with both Vernolac and cancer were identified (Fig 3).

**Fig 3.**
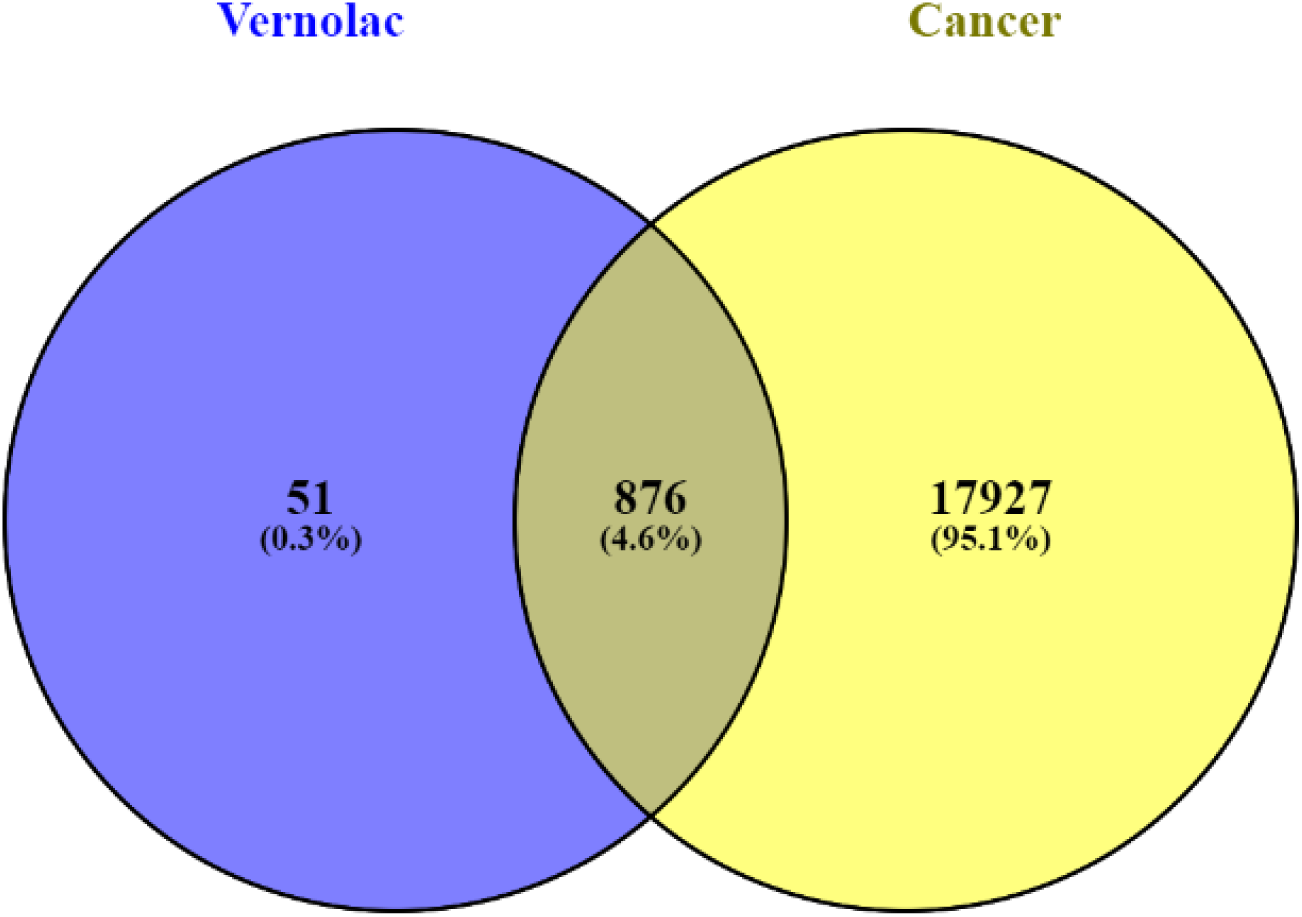
Venn diagram illustrating the intersection between predicted targets of Vernolac and cancer targets.

### Core target selection for further analyses

A stepwise filtering strategy was applied to extract the core targets from the PPI network, prioritizing highly connected and biologically significant nodes. To ensure the inclusion of only strongly supported protein-protein interactions, the initial PPI network was constructed using high-confidence interactions (confidence score >0.9) from the STRING database. This resulted in a network consisting of 679 nodes and 4934 edges. Disconnected nodes were hidden in the preliminary PPI network depicted in Fig 4.

**Fig 4.**
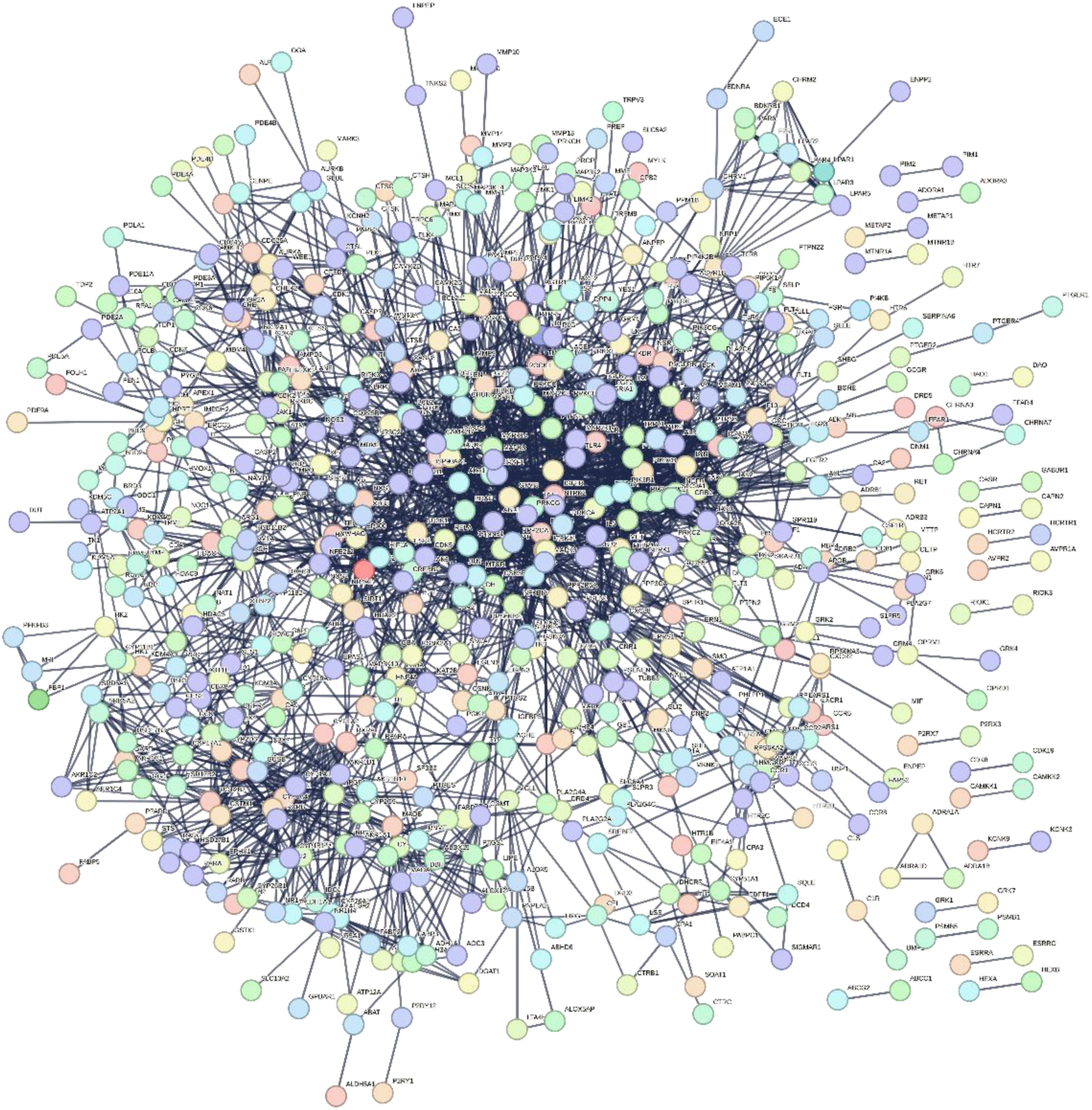
Preliminary protein-protein interaction (PPI) network. Nodes-proteins, Edges- interactions

To further refine the network and select core proteins, the DC topology parameter was applied as the primary filtering criterion. DC is the number of edges/interactions of a particular node [25]. Nodes with high DC value represent hub proteins with significant interactions with many other proteins in the network, signifying their potential importance in cellular processes. Therefore, targets with DC ≥20 were selected. After applying this threshold, the number of targets was reduced to 137 nodes, representing central nodes located at the center of the network. These 137 core targets were selected for further analyses, including topology analysis, clustering, pathway enrichment, and hub node identification, to explore their role in the molecular mechanisms.

### Core target PPI network

The core target PPI network was constructed using the STRING database with 137 key targets (Fig 5). The resulting network consisted of 137 nodes and 3183 edges. According to the network statistics, the average node degree and average local clustering coefficient were 46.5 and 0.648, respectively. The PPI enrichment p-value was < 1.0e-16, indicating that the observed interactions were significantly more than expected by random chance and a strong functional association among the targets.

**Fig 5.**
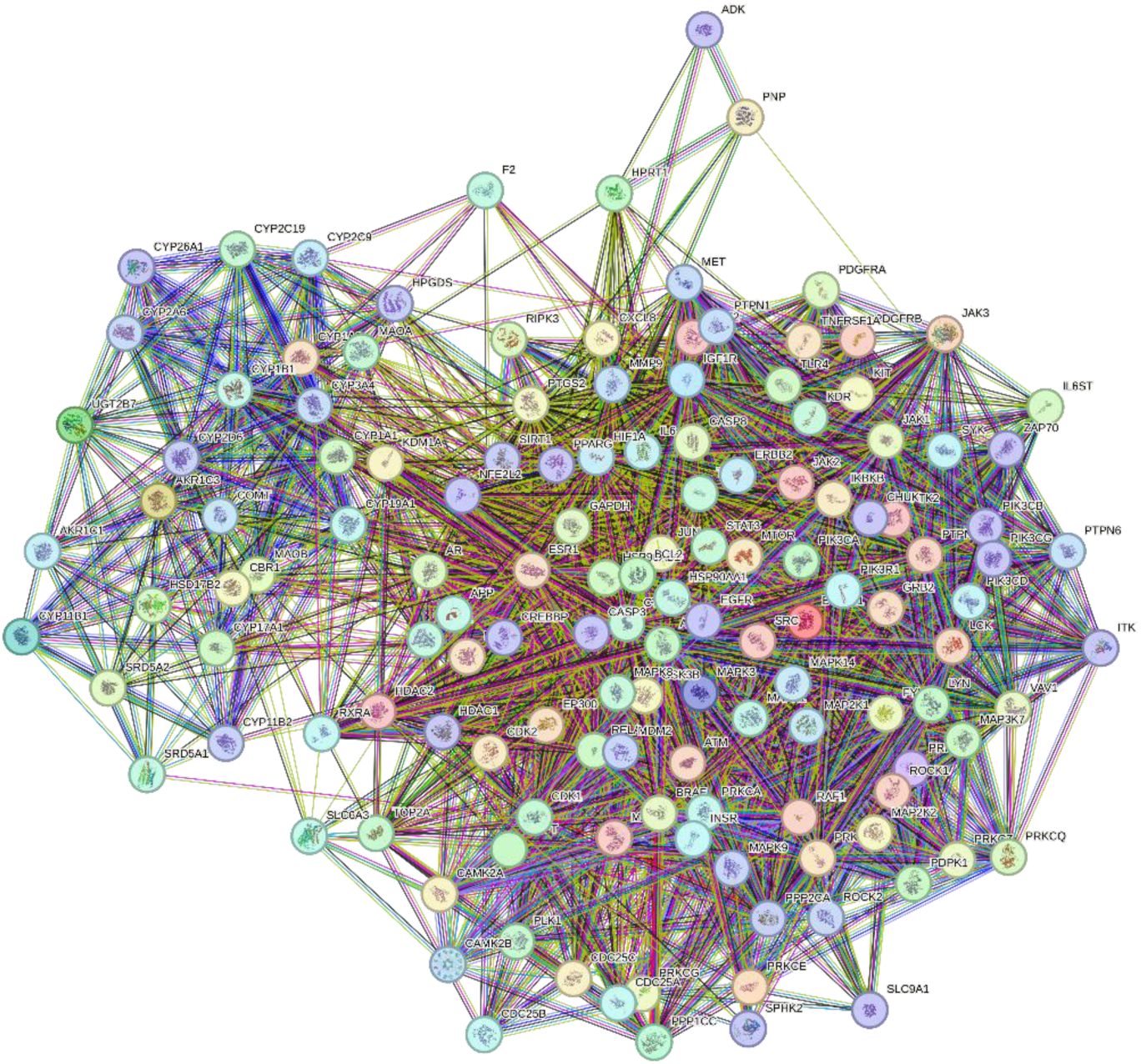
Core target Protein-Protein Interaction network. Nodes represent target proteins. Interactions between proteins are presented by edges.

### Topological analysis of the PPI network

The key nodes with DC>100 are presented in Table 1, including their respective BC and CC values. According to the CytoNCA analysis, the top 20 nodes include AKT1, SRC, EGFR, STAT3, CTNNB1, HSP90AA1, ESR1, JUN, MAPK1, BCL2, CASP3, IL6, MAPK3, GAPDH, TNF, PIK3CA, HSP90AB1, PIK3R1, and JAK2. Results of CytoNCA topology analysis of 137 core target proteins are provided in the S1 table.

**Table 1.** CytoNCA topology analysis of the top 20 core target proteins.

| Target | Degree centrality | Betweenness centrality | Closeness centrality |
| --- | --- | --- | --- |
| AKT1 | 154.96 | 1766 | 0.6031069 |
| SRC | 151.38 | 449 | 0.59821291 |
| EGFR | 151.24 | 134 | 0.60475951 |
| STAT3 | 148.51 | 28 | 0.59421991 |
| CTNNB1 | 140.32 | 934 | 0.57684926 |
| HSP90AA1 | 136.88 | 226 | 0.56991102 |
| ESR1 | 133.19 | 1334 | 0.58049449 |
| JUN | 131.76 | 264 | 0.56165885 |
| MAPK1 | 129.59 | 0.5 | 0.55806456 |
| BCL2 | 128.02 | 248 | 0.5489157 |
| CASP3 | 125.30 | 65.5 | 0.54274189 |
| IL6 | 123.82 | 325 | 0.53371403 |
| MAPK3 | 123.47 | 33 | 0.5512886 |
| GAPDH | 123.12 | 913 | 0.54348646 |
| TNF | 122.80 | 528.5 | 0.52283423 |
| PIK3CA | 120.52 | 14 | 0.52999167 |
| HSP90AB1 | 117.66 | 68 | 0.53992448 |
| PIK3R1 | 109.57 | 221 | 0.52096769 |
| JAK2 | 109.47 | 70 | 0.49898347 |
| MTOR | 108.27 | 36 | 0.50260174 |

In addition to degree centrality, closeness centrality (CC) and betweenness centrality (BC) were analyzed to further evaluate the significance of nodes in the network. Closeness centrality (CC) is defined as the reciprocal of the average length of the shortest paths between a node and all other nodes in the network. Higher CC values indicate that a node is more centrally located, allowing for faster information or signal transmission across the network. Betweenness centrality (BC) measures how often a node appears on the shortest paths between other node pairs in the network. A higher BC value suggests that the node acts as a critical bridge in the network, playing a key role in signal transduction and molecular interactions.

### Identification of hub nodes

To identify hub nodes in the PPI network, a multi-method approach was applied using the CytoHubba plug-in. Five different methods in the CytoHubba, including MCC, closeness, MNC, degree, and betweenness, were used to rank nodes. As Fig 6A-E shows, the top 20 nodes were selected for each method. Table 2 summarizes the top 20 nodes with scores for each method. The top-ranked nodes from all five methods were compared using a Venn diagram (Fig 6F) to identify hub nodes. The intersection of five methods resulted in the identification of 14 hub proteins that were common across all methods. The 14 intersecting hub proteins include, AKT1, BCL2, CASP3, CTNNB1, EGFR, ESR1, GAPDH, HSP90AA1, HSP90AB1, IL6, JUN, SRC, STAT3, and TNF. These proteins are highly connected within the network, indicating their functional significance and potential role as key regulators in cancer-related pathways.

**Fig 6.**
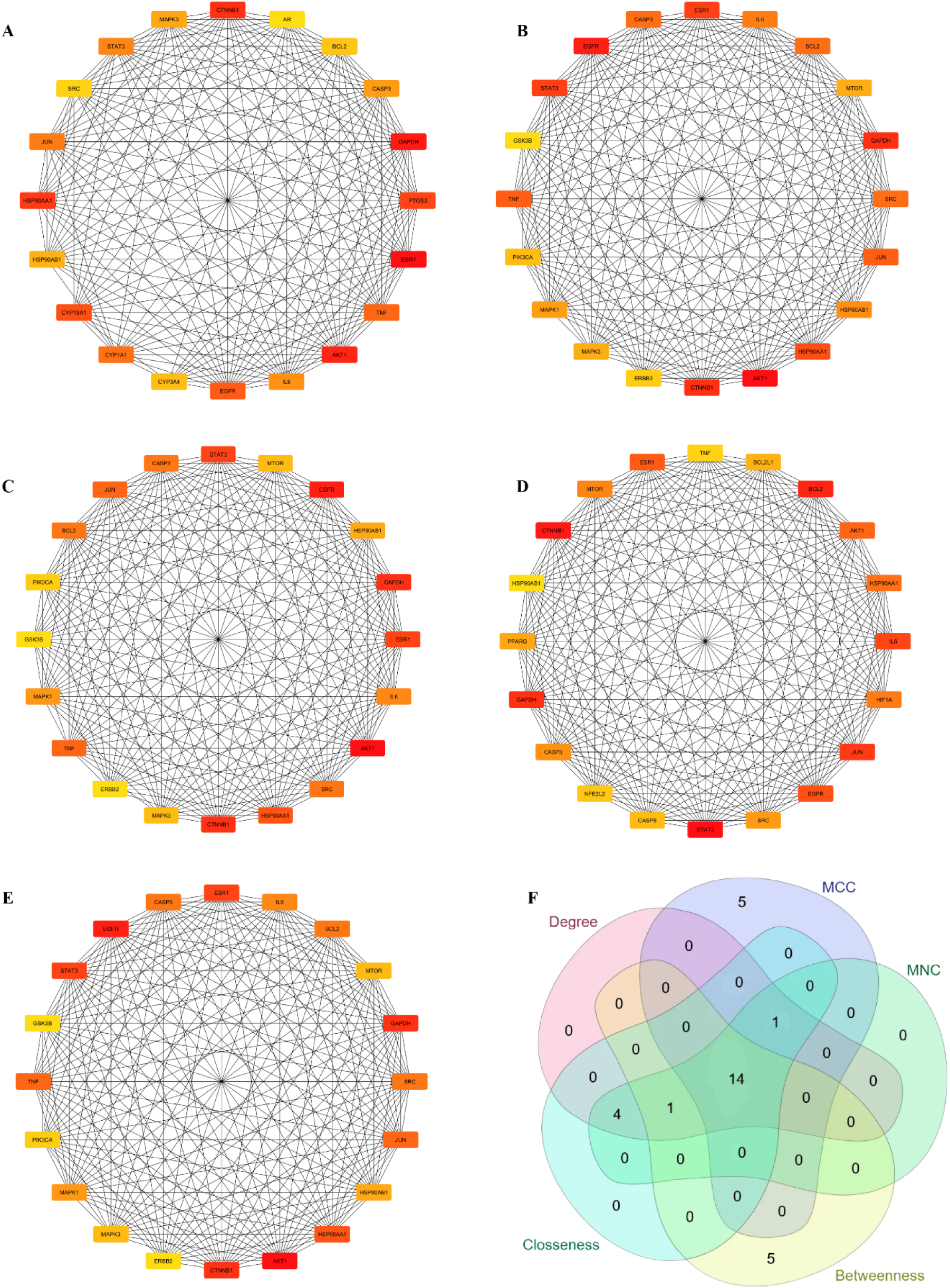
Key subnetworks of the top 20 hub nodes identified using five different algorithms in CytoHubba and their intersection. (A) Betweenness; (B) Closeness; (C) Degree; (D) MCC; (E) MNC; (F) The intersection of five methods. The color intensity of nodes represents the degree of interaction (red = highest score, yellow = lowest score). Rectangles and edges represent proteins and protein-protein interactions, respectively.

**Table 2.**
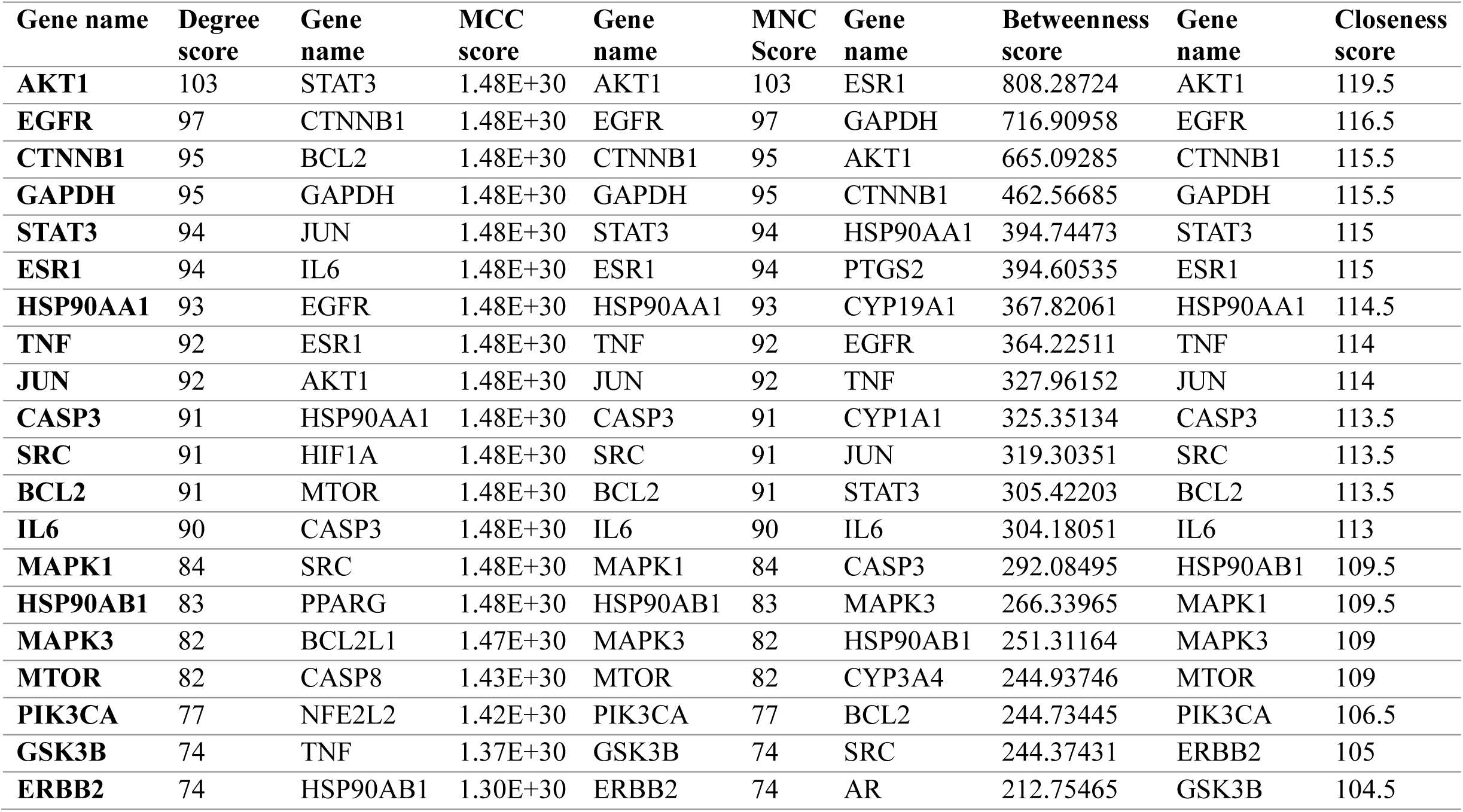
Top 20 hub proteins identified using five centrality algorithms in CytoHubba.

### Identification of core clusters

The MCODE analysis identified 7 clusters within the core PPI network, highlighting densely connected functional clusters (Fig 7). Each cluster represents potential functional modules associated with key biological processes. Results of MCODE analysis, including MCODE status, MCODE score, and degree, are presented in the S2 table.

**Fig 7.**
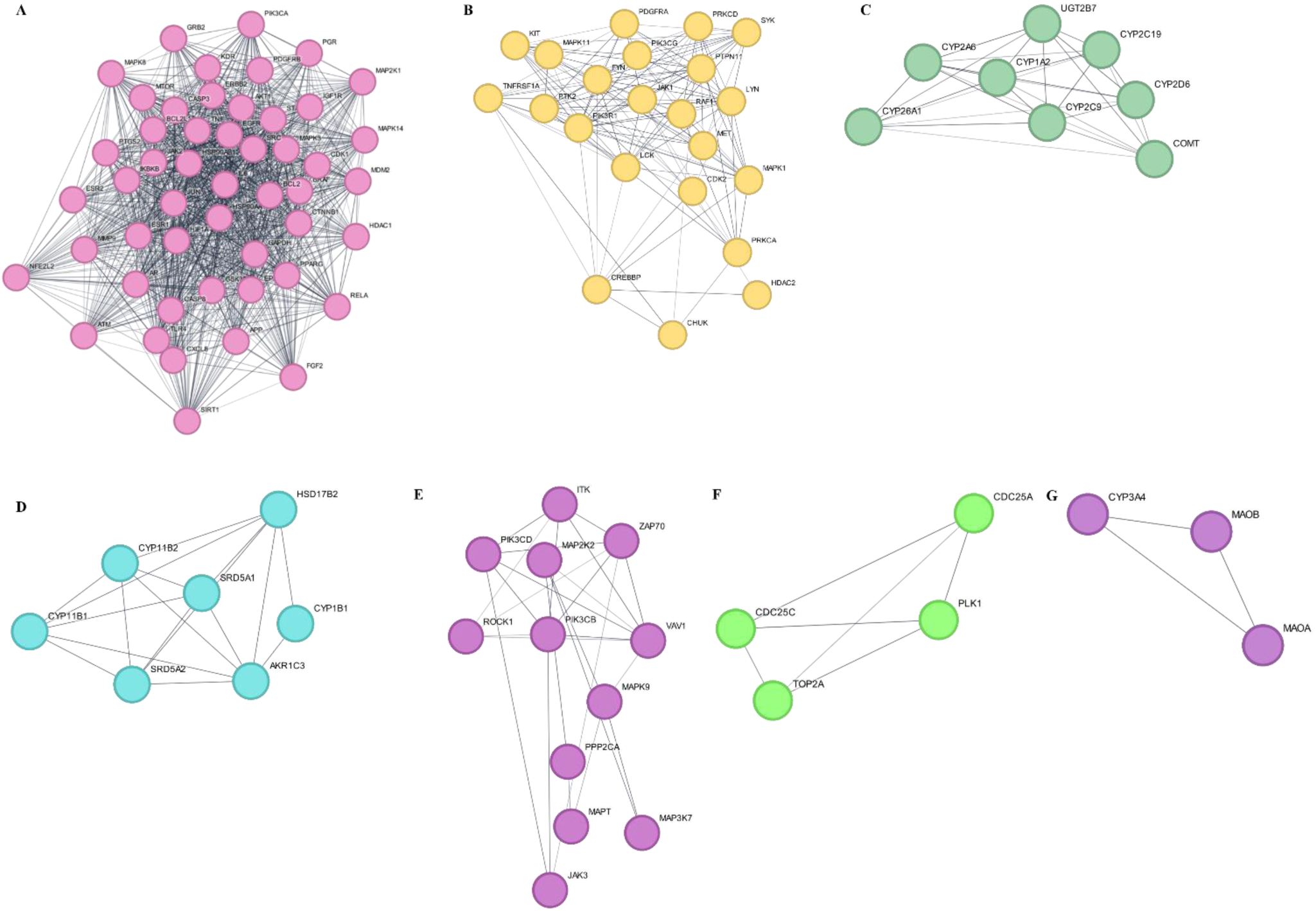
Clustering of the protein–protein interaction (PPI) network using the MCODE analysis. (A) Cluster 1, (B) Cluster 2, (C) Cluster 3, (D) Cluster 4, (E) Cluster 5, (F) Cluster 6, (G) Cluster. Colored circles and edges represent the proteins and protein-protein interactions, respectively.

### Formula-herb-compound-target-disease network construction and analysis

A network was constructed using Cytoscape 3.10.3 to visualize interactions between herbs, compounds, and cancer-related targets. The network comprised 265 nodes and 4709 edges (Fig 8). The topology analysis of the network revealed that one compound can target multiple genes, and one gene can be targeted by multiple compounds. Among all the compounds, the top 10 with the highest degree were carvacrol, cycloartenol, 24-methylene-cycloartanol, 4-terpineol, campesterol, cycloeucalenol, dithymoquinone, gramisterol, lophenol, and nigellicine, highlighting their significant involvement in the regulation of multiple cancer-related targets. The AR was the gene associated with the highest number of compounds, followed by high-degree targets with degree ≥90, including CYP19A1, JAK1, JAK2, PRKCA, SLC6A3, CYP2C19, PTPN1, ESR1, UGT2B7, PTGS2, CYP17A1, and ESR2. As the network illustrates, the interactions between compounds of Vernolac and cancer-related targets signify the therapeutic effect of Vernolac through synergistic actions of multiple compounds and targets.

**Fig 8.**
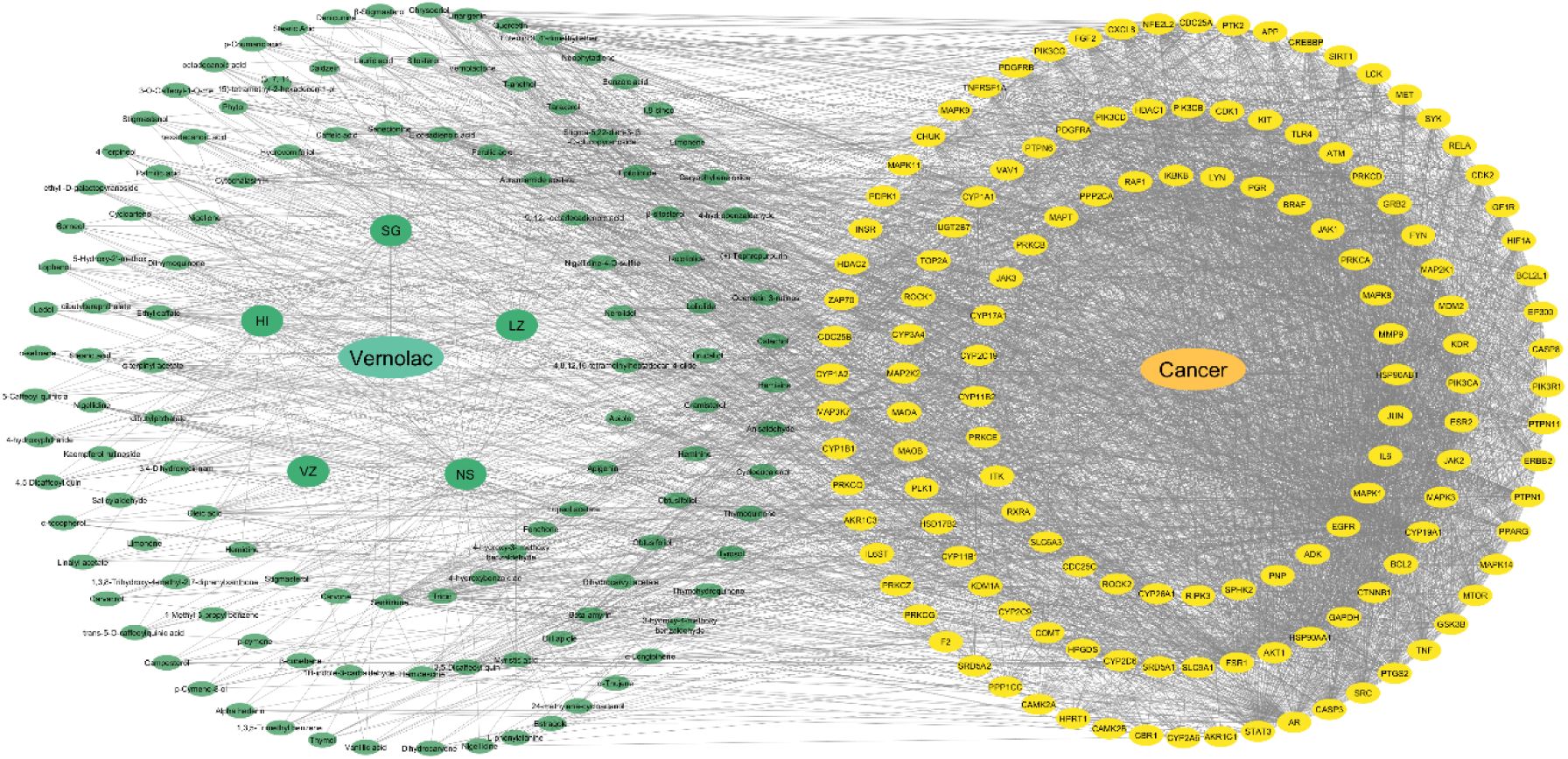
Formula-Herb-Compound-Target-Disease network. The central green ellipse represents the Vernolac formula, connected to green circles (herbs) and surrounding green ellipses (compounds). Yellow ellipses indicate cancer (center) and associated targets (outer rings), illustrating the multi-component, multi-target nature of Vernolac in cancer therapy. Edges represent interactions between each component in the network.

### GO term enrichment analysis

In the GO term enrichment analysis, a total of 1000 BP terms, 276 CC terms, and 523 MF terms were identified. The top 20 enriched terms in each category are presented in Fig 9 and Table 3. According to the results, the BP terms were mainly related to the immune regulation, cellular responses, metabolic pathways, cellular transport processes, and cellular differentiation. The enriched CC terms highlight membrane-bound signaling complexes, cytoplasmic structures, chromatin regulators, and neuronal components. The MF GO terms are primarily enriched in kinase activity, signal transduction, receptor interactions, and enzymatic activities.

**Fig 9.**
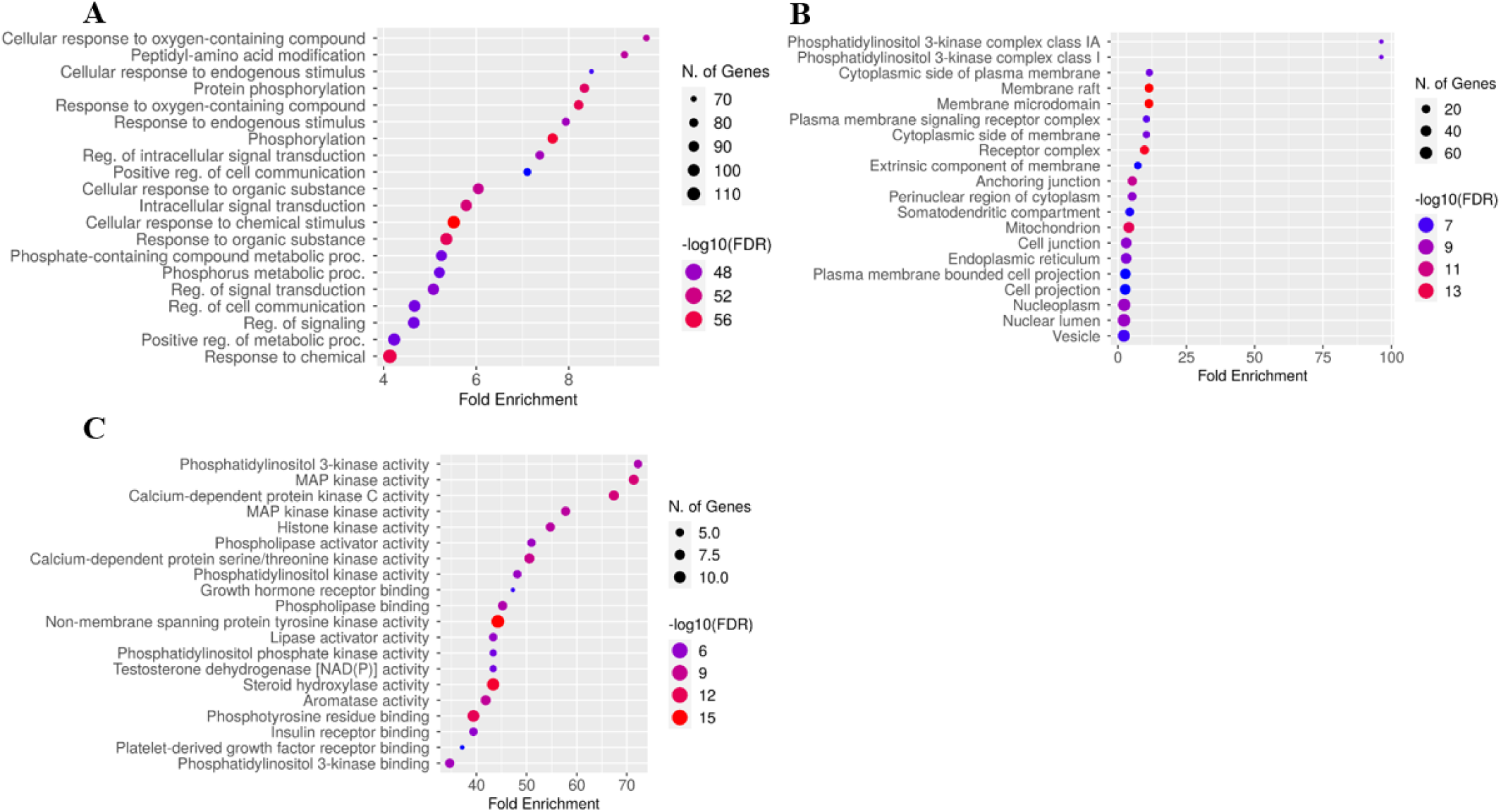
GO enrichment analysis of core targets. (A) GO biological process terms. (B) GO cellular process terms. (C) GO molecular function terms. The size of each dot corresponds to the number of genes annotated in the entry. The color of each dot corresponds to the false discovery rate (FDR).

**Table 3.** GO enrichment analysis.

| Category | Term | Genes | Count | Enrichment FDR |
| --- | --- | --- | --- | --- |
| Biological processes | GO:0060439 - Trachea morphogenesis | MAPK1, MAPK3, MAP2K2, CTNNB1, MAP2K1 | 5 | 2.04E-08 |
|  | GO:0097267 - Omega-hydroxylase P450 pathway | CYP1B1, CYP2C9, CYP1A1, CYP1A2, CYP2C19 | 5 | 2.04E-08 |
|  | GO:0002223 - Stimulatory C-type lectin receptor signaling pathway | SYK, CREBBP, FYN, EP300, IKBKB, MAP3K7, SRC, LYN | 8 | 2.45E-12 |
|  | GO:1990840 - Response to lectin | SYK, CREBBP, FYN, EP300, IKBKB, MAP3K7, SRC, LYN | 8 | 2.45E-12 |
|  | GO:1990858 - Cellular response to lectin | SYK, CREBBP, FYN, EP300, IKBKB, MAP3K7, SRC, LYN | 8 | 2.45E-12 |
|  | GO:0060020 - Bergmann glial cell differentiation | MAPK1, MAPK3, MAP2K1, PTPN11 | 4 | 1.88E-06 |
|  | GO:0097709 - Connective tissue replacement | HIF1A, PPARG, ROCK1, ROCK2 | 4 | 1.88E-06 |
|  | GO:0048308 - Organelle inheritance | MAPK1, MAPK3, CDK1, MAP2K2, MAP2K1 | 5 | 1.16E-07 |
|  | GO:0048313 - Golgi inheritance | MAPK1, MAPK3, CDK1, MAP2K2, MAP2K1 | 5 | 1.16E-07 |
|  | GO:0019373 - Epoxygenase P450 pathway | CYP2C9, CYP2C19, CYP2A6, CYP1B1, CYP1A1, CYP1A2 | 6 | 6.93E-09 |
|  | GO:0070102 - Interleukin-6-mediated signaling pathway | IL6ST, IL6, STAT3, SRC, JAK2, JAK1 | 6 | 6.93E-09 |
|  | GO:0035994 - Response to muscle stretch | MAPK14, PIK3CA, RAF1, RELA, JUN, SLC9A1, CTNNB1, PTK2 | 8 | 1.66E-11 |
|  | GO:1904385 - Cellular response to angiotensin | CAMK2A, PRKCD, NFE2L2, IGF1R, SRC, ROCK1, RELA, ROCK2 | 8 | 1.66E-11 |
|  | GO:1990776 - Response to angiotensin | CAMK2A, PRKCD, ROCK1, PTGS2, NFE2L2, IGF1R, SRC, RELA, ROCK | 9 | 9.60E-13 |
|  | GO:0038095 - Fc-epsilon receptor signaling pathway | MAPK9, PRKCQ, IKBKB, MAPK8, MAP3K7, VAV1, SYK, LYN | 8 | 2.30E-11 |
|  | GO:0060396 - Growth hormone receptor signaling pathway | JAK2, JAK3, JAK1, STAT3, PIK3R1, PTK2, PTPN1, LYN | 8 | 2.30E-11 |
|  | GO:0071378 - Cellular response to growth hormone stimulus | JAK2, JAK3, JAK1, STAT3, PIK3R1, PTK2, PTPN1, LYN | 8 | 2.30E-11 |
|  | GO:2000641 - Reg. of early endosome to late endosome transport | MAPK1, MAPK3, SRC, MAP2K2, MAP2K1 | 5 | 2.23E-07 |
|  | GO:0002433 - Immune response-regulating cell surface receptor signaling pathway in | FYN, VAV1, PRKCD, SYK, PTK2, PRKCE, SRC, LYN | 8 | 3.13E-11 |
|  | GO:0038096 - Fc-gamma receptor signaling pathway involved in phagocytosis | FYN, VAV1, PRKCD, SYK, PTK2, PRKCE, SRC, LYN | 8 | 3.13E-11 |
| Cellular components | GO:0099091 - Postsynaptic specialization intracellular component | CTNNB1, SRC, LYN | 3 | 0.000201 |
|  | GO:0008385 - Kappa kinase complex | IKBKB, CHUK, MAP3K7 | 3 | 0.00033 |
|  | GO:0044292 - Dendrite terminus | HSP90AA1, HSP90AB1, SRC | 3 | 0.0004 |
|  | GO:0042101 - T cell receptor complex | ZAP70, SYK, PTPN6 | 3 | 0.000707 |
|  | GO:0035631 - CD40 receptor complex | IKBKB, CHUK | 2 | 0.007999 |
|  | GO:0005942 - Phosphatidylinositol 3-kinase complex | PIK3CB, PIK3CG, PIK3CA, PIK3R1, PIK3CD | 5 | 8.38E-06 |
|  | GO:0031143 - Pseudopodium | MAPK1, MAPK3, RAF1 | 3 | 0.000972 |
|  | GO:0030877 - Beta-catenin destruction complex | GSK3B, CTNNB1 | 2 | 0.009333 |
|  | GO:0099522 - Cytosolic region | PRKCG, PRKCB | 2 | 0.013034 |
|  | GO:1990909 - Wnt signalosome | CTNNB1, GSK3B | 2 | 0.013034 |
|  | GO:0016581 - NuRD complex | HDAC1, HDAC2 | 2 | 0.014363 |
|  | GO:0090545 - CHD-type complex | HDAC1, HDAC2 | 2 | 0.014363 |
|  | GO:0097524 - Sperm plasma membrane | HSP90AA1, HSP90AB1 | 2 | 0.014363 |
|  | GO:0035098 ESC/E(Z) complex | SIRT1, HDAC2 | 2 | 0.015598 |
|  | GO:0044214 - Spanning component of plasma membrane | EGFR, ERBB2 | 2 | 0.015598 |
|  | GO:0044305 - Calyx of Held | PRKCG, PRKCB | 2 | 0.015598 |
|  | GO:0043209 - Myelin sheath | HSP90AA1, PRKCZ, ERBB2, SRD5A1, BCL2 | 5 | 6.28E-05 |
|  | GO:0097038 - Perinuclear endoplasmic reticulum | PIK3R1, FYN | 2 | 0.020688 |
|  | GO:0000791 - Euchromatin | ESR1, SIRT1, CTNNB1, JUN, JAK2, HIF1A | 6 | 1.81E-05 |
|  | GO:0005901 - Caveola | INSR, SRC, PTGS2, JAK2, MAPK1, MAPK3, IGF1R, CTNNB1 | 8 | 1.04E-06 |
| Molecular functions | GO:0035004 - Phosphatidylinositol 3-kinase activity | PIK3CB, PIK3CG, PIK3CA, PIK3CD, ATM | 5 | 4.31E-08 |
|  | GO:0004707 - MAP kinase activity | MAPK9, MAPK1, MAPK3, MAPK8, MAPK14, MAPK11, MAP3K7 | 7 | 4.96E-11 |
|  | GO:0004698 - Calcium-dependent protein kinase C activity | PRKCQ, PRKCZ, PRKCG, PRKCA, PRKCD, PRKCB, PRKCE | 7 | 7.73E-11 |
|  | GO:0004708 - MAP kinase activity | MAP2K2, MAP2K1, MAPK1, MAPK3, BRAF, MAPK14 | 6 | 6.61E-09 |
|  | GO:0035173 - Histone kinase activity | JAK2, CDK2, PRKCA, PRKCB, CDK1, ATM | 6 | 9.30E-09 |
|  | GO:0016004 - Phospholipase activator activity | FYN, LCK, SRC, CASP3, PDPK1 | 5 | 2.99E-07 |
|  | GO:0009931 - Calcium-dependent protein serine/threonine kinase activity | PRKCQ, PRKCZ, PRKCG, PRKCA, PRKCD, PRKCB, PRKCE | 7 | 7.84E-10 |
|  | GO:0052742 - Phosphatidylinositol kinase activity | PIK3CB, PIK3CG, PIK3CA, PIK3CD, ATM | 5 | 3.87E-07 |
|  | GO:0005131 - Growth hormone receptor binding | JAK2, JAK3, JAK1 | 3 | 0.000162 |
|  | GO:0043274 - Phospholipase binding | PRKCZ, FYN, PDPK1, SYK, LCK, SRC | 6 | 3.04E-08 |
|  | GO:0004715 - Non-membrane spanning protein tyrosine kinase activity | FYN, JAK2, JAK3, ITK, ZAP70, JAK1, SYK, PTK2, LCK, SRC, LYN, PRKCD | 12 | 7.70E-16 |
|  | GO:0060229 - Lipase activator activity | FYN, LCK, SRC, CASP3, PDPK1 | 5 | 6.70E-07 |
|  | GO:0016307 - Phosphatidylinositol phosphate kinase activity | PIK3CB, PIK3CG, PIK3CA, PIK3CD | 4 | 1.19E-05 |
|  | GO:0030283 - Testosterone dehydrogenase [NAD(P)] activity | HSD17B2, SRD5A2, AKR1C1, AKR1C3 | 4 | 1.19E-05 |
|  | GO:0008395 - Steroid hydroxylase activity | CYP2C9, CYP17A1, CYP3A4, CYP11B1, CYP2C19, CYP11B2, CYP2A6, CYP1B1, CYP1A1 | 9 | 1.97E-14 |
|  | GO:0070330 - Aromatase activity | CYP19A1, CYP1B1, CYP2C9, CYP1A1, CYP1A2, CYP3A4, CYP2C19 | 7 | 3.14E-09 |
|  | GO:0001784 - Phosphotyrosine residue binding | MAPK1, MAPK3, PTPN6, ZAP70, GRB2, SYK, PTPN11, VAV1, PIK3R1, LCK | 10 | 1.01E-12 |
|  | GO:0005158 - Insulin receptor binding | IGF1R, PTPN1, SRC, PIK3R1, PTPN11 | 5 | 1.09E-06 |
|  | GO:0005161 - Platelet-derived growth factor receptor binding | LYN, PDGFRB, PDGFRA | 3 | 0.000334 |
|  | GO:0043548 - Phosphatidylinositol 3-kinase binding | JAK2, PDGFRB, IGF1R, PIK3R1, INSR, LCK | 6 | 1.54E-07 |

### KEGG pathway enrichment analysis

The 137 common targets were input into ShinyGO for KEGG pathway enrichment analysis to explore the critical signaling pathways of Vernolac targeting cancer. A total of 217 KEGG pathways were significantly enriched. The top 20 KEGG pathways were selected based on fold enrichment and FDR, and they were primarily related to cancer, immune regulation, endocrine pathways, and metabolic pathways (Table 4). The dot plot of the top 20 KEGG pathways is shown in Fig 10. Notably, these pathways signify Vernolac’s potential role in targeting various types of cancers, such as prostate cancer, glioma, non-small cell lung carcinoma, endometrial cancer, bladder cancer, pancreatic cancer, renal cell carcinoma, as well as hematological malignancies, including acute myeloid leukemia and chronic myeloid leukemia.

**Fig 10.**
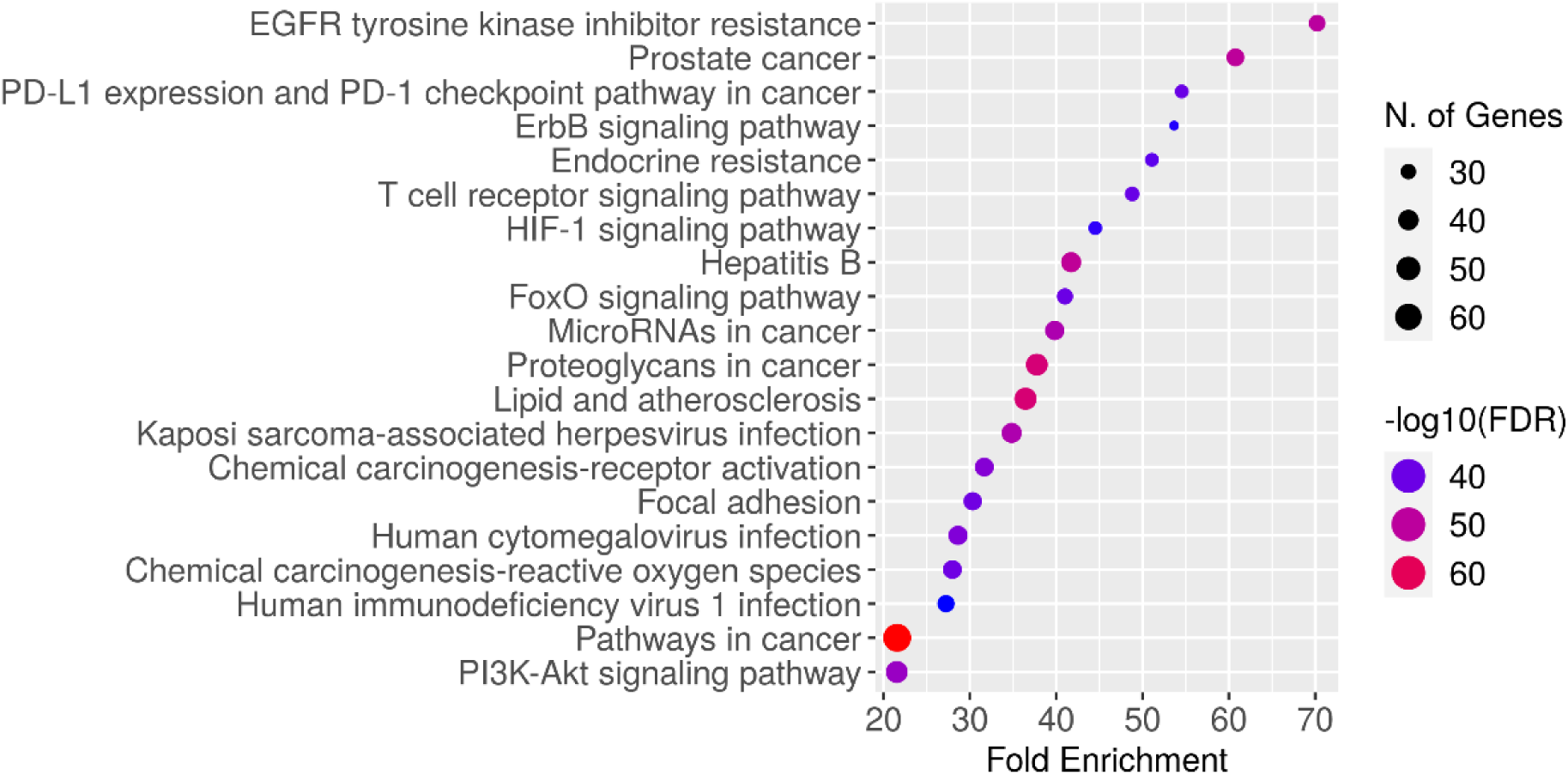
KEGG pathway enrichment analysis. The size of each dot corresponds to the number of genes annotated in the entry. The color of each dot corresponds to the false discovery rate (FDR).

**Table 4.** Top 20 KEGG pathways.

| Pathway | Genes | Count | Enrichment FDR |
| --- | --- | --- | --- |
| Path:hsa01521 - EGFR tyrosine kinase inhibitor resistance | EGFR, ERBB2, AKT1, FGF2, MTOR, GRB2, GSK3B, IGF1R, IL6, JAK1, JAK2, KDR, MET, PDGFRA, PDGFRB, PIK3CA, PIK3CB, PIK3CD, PIK3R1, PRKCA, PRKCB, PRKCG, MAPK1, MAPK3, MAP2K1, MAP2K2, RAF1, BCL2, BCL2L1, SRC, BRAF, STAT3 | 32 | 1.63E-50 |
| Path:hsa05215 - Prostate cancer | CDK2, CHUK, CREBBP, CTNNB1, EGFR, EP300, ERBB2, AKT1, MTOR, GRB2, GSK3B, HSP90AA1, HSP90AB1, IGF1R, IKBKB, AR, MDM2, MMP9, PDGFRA, PDGFRB, PDPK1, PIK3CA, PIK3CB, PIK3CD, PIK3R1, MAPK1, MAPK3, MAP2K1, MAP2K2, RAF1, BCL2, RELA, SRD5A2, BRAF | 34 | 4.34E-51 |
| Path:hsa04370 - VEGF signaling pathway | MAPK14, AKT1, KDR, PIK3CA, PIK3CB, PIK3CD, PIK3R1, PRKCA, PRKCB, PRKCG, MAPK1, MAPK3, MAPK11, MAP2K1, MAP2K2, SPHK2, PTGS2, PTK2, RAF1, SRC | 20 | 4.61E-30 |
| Path:hsa04917 - Prolactin signaling pathway | MAPK14, CYP17A1, AKT1, ESR1, ESR2, GRB2, GSK3B, JAK2, PIK3CA, PIK3CB, PIK3CD, PIK3R1, MAPK1, MAPK3, MAPK8, MAPK11, MAPK9, MAP2K1, MAP2K2, RAF1, RELA, SRC, STAT3 | 23 | 5.76E-34 |
| Path:hsa05235 - PD-L1 expression and PD-1 checkpoint pathway in cancer | CHUK, MAPK14, EGFR, AKT1, MTOR, HIF1A, IKBKB, JAK1, JAK2, JUN, LCK, PIK3CA, PIK3CB, PIK3CD, PIK3R1, PRKCQ, MAPK1, MAPK3, MAPK11, MAP2K1, MAP2K2, PTPN6, PTPN11, RAF1, RELA, STAT3, TLR4, ZAP70 | 28 | 1.22E-40 |
| Path:hsa04012 - ErbB signaling pathway | EGFR, ERBB2, AKT1, MTOR, GRB2, GSK3B, JUN, PIK3CA, PIK3CB, PIK3CD, PIK3R1, PRKCA, PRKCB, PRKCG, MAPK1, MAPK3, MAPK8, MAPK9, MAP2K1, MAP2K2, PTK2, RAF1, SRC, BRAF, CAMK2A, CAMK2B | 26 | 1.49E-37 |
| Path:hsa04664 - Fc epsilon RI signaling pathway | MAPK14, AKT1, FYN, GRB2, LYN, PDPK1, PIK3CA, PIK3CB, PIK3CD, PIK3R1, PRKCA, MAPK1, MAPK3, MAPK8, MAPK11, MAPK9, MAP2K1, MAP2K2, RAF1, SYK, VAV1 | 21 | 1.57E-30 |
| Path:hsa05214 - Glioma | EGFR, AKT1, MTOR, GRB2, IGF1R, MDM2, PDGFRA, PDGFRB, PIK3CA, PIK3CB, PIK3CD, PIK3R1, PRKCA, PRKCB, PRKCG, MAPK1, MAPK3, MAP2K1, MAP2K2, RAF1, BRAF, CAMK2A, CAMK2B | 23 | 3.34E-33 |
| Path:hsa05223 - Non-small cell lung cancer | EGFR, ERBB2, AKT1, GRB2, JAK3, MET, PDPK1, PIK3CA, PIK3CB, PIK3CD, PIK3R1, PRKCA, PRKCB, PRKCG, MAPK1, MAPK3, MAP2K1, MAP2K2, RAF1, RXRA, BRAF, STAT3 | 22 | 8.67E-32 |
| Path:hsa04930 - Type II diabetes mellitus | MTOR, IKBKB, INSR, PIK3CA, PIK3CB, PIK3CD, PIK3R1, PRKCD, PRKCE, PRKCZ, MAPK1, MAPK3, MAPK8, MAPK9 | 14 | 1.15E-20 |
| Path:hsa04960 - Aldosterone-regulated sodium reabsorption | INSR, PDPK1, PIK3CA, PIK3CB, PIK3CD, PIK3R1, PRKCA, PRKCB, PRKCG, MAPK1, MAPK3 | 11 | 2.38E-16 |
| Path:hsa01522 - Endocrine resistance | MAPK14, EGFR, ERBB2, AKT1, ESR1, ESR2, MTOR, GRB2, IGF1R, JUN, MDM2, MMP9, PIK3CA, PIK3CB, PIK3CD, PIK3R1, MAPK1, MAPK3, MAPK8, MAPK11, MAPK9, MAP2K1, MAP2K2, PTK2, RAF1, BCL2, SRC, BRAF | 28 | 9.12E-40 |
| Path:hsa05213 Endometrial cancer | CTNNB1, EGFR, ERBB2, AKT1, GRB2, GSK3B, PDPK1, PIK3CA, PIK3CB, PIK3CD, PIK3R1, MAPK1, MAPK3, MAP2K1, MAP2K2, RAF1, BRAF | 17 | 1.41E-24 |
| Path:hsa05219 Bladder cancer | EGFR, ERBB2, CXCL8, MDM2, MMP9, MAPK1, MAPK3, MAP2K1, MAP2K2, RAF1, SRC, BRAF | 12 | 1.20E-17 |
| Path:hsa04660 T cell receptor signaling pathway | CHUK, MAPK14, AKT1, FYN, GRB2, GSK3B, IKBKB, ITK, JUN, LCK, PDPK1, PIK3CA, PIK3CB, PIK3CD, PIK3R1, PRKCQ, MAPK1, MAPK3, | 29 | 1.56E-40 |
|  | MAPK8, MAPK11, MAPK9, MAP2K1, MAP2K2, PTPN6, RAF1, RELA, MAP3K7, VAV1, ZAP70 |  |  |
| Path:hsa05212<br>Pancreatic cancer | CHUK, EGFR, ERBB2, AKT1, MTOR, IKBKB, JAK1, PIK3CA, PIK3CB, PIK3CD, PIK3R1, MAPK1, MAPK3, MAPK8, MAPK9, MAP2K1, RAF1, RELA, BCL2L1, BRAF, STAT3 | 21 | 2.09E-29 |
| Path:hsa05221<br>Acute myeloid leukemia | CHUK, AKT1, MTOR, GRB2, IKBKB, KIT, PIK3CA, PIK3CB, PIK3CD, PIK3R1, MAPK1, MAPK3, MAP2K1, MAP2K2, RAF1, RELA, BRAF, STAT3 | 18 | 3.65E-25 |
| Path:hsa05211<br>Renal cell carcinoma | CREBBP, EP300, AKT1, GRB2, HIF1A, JUN, MET, PIK3CA, PIK3CB, PIK3CD, PIK3R1, MAPK1, MAPK3, MAP2K1, MAP2K2, PTPN11, RAF1, BRAF | 18 | 4.87E-25 |
| Path:hsa05220 -<br>Chronic myeloid leukemia | CHUK, AKT1, GRB2, HDAC1, HDAC2, IKBKB, MDM2, PIK3CA, PIK3CB, PIK3CD, PIK3R1, MAPK1, MAPK3, MAP2K1, MAP2K2, PTPN11, RAF1, RELA, BCL2L1, BRAF | 20 | 1.30E-27 |
| Path:hsa04662 - B<br>cell receptor signaling pathway | CHUK, AKT1, GRB2, GSK3B, IKBKB, JUN, LYN, PIK3CA, PIK3CB, PIK3CD, PIK3R1, PRKCB, MAPK1, MAPK3, MAP2K1, MAP2K2, PTPN6, RAF1, RELA, SYK, VAV1 | 21 | 8.81E-29 |

### Target-pathway network construction

A network was constructed to visualize interactions between the top 20 KEGG pathways and targets (Fig 11). The target-pathway network consisted of 242 nodes and 6804 edges, illustrating the complex interconnectivity between common targets and KEGG pathways. According to the results of network analysis, the top 10 high-degree targets were identified, including AKT1, EGFR, STAT3, CTNNB1, GAPDH, ESR1, TNF, JUN, SRC, and BCL2. The highest degree pathway, representing the most interconnected KEGG pathway, was prostate cancer (hsa05215).

**Fig 11.**
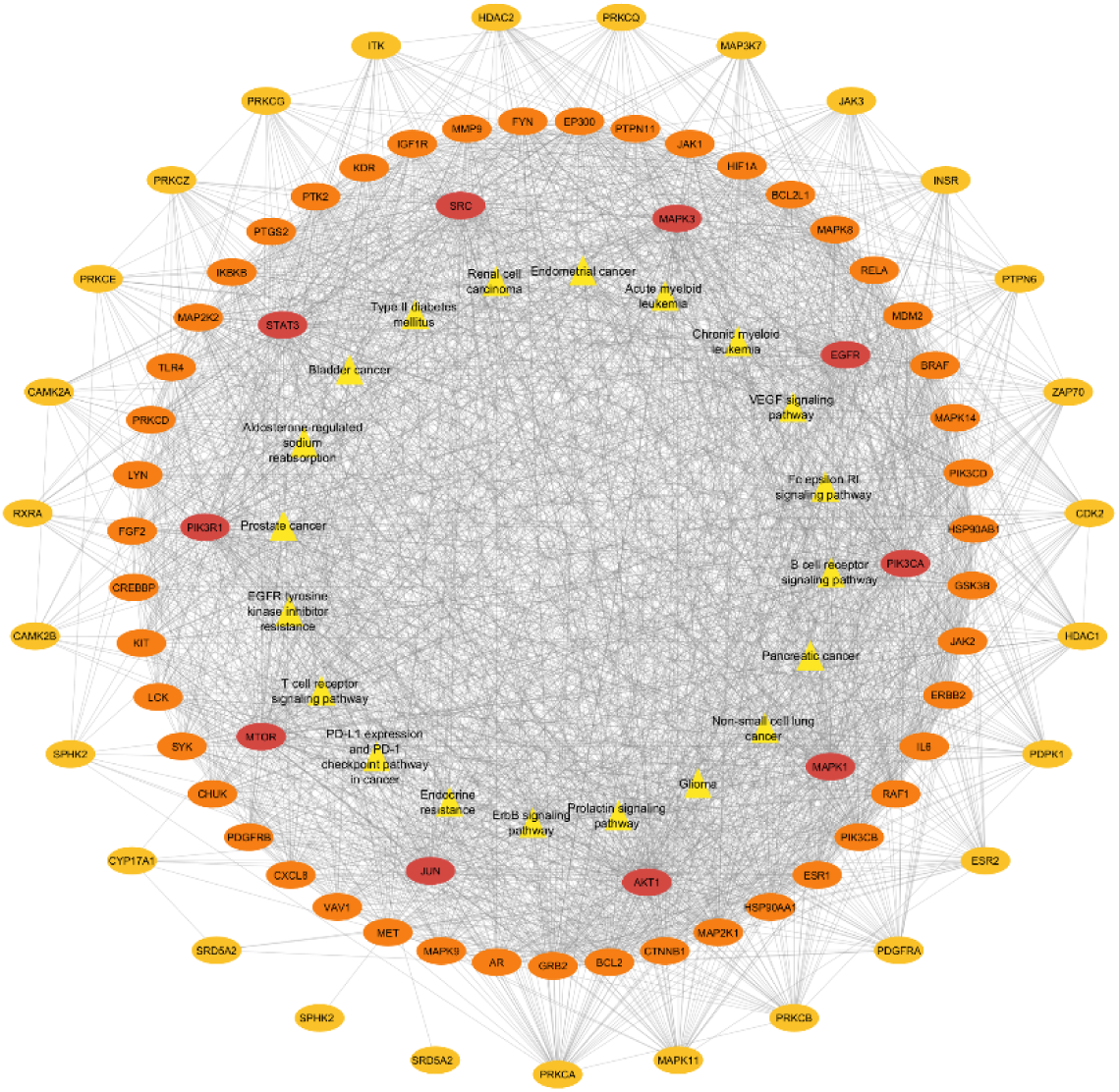
Target-pathway network. Yellow rectangles represent significantly enriched pathways, while elliptical nodes represent target genes. The ellipses are colored from red (innermost, highest degree) to orange and yellow (outermost, lower degree), indicating the decreasing node degree (connectivity) of the targets. Edges represent interactions between target-target and target-pathways. The arrangement highlights central targets with greater involvement across multiple pathways, emphasizing their potential as key regulators in Vernolac’s anticancer activity.

### Molecular docking studies of key target proteins

The ten potential target proteins with high node scores and confidence, including AKT1, CTNNB1, PIK3CA, STAT3, MAPK3, CDK4, CDK6, JAK1, JAK2, and SRC, were selected through network and pathway analysis. Molecular docking studies were conducted using Autodock Vina in PyRx to identify the interactions between the compounds and cancer-related potential target proteins at the molecular level. Table 5 presents the compounds that achieved the highest docking scores with their respective target proteins. The lower the docking score, the higher the binding affinity between the compound and the target protein.

**Table 5.**
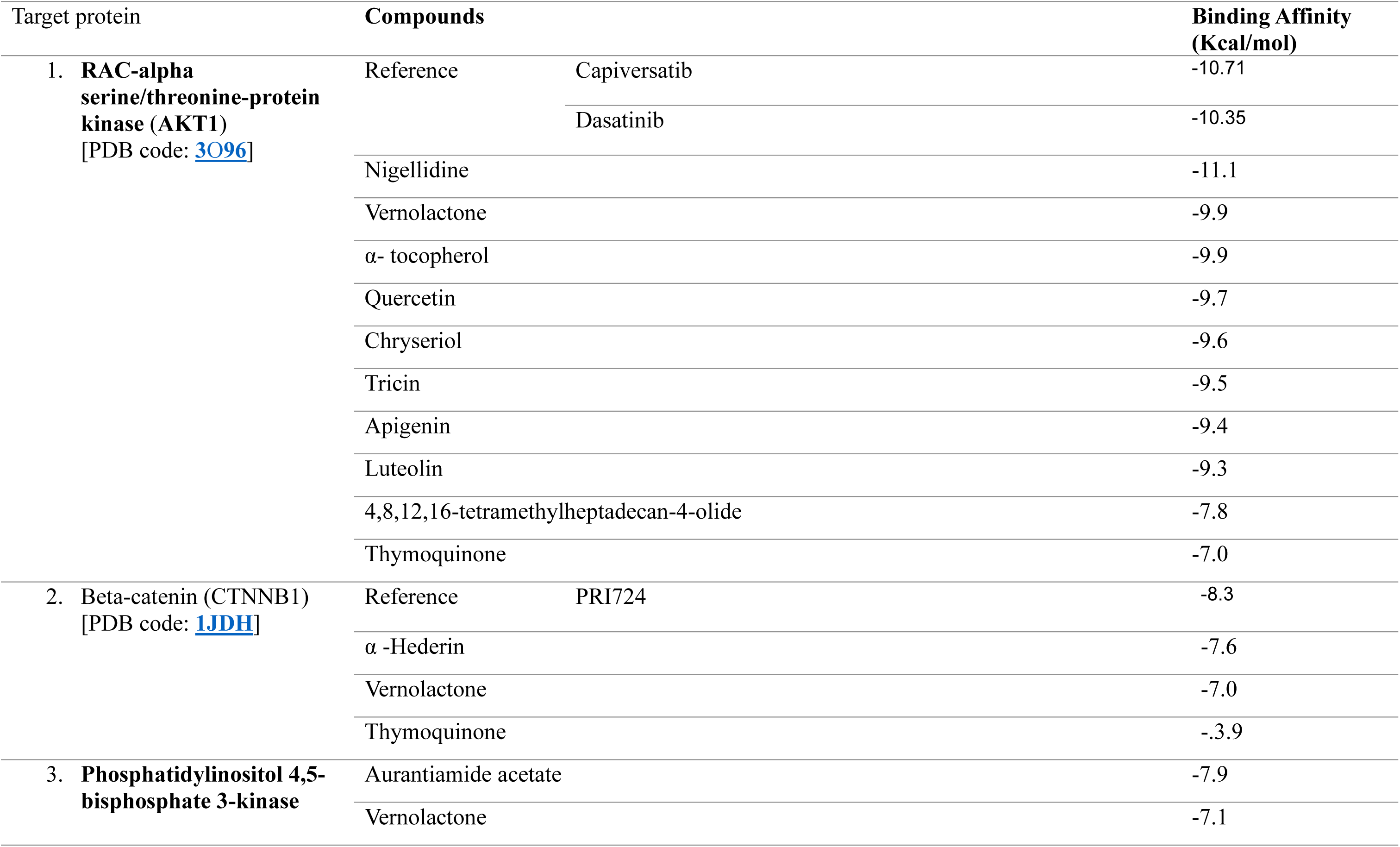
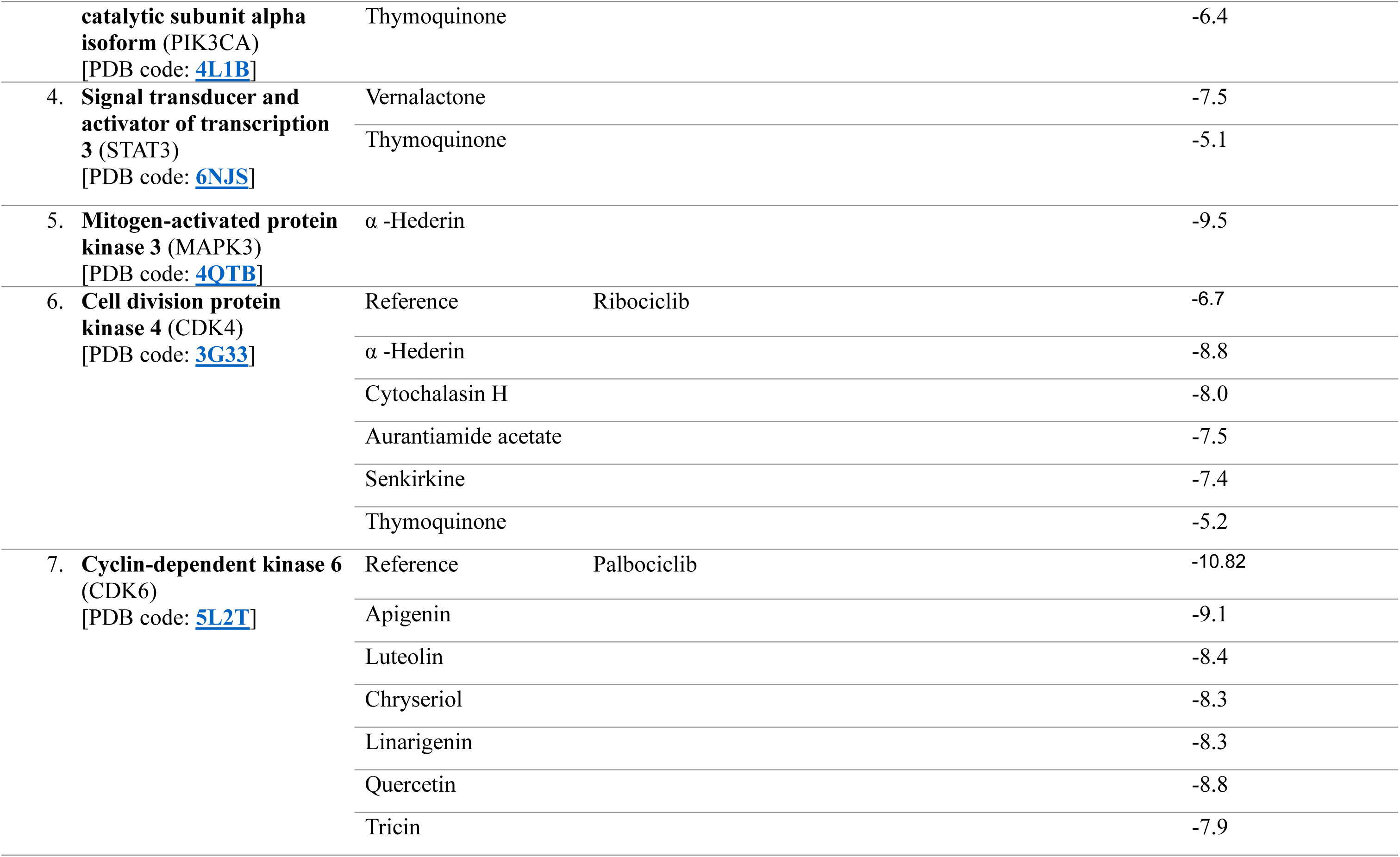
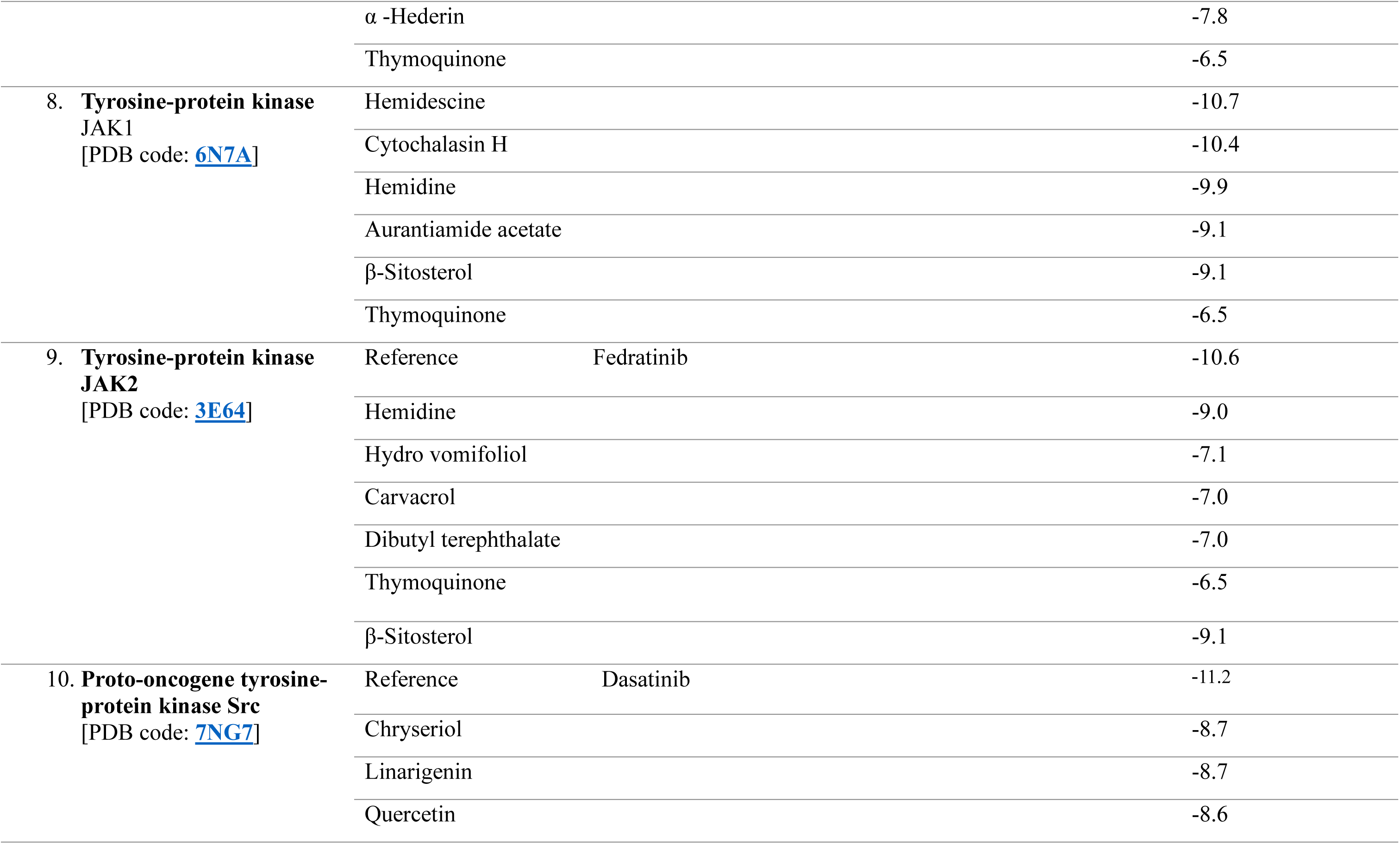
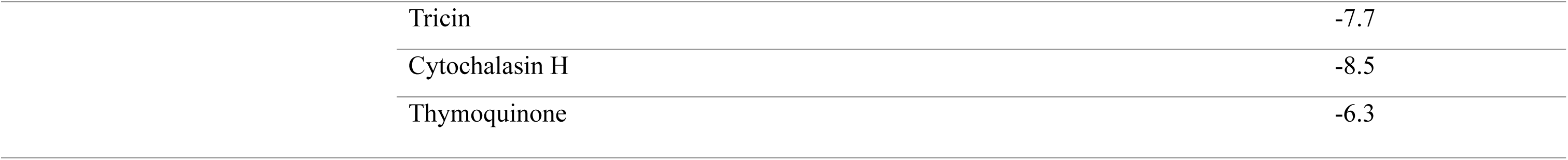
Docking scores of targets with the corresponding compounds.

**Table 6.** Summary of the mean Root Mean Square Deviation (RMSD) for natural compounds docked to the AKT1, Beta-catenin, CDK4, CDK6, JAK1, JAK2, MAPK3, STAT3, PIK3CA, SRC protein.

| Target protein | Compound | Mean RMSD (nm) |
| --- | --- | --- |
| AKT1 | Quercetin | 0.17 ± 0.05 |
|  | Dasatinib (reference) | 0.21 ± 0.04 |
|  | Vernolactone | 0.21 ± 0.05 |
|  | Capivasertib (reference) | 0.24 ± 0.03 |
|  | Tricin | 0.25 ± 0.13 |
|  | Chryseriol | 0.33 ± 0.10 |
|  | 4,8,12,16-Tetramethylheptadecan-4-olide | 0.35 ± 0.05 |
|  | Luteolin | 0.38 ± 0.07 |
|  | Thymoquinone | 0.43 ± 0.20 |
|  | Alpha tocopherol | 0.47 ± 0.11 |
|  | Nigellidine | 0.50 ± 0.05 |
|  | Apigenin | 0.53 ± 0.18 |
| **CTNNB1** | PRI724 (reference) | $0.86 \pm 0.17$ |
| | Vernolactone | $0.88 \pm 0.13$ |
| | alpha-Hederin | $0.33 \pm 0.07$ |
| | Thymoquinone | $5.08 \pm 2.29$ |
| **CDK4** | Ribociclib (reference) | $1.02 \pm 0.17$ |
| | alpha-Hederin | $0.46 \pm 0.10$ |
| | Cytochalasin H | $0.29 \pm 0.07$ |
| | Senkirkine | $0.57 \pm 0.12$ |
| | Thymoquinone | $1.36 \pm 1.41$ |
| | Aurantiamide Acetate | $1.18 \pm 0.27$ |
| **CDK6** | Palcociclib (reference) | $0.48 \pm 0.05$ |
| | Chryseriol | $0.13 \pm 0.03$ |
| | Quercetin | $0.31 \pm 0.05$ |
| | Apigenin | $0.20 \pm 0.03$ |
| | alpha-Hederin | $1.08 \pm 0.29$ |
| | Tricin | $0.49 \pm 0.07$ |
| | Thymoquinone | $2.49 \pm 1.29$ |
| | Luteolin | $1.15 \pm 0.62$ |
| | Linarigenin | $0.30 \pm 0.07$ |
| **JAK1** | Cytochalasin H | $10.26 \pm 0.07$ |
| | Aurantiamide acetate | $10.69 \pm 0.16$ |
| | Hemidescine | $10.92 \pm 0.14$ |
| | β-sitosterol | $10.81 \pm 0.27$ |
| | Hemidine | $10.19 \pm 0.11$ |
| | Thymoquinone | $10.10 \pm 1.48$ |
| **JAK2** | Fedratinib (reference) | $0.28 \pm 0.06$ |
| | Carvacrol | $0.42 \pm 0.06$ |
| | Dibutyl terephthalate | $0.43 \pm 0.15$ |
| | Thymoquinone | $0.38 \pm 0.04$ |
| | Hemidine | $0.48 \pm 0.10$ |
| | Hydro vomifoliol | $0.55 \pm 0.12$ |
| | β-sitosterol | $0.58 \pm 0.11$ |
| **MAPK3** | α-Hederin | $1.34 \pm 0.40$ |
| | Vernolactone | $0.48 \pm 0.07$ |
| | Thymoquinone | $2.99 \pm 1.75$ |
| **STAT3** | Vernolactone | $0.34 \pm 0.08$ |
| | Thymoquinone | $7.69 \pm 3.54$ |
| **PIK3CA** | Vernolactone | $0.40 \pm 0.07$ |
| | Aurantiamide acetate | $0.50 \pm 0.16$ |
| | Thymoquinone | $0.79 \pm 1.84$ |
| SRC | Dasatinib (reference) | $0.28 \pm 0.06$ |
| | Linarigenin | $0.80 \pm 0.11$ |
| | Chryseriol | $0.37 \pm 0.07$ |
| | Cytochalasin H | $0.74 \pm 0.12$ |
| | Quercetin | $0.32 \pm 0.09$ |
| | Tricin | $0.36 \pm 0.19$ |
| | Thymoquinone | $1.11 \pm 0.87$ |

**Table 7.** Summary of the mean Root Mean Square Fluctuation (RMSF) for natural compounds docked to the AKT1, Beta-catenin, CDK4, CDK6, JAK1, JAK2, MAPK3, STAT3, PIK3CA, and SRC proteins.

| Target protein | Compound | Mean RMSF (nm) |
| --- | --- | --- |
| AKT1 | Thymoquinone | 0.15 ± 0.11 |
|  | Capivasertib | 0.15 ± 0.15 |
|  | Vernolac | 0.15 ± 0.12 |
|  | Dasatinib | 0.16 ± 0.14 |
|  | Luteoline | 0.16 ± 0.16 |
|  | Nigellidine | 0.16 ± 0.11 |
|  | Apigenin | 0.16 ± 0.17 |
|  | Quercetin | 0.16 ± 0.16 |
|  | α-tocopherol | 0.16 ± 0.16 |
|  | Tetramethylheptadecan-4-olide | 0.17 ± 0.15 |
|  | Chryseriol | 0.17 ± 0.16 |
|  | Tricin | 0.17 ± 0.15 |
| CTNNB1 | PRI724 | 0.15 ± 0.09 |
| | Thymoquinone | $0.16 \pm 0.09$ |
| | α-hederin | $0.16 \pm 0.07$ |
| | Vernolactone | $0.18 \pm 0.14$ |
| **CDK4** | Aurantiamide acetate | $0.11 \pm 0.07$ |
| | Ribociclib | $0.12 \pm 0.07$ |
| | Cytochalasin-H | $0.12 \pm 0.07$ |
| | α-hederin | $0.12 \pm 0.07$ |
| | Senkirkine | $0.12 \pm 0.09$ |
| | Thymoquinone | $0.14 \pm 0.14$ |
| **CDK6** | Chrysoeriole | $0.11 \pm 0.08$ |
| | Linarigenin | $0.11 \pm 0.12$ |
| | Palbociclib | $0.12 \pm 0.11$ |
| | Thymoquinone | $0.12 \pm 0.09$ |
| | α-hederin | $0.13 \pm 0.14$ |
| | Quercetin | $0.13 \pm 0.12$ |
| | Apigenin | $0.13 \pm 0.14$ |
| | Tricin | $0.14 \pm 0.13$ |
| | Luteolin | $0.15 \pm 0.13$ |
| **JAK1** | Aurantiamide acetate | $0.14 \pm 0.11$ |
| | Thymoquinone | $0.14 \pm 0.14$ |
| | Cytochalasin-H | $0.16 \pm 0.19$ |
| | Hemidesine | $0.18 \pm 0.26$ |
| | β- sitosterol | $0.20 \pm 0.26$ |
| **JAK2** | Dibutyl terephthalate | $0.09 \pm 0.05$ |
| | Hemidine | $0.09 \pm 0.06$ |
| | Hydro vomifoliol | $0.09 \pm 0.06$ |
| | Fedratinib | $0.09 \pm 0.06$ |
| | Thymoquinone | $0.10 \pm 0.06$ |
| | Carvacrol | $0.10 \pm 0.06$ |
| | β- sitosterol | $0.10 \pm 0.06$ |
| **MAPK3** | Vernolactone | $0.15 \pm 0.16$ |
| | α-hederin | $0.160 \pm 0.14$ |
| | Thymoquinone | $0.19 \pm 0.19$ |
| **PIK3CA** | Thymoquinone | $0.12 \pm 0.07$ |
| | Aurantiamide acetate | $0.15 \pm 0.10$ |
| | Vernolactone | $0.15 \pm 0.12$ |
| **SRC** | Chrysoeriol | $0.11 \pm 0.10$ |
| | Quercetin | $0.17 \pm 0.09$ |
| | Dasatinib | $0.12 \pm 0.10$ |
| | Cytochalasin-H | $0.13 \pm 0.13$ |
| | Thymoquinone | $0.13 \pm 0.13$ |
| | Linarigenin | $0.15 \pm 0.16$ |
| | Tricin | $0.16 \pm 0.21$ |

Based on the docking scores presented in the table, several key factors stand out regarding the binding affinity of the compounds. While reference compounds such as Palbociclib for CDK6, Fedratinib for JAK2, and Dasatinib for SRC displayed a superior binding affinity compared to the tested compounds, the results also highlight the strong potential of several compounds. For instance, Nigellidine and a group of compounds including alpha-Hederin, Cytochalasin H, Aurantiamide acetate, and Senkirkine demonstrated a superior binding affinity to the reference compounds for AKT1 and CDK4, respectively.

Importantly, a large number of the tested compounds exhibited excellent docking scores, with most compounds achieving a binding affinity of -7 kcal/mol or better. This indicates a strong potential for binding to the target proteins. The compound with the highest overall binding affinity was Nigellidine at -11.1 kcal/mol for AKT1. Other top performers among the compounds included Hemidescine (-10.7 kcal/mol) for JAK1, alpha-Hederin (-9.5 kcal/mol) for MAPK3, and Apigenin (-9.1 kcal/mol) for CDK6. These results collectively suggest that while some reference controls have the strongest binding, the tested compounds still show promising docking scores and significant potential as inhibitors for the target proteins.

S3 table presents the bond analysis of the docked compounds in relation to their respective target proteins. For further investigations utilizing molecular dynamic simulations, evaluation criteria encompassed not only docking scores but also the binding interaction within the active site pocket and bond analysis to ensure a comprehensive assessment of molecular stability and interaction strength.

The reference compound Capiversatib primarily relies on hydrogen bonds to interact with AKT1, engaging residues such as THR: A:82, VAL: A:271, TYR: A:272, and ARG: A:273. This indicates a highly specific, polar-driven binding. In contrast, natural compounds such as Nigellidine, Thymoquinone, and Apigenin demonstrate a greater dependence on hydrophobic interactions with a similar group of residues, including TRP: A:80, LEU: A:264, and VAL: A:270. This suggests that while they may occupy a similar binding pocket, their mode of interaction is fundamentally different.

PRI724, the reference compound for CTNNB1, utilizes a diverse set of interactions, including hydrogen bonds, a hydrophobic interaction with VAL: A:564, and a unique electrostatic interaction with ARG: A:469. This broad approach allows it to anchor firmly to the protein. The natural compounds, such as α-Hederin and Vernolactone, exhibit a more focused binding profile. Alpha-Hederin and Thymoquinone mainly form hydrogen bonds, while Vernolactone uses a combination of hydrogen bonds and hydrophobic interactions. The reference compound Ribociclib primarily uses hydrophobic interactions with a broad array of residues such as ILE: A:17, VAL: A:25, and LEU: A:152, complemented by a single hydrogen bond with ASP: A:163. The natural compounds, including α-Hederin, Cytochalasin H, and Asperglaucide, mirror this hydrophobic-dominant strategy. They engage in a similar mix of hydrogen bonds and hydrophobic contacts, indicating that they bind to a comparable hydrophobic pocket on the CDK4 protein.

Fedratinib, the reference compound for JAK2, binds through a combination of hydrogen bonds and extensive hydrophobic interactions. The natural compounds like Hemidine, Carvacrol, and Dibutyl terephthalate primarily rely on hydrophobic interactions with a core set of residues, including LEU: A:855, VAL: A:863, and LEU: A:983. This indicates that they effectively occupy the same hydrophobic pocket as Fedratinib, suggesting a similar mechanism of action.

The reference compound Dasatinib uses a mixed-mode binding strategy, with hydrogen bonds, hydrophobic interactions, and a few unique electrostatic interactions. The natural compounds, including Chryseriol, Linarigenin, and Quercetin, also employ a blend of hydrogen bonds and hydrophobic interactions. Notably, they often interact with key residues such as MET: A:344 and THR: A:341, which are also crucial for Dasatinib’s binding. This overlap suggests that these natural compounds bind to the same active site and share a common binding pattern with Dasatinib. Additionally, compounds with previously reported anticancer activities were considered for further investigations.

**Fig 12-1.**
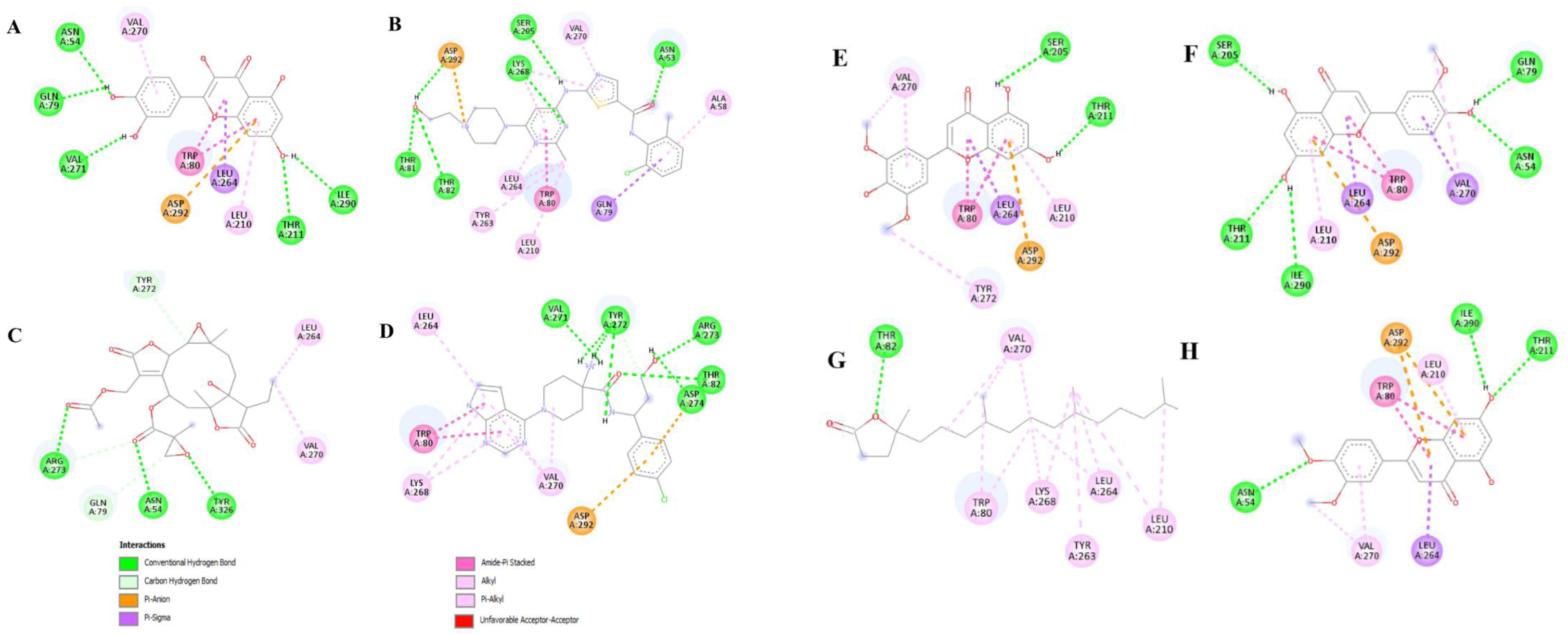
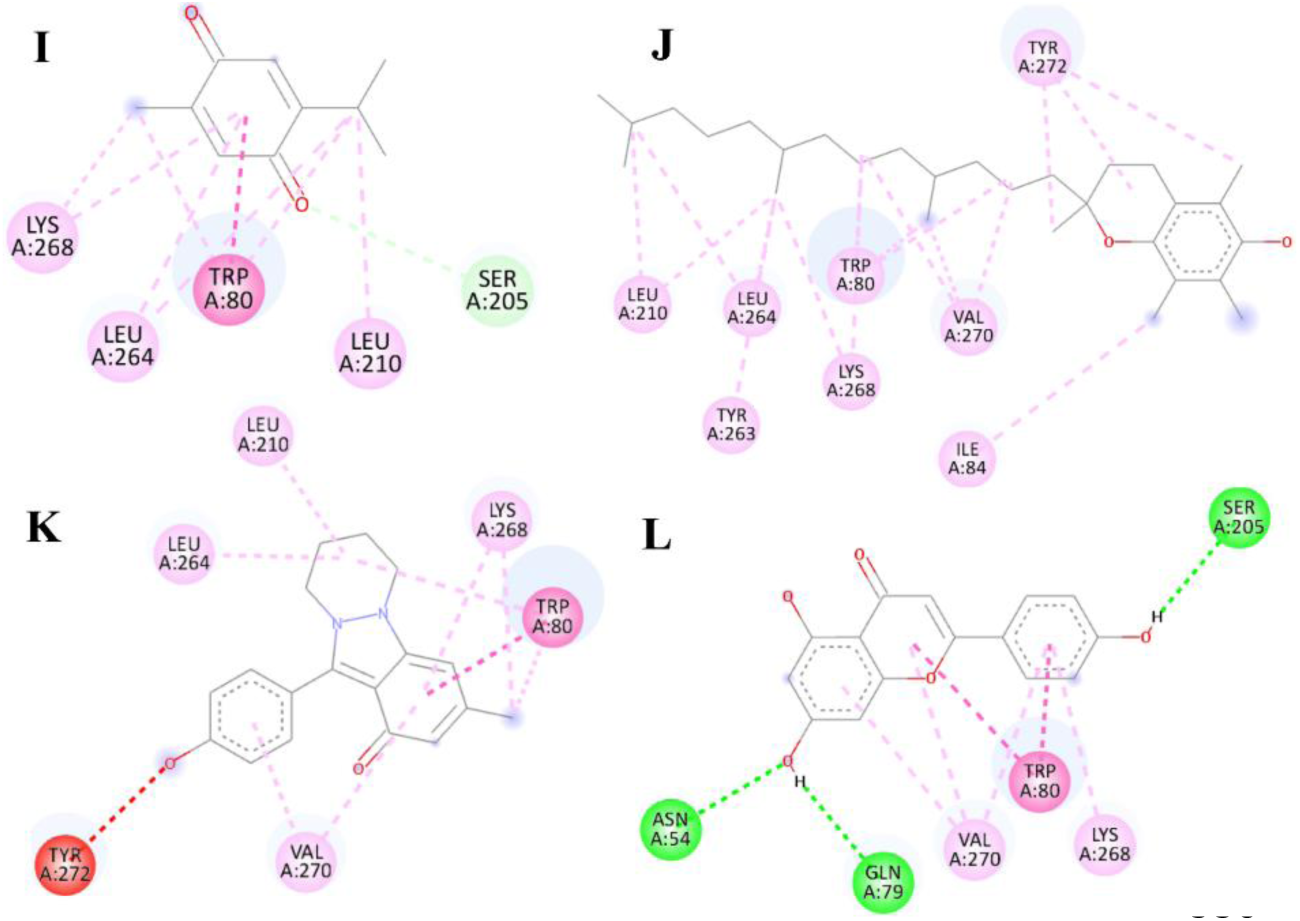
2D protein-ligand interaction diagram of AKT1 protein with compounds; A: Quercetin, B: Dasatinib (reference), C: Vernolactone, D: Capivasertib (reference), E: Tricin, F: Chryseriol, G:4,8,12,16-Tetramethylheptadecan-4-olide, H: Luteolin, I: Thymoquinone, J: Alpha tocopherol, K: Nigellidine and L: Apigenin

**Fig 12-2.**
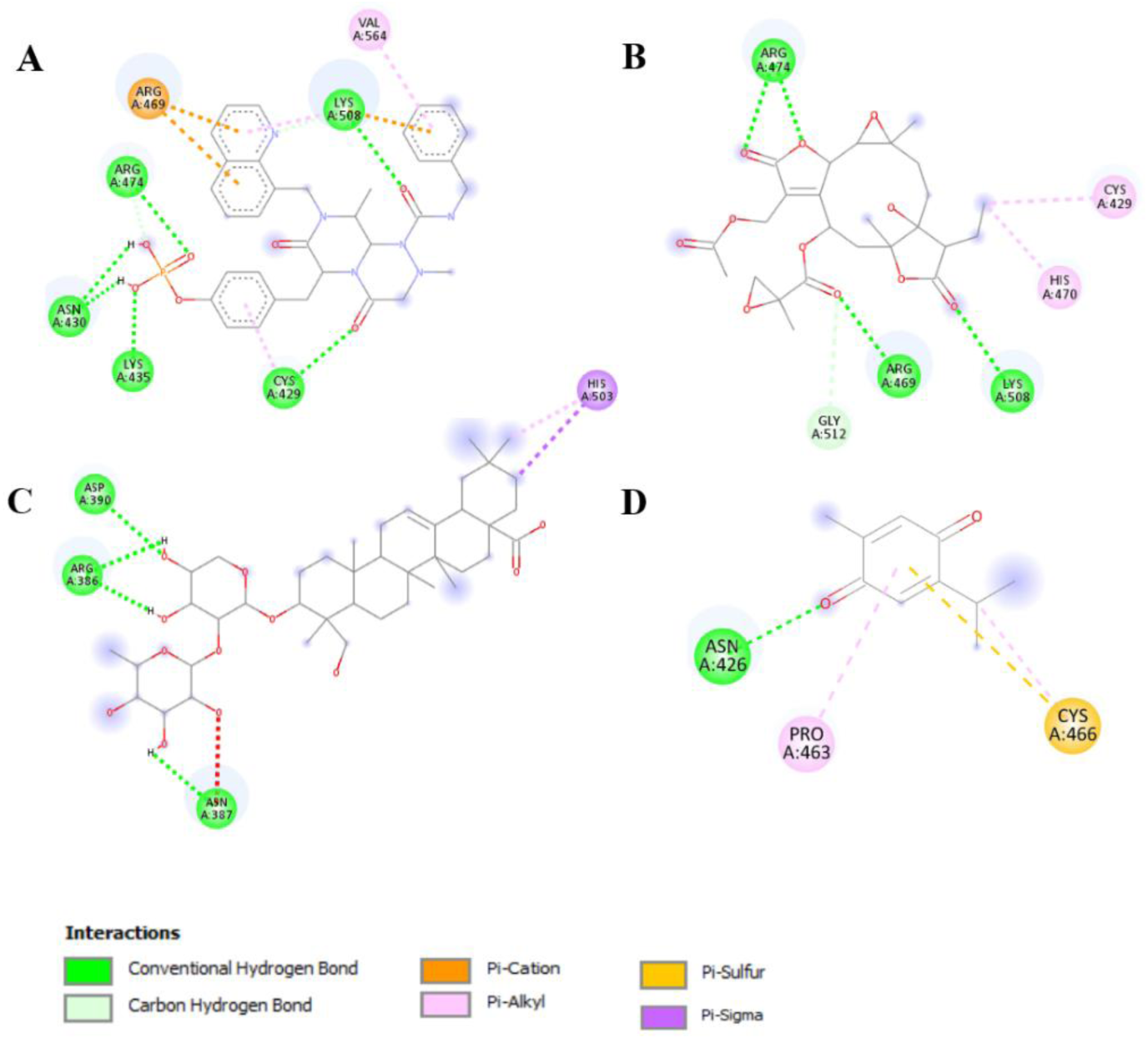
2D protein-ligand interaction diagram of CTNNB1 protein with compounds; A: PRI724 (reference), B: Vernolactone, C: α-Hederin and D: Thymoquinone

**Fig. 12-3.**
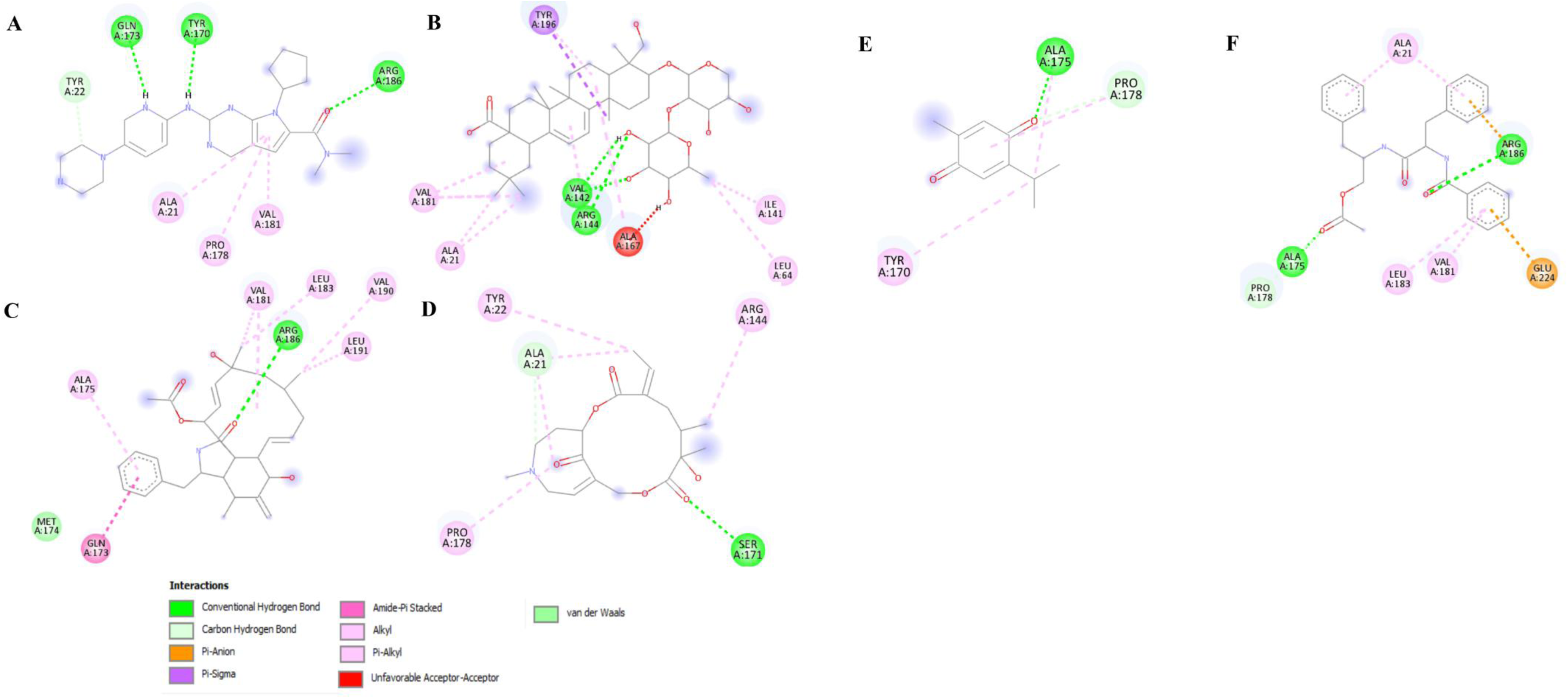
2D protein-ligand interaction diagram of CDK4 protein with compounds; A: Ribociclib (reference), B: α-Hederin, C: Cytochalasin H, D: Senkirkine, E: Thymoquinone and F: Aurantiamide Acetate

**Fig. 12-4.**
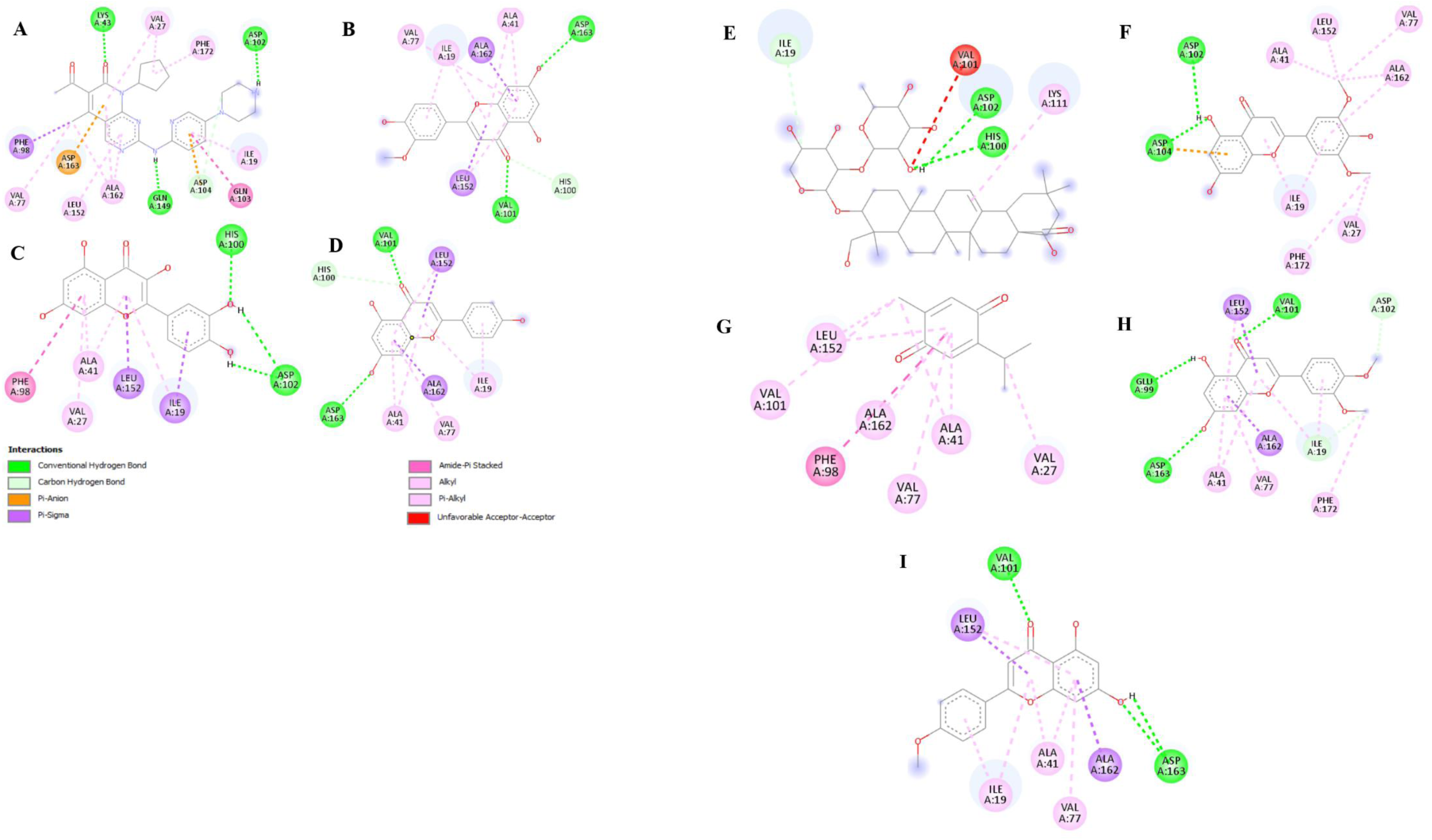
2D protein-ligand interaction diagram of CDK6 protein with compounds; A: Palcociclib (reference), B: Chryseriol, C: Quercetin, D: Apigenin, E: α-Hederin, F: Tricin, G: Thymoquinone, H: Luteolin and I: Linarigenin

**Fig 12-5.**
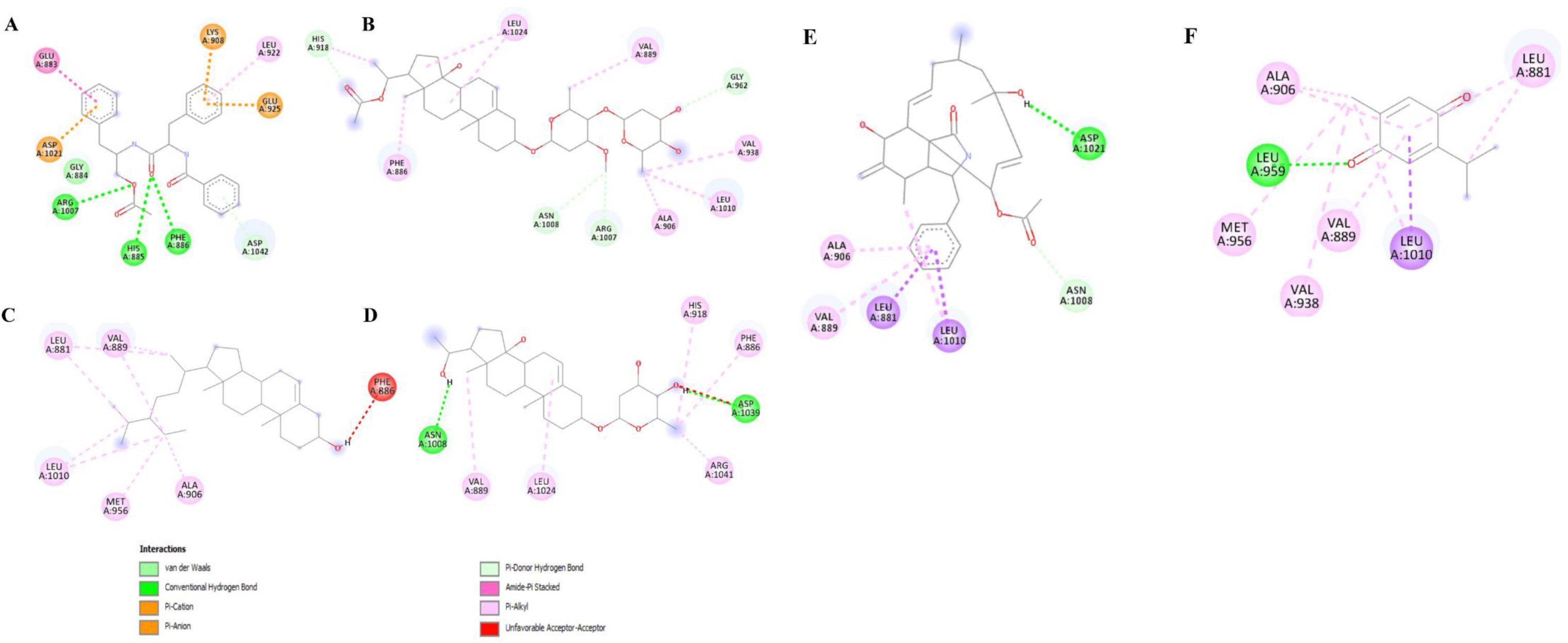
2D protein-ligand interaction diagram of JAK1 protein with compounds; A: Aurantiamide acetate, B: Hemidescine, C: β-sitosterol, D: Hemidine, E: Cytochalasin H and F: Thymoquinone

**Fig 12-6.**
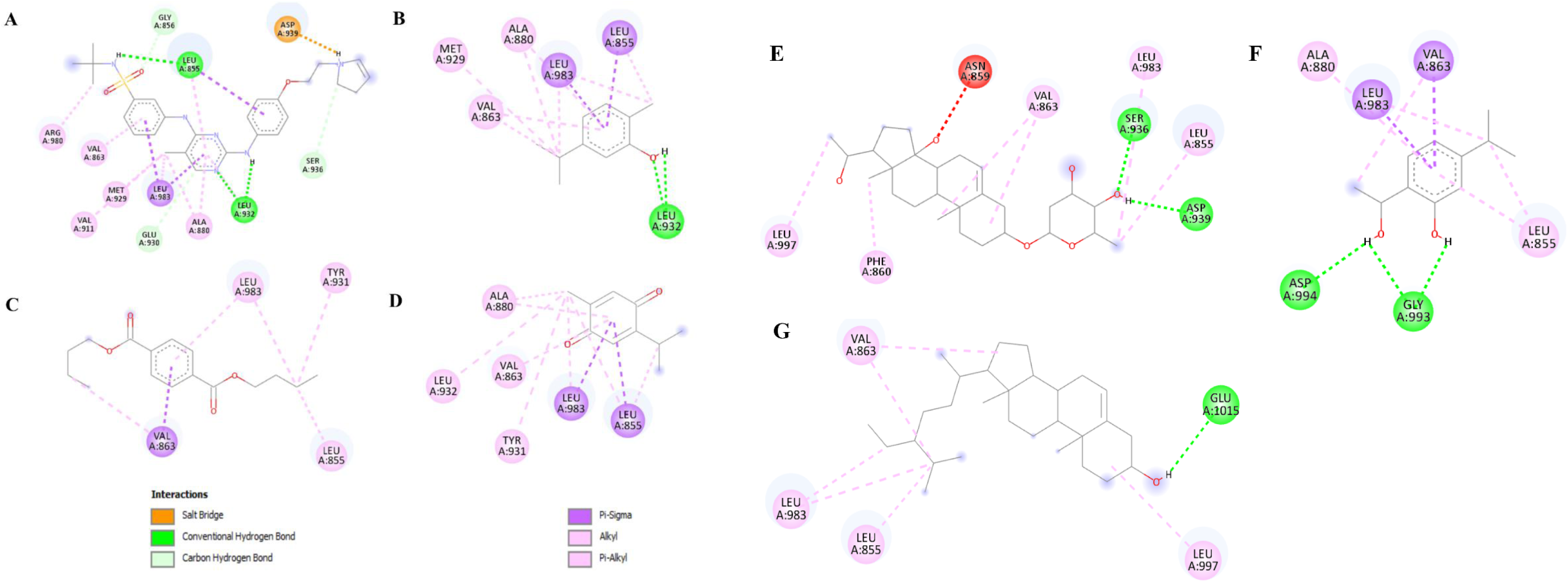
2D protein-ligand interaction diagram of JAK2 protein with compounds; A: Fedratinib (reference), B: Carvacrol, C: Dibutyl terephthalate, D: Thymoquinone, E: Hemidine, F: Hydro vomifoliol and G: β-sitosterol

**Fig 12-7.**
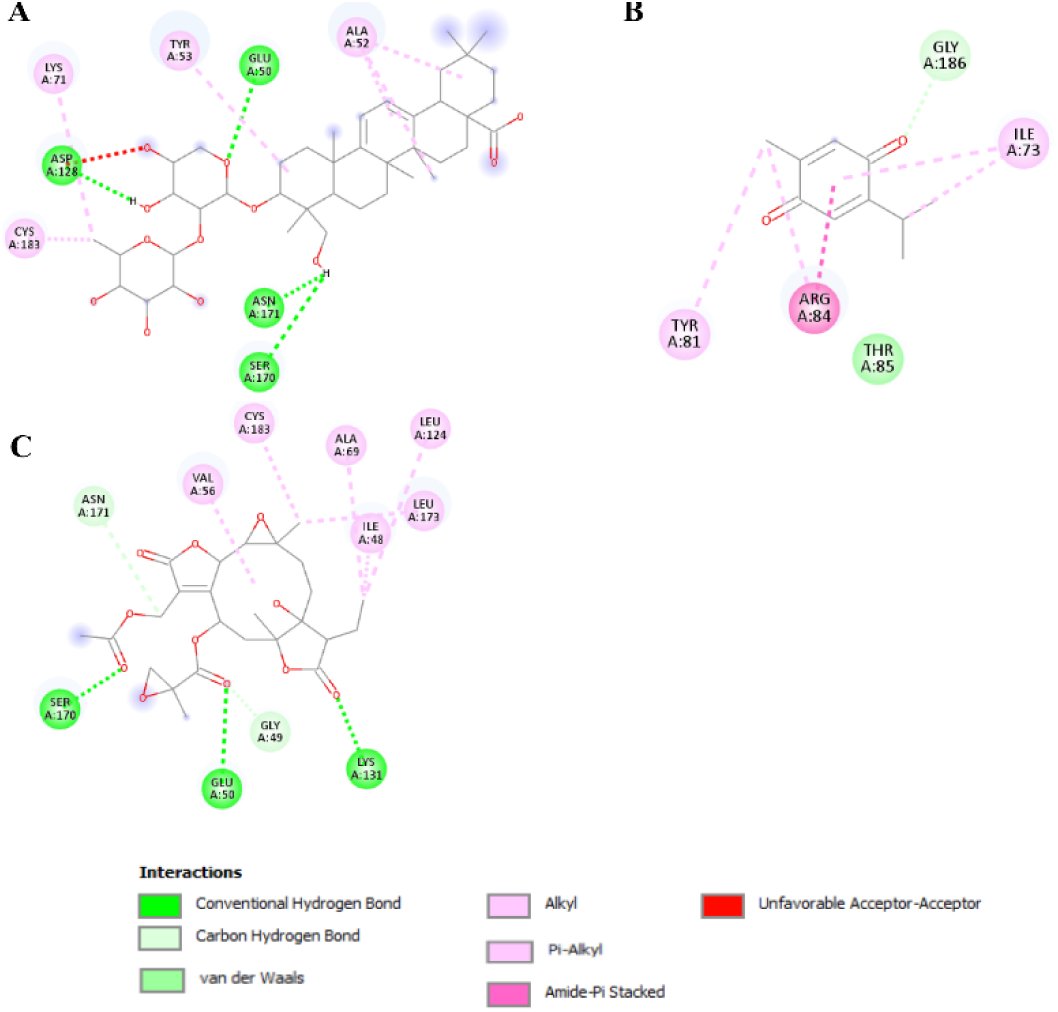
2D protein-ligand interaction diagram of MAPK3 protein with compounds: A: α-Hederin, B: Thymoquinone and C: Vernolactone

**Fig 12-8.**
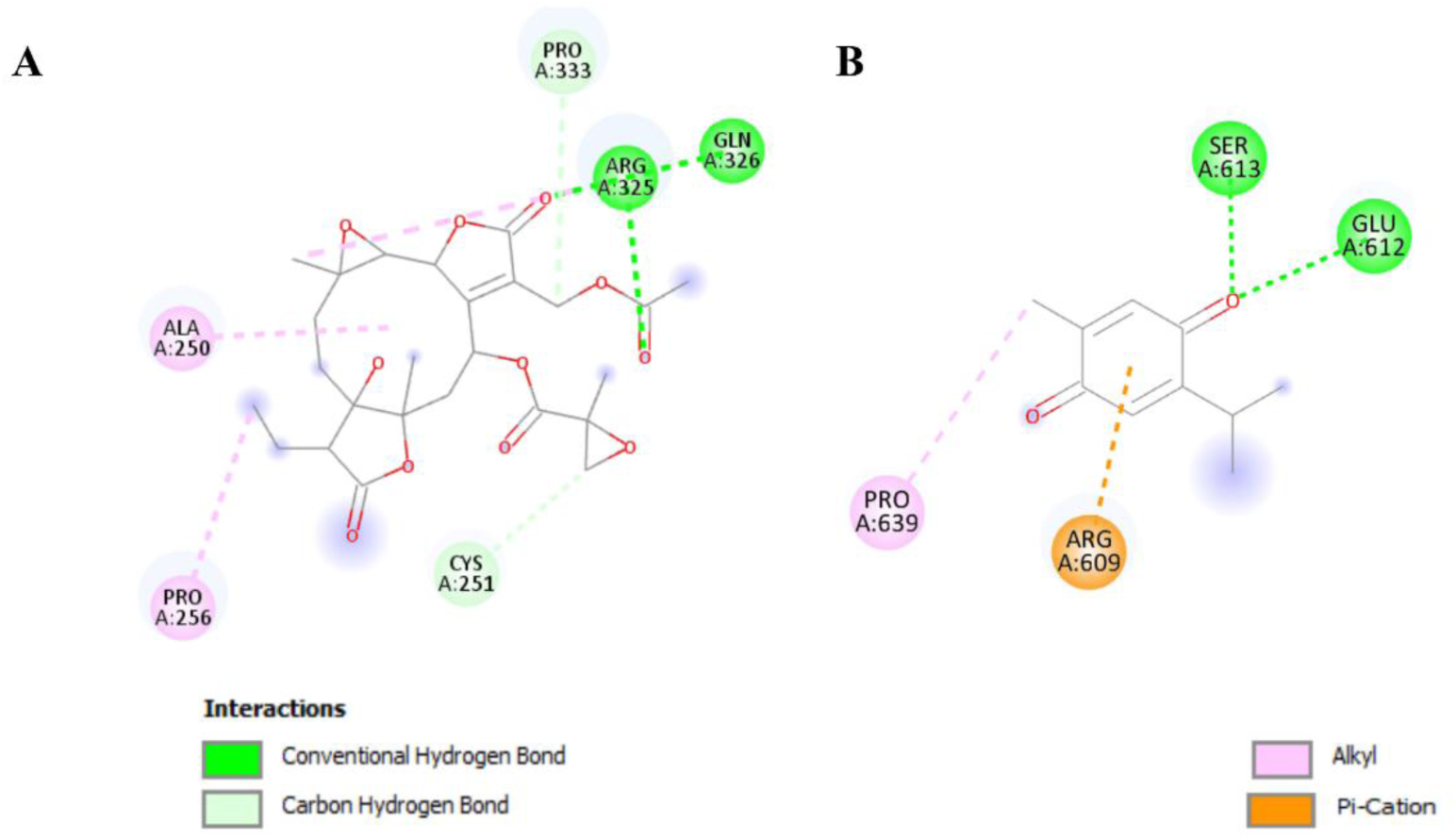
2D protein-ligand interaction diagram of STAT3 protein with compounds; A: Vernolactone and B: Thymoquinone

**Fig 12-9.**
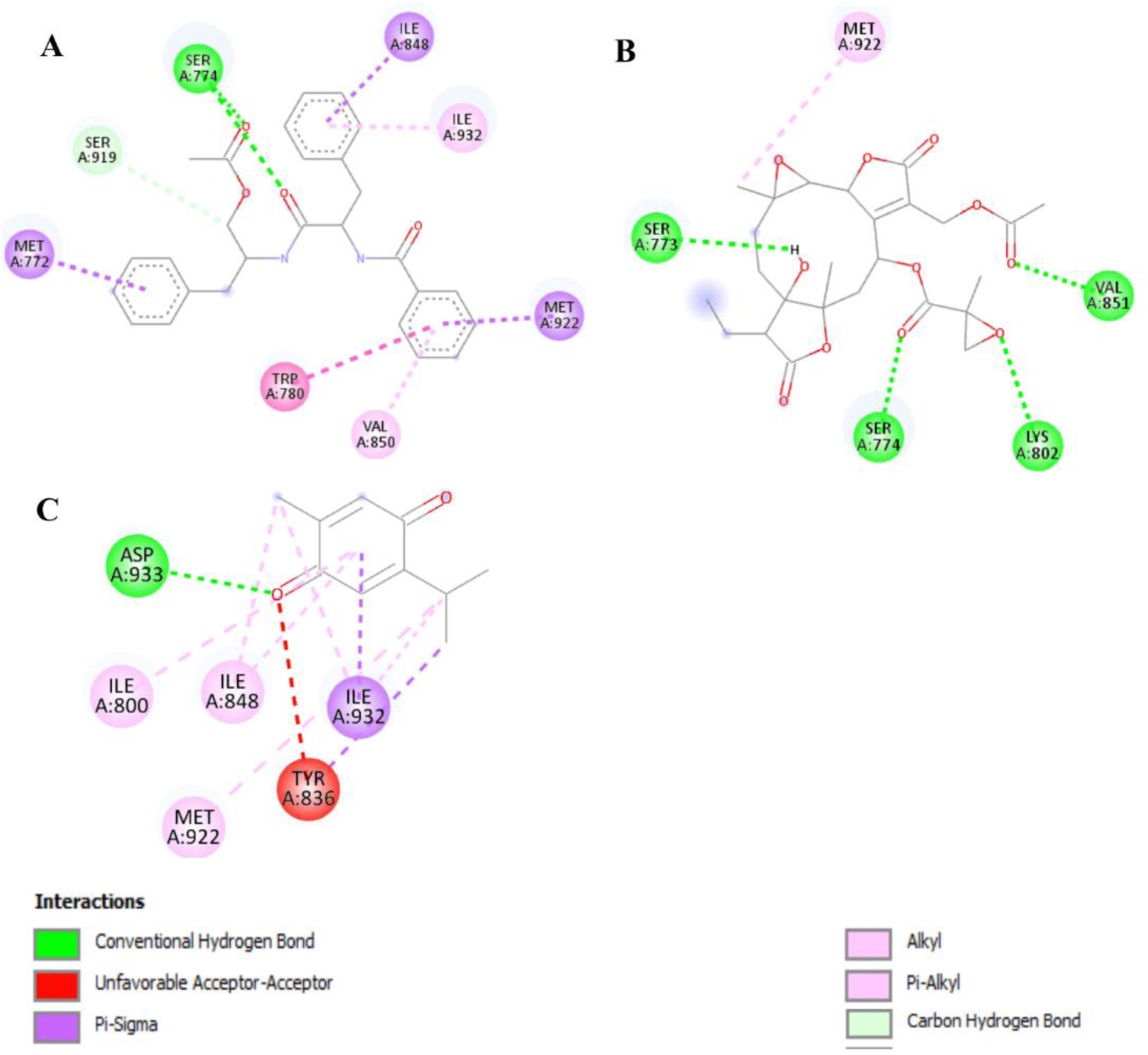
2D protein-ligand interaction diagram of PIK3CA protein with compounds; A: Vernolactone, B: Aurantiamide acetate and C: Thymoquinone

**Fig. 12-10.**
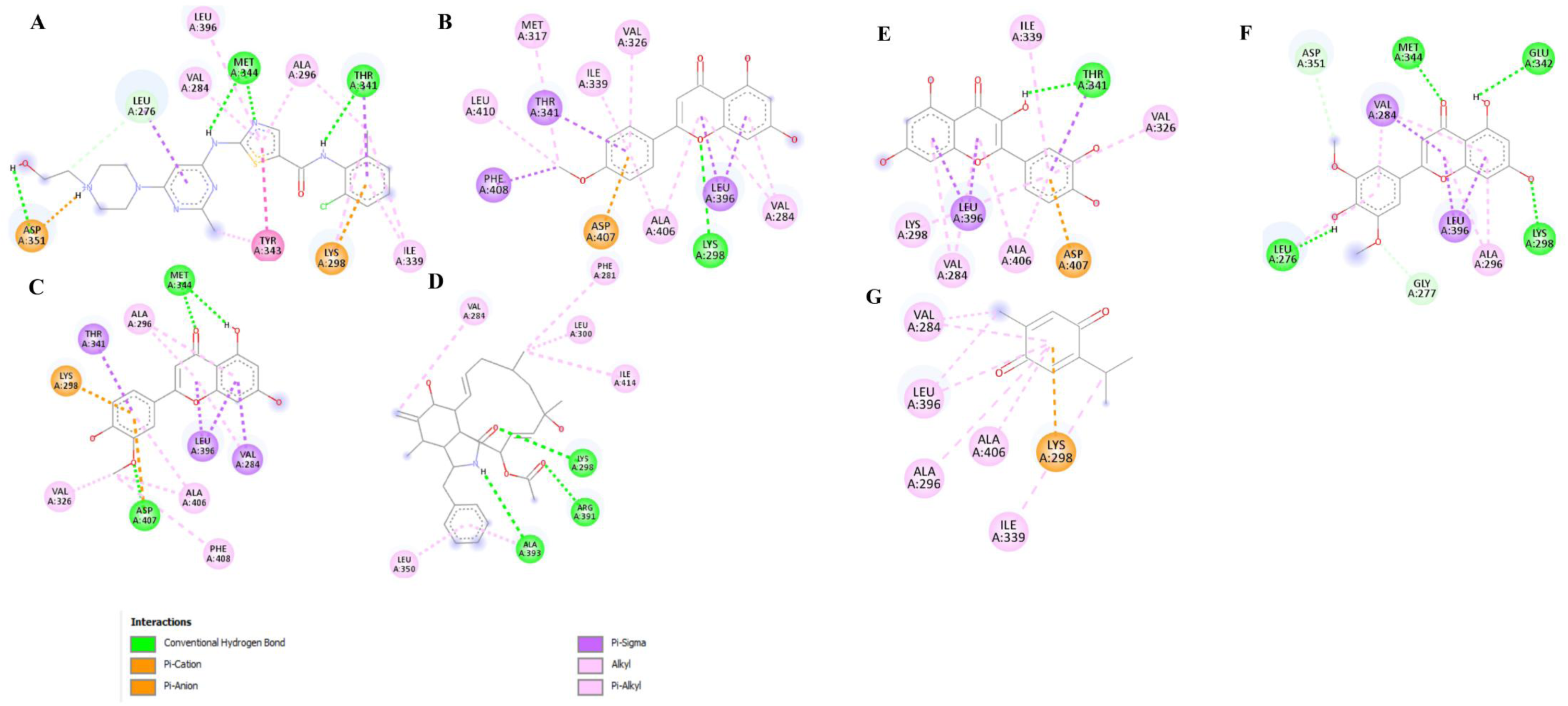
2D protein-ligand interaction diagram of SRC protein with compounds; A: Dasatinib (reference), B: Linarigenin, C: Chryseriol, D: Cytochalasin H, E: Quercetin, F: Tricin and G: Thymoquinone

### Molecular dynamics (MD) simulations

The docking results were verified using a 100 ns MD simulation, providing significant structural insights such as conformational changes and stability of the protein-ligand complex. The RMSD and RMSF analyses provided insights into structural dynamics (Fig 13 and Fig 14). Fig 13 depicts the RMSD plots for the ten target proteins, AKT1, CTNNB1, PIK3CA, STAT3, MAPK3, CDK4, CDK6, JAK1, JAK2, and SRC and their respective compounds. A (0.1-0.2) nm deviation range between the RMSD of the ligand in the complex is considered acceptable and stable, while deviations reaching up to (0.3-0.4) nm are still acceptable for some compounds, depending on the presence of specific flexible regions. The evaluation is based on the mean Root Mean Square Deviation (RMSD), where a lower value signifies a more stable and rigid binding pose, while a higher value indicates greater conformational flexibility. The standard deviation further quantifies the extent of fluctuations in each compound’s binding.

**Fig 13.**
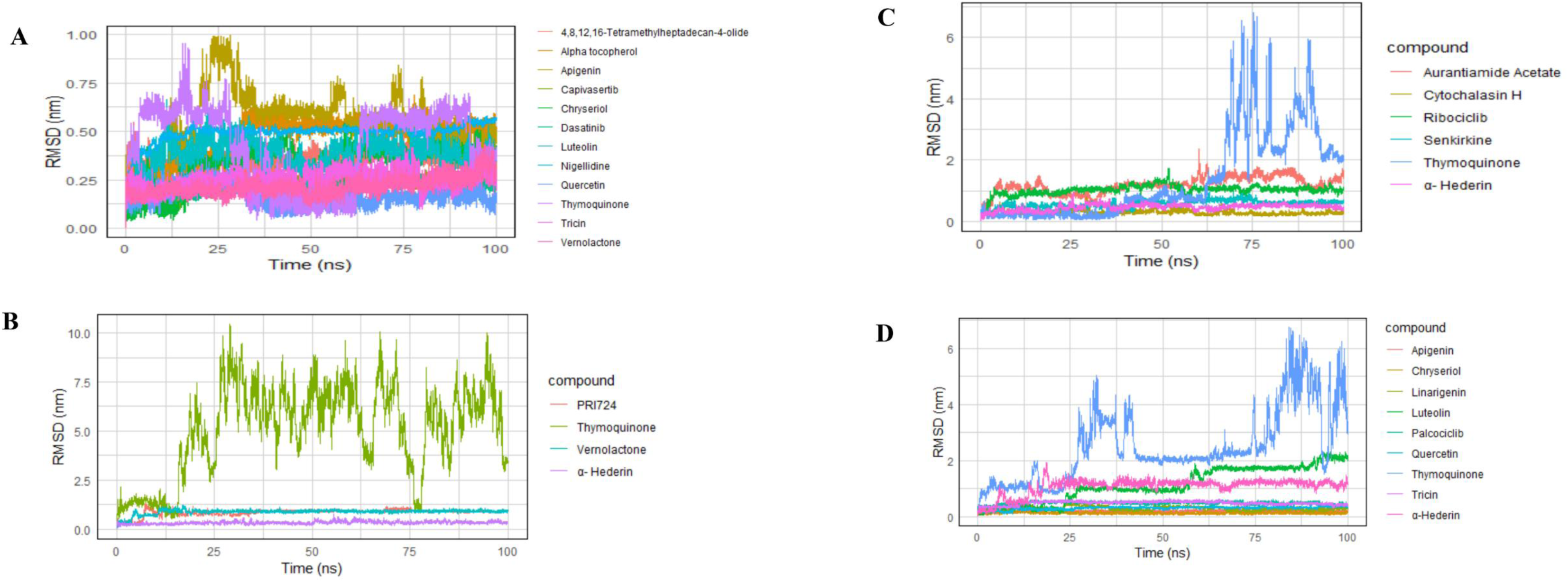
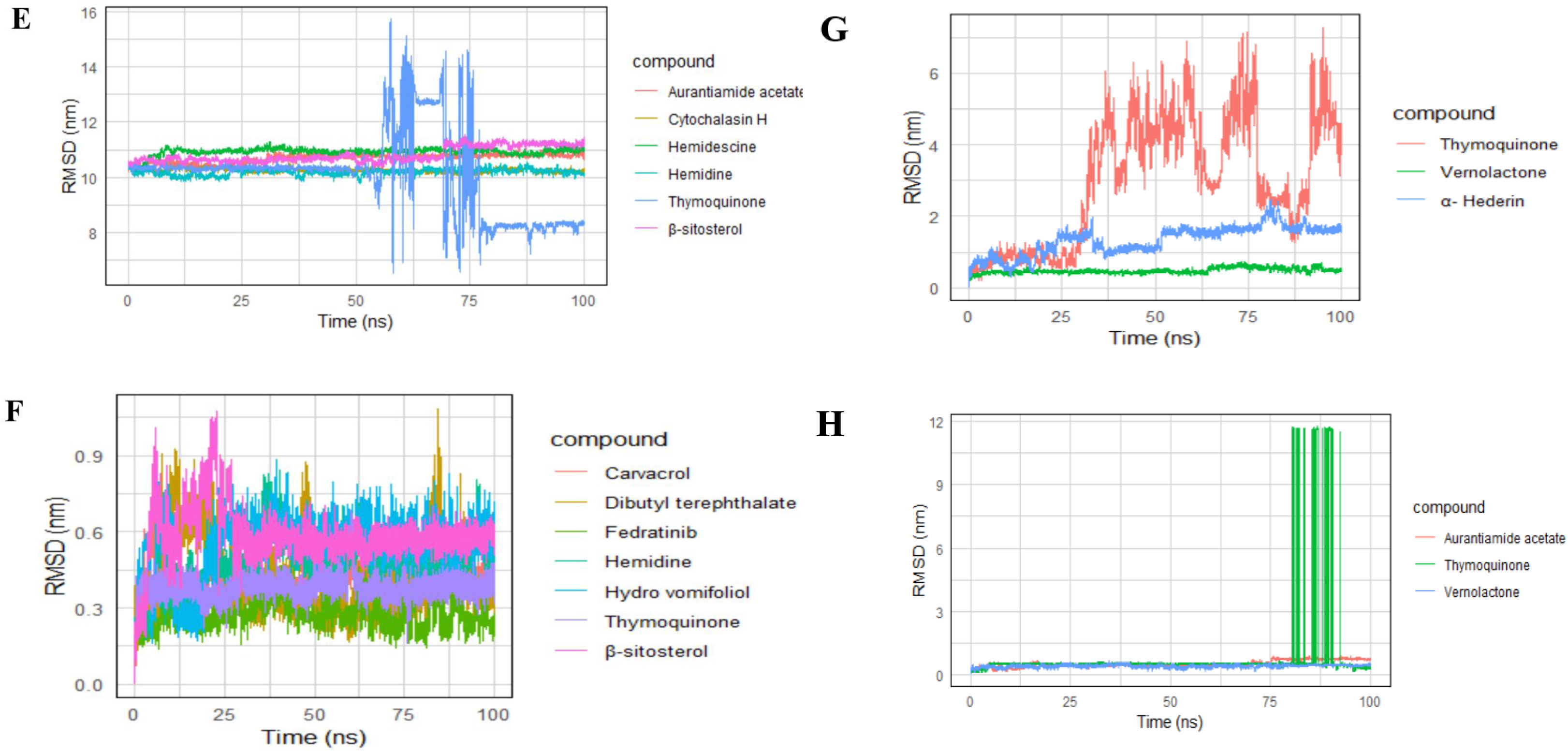
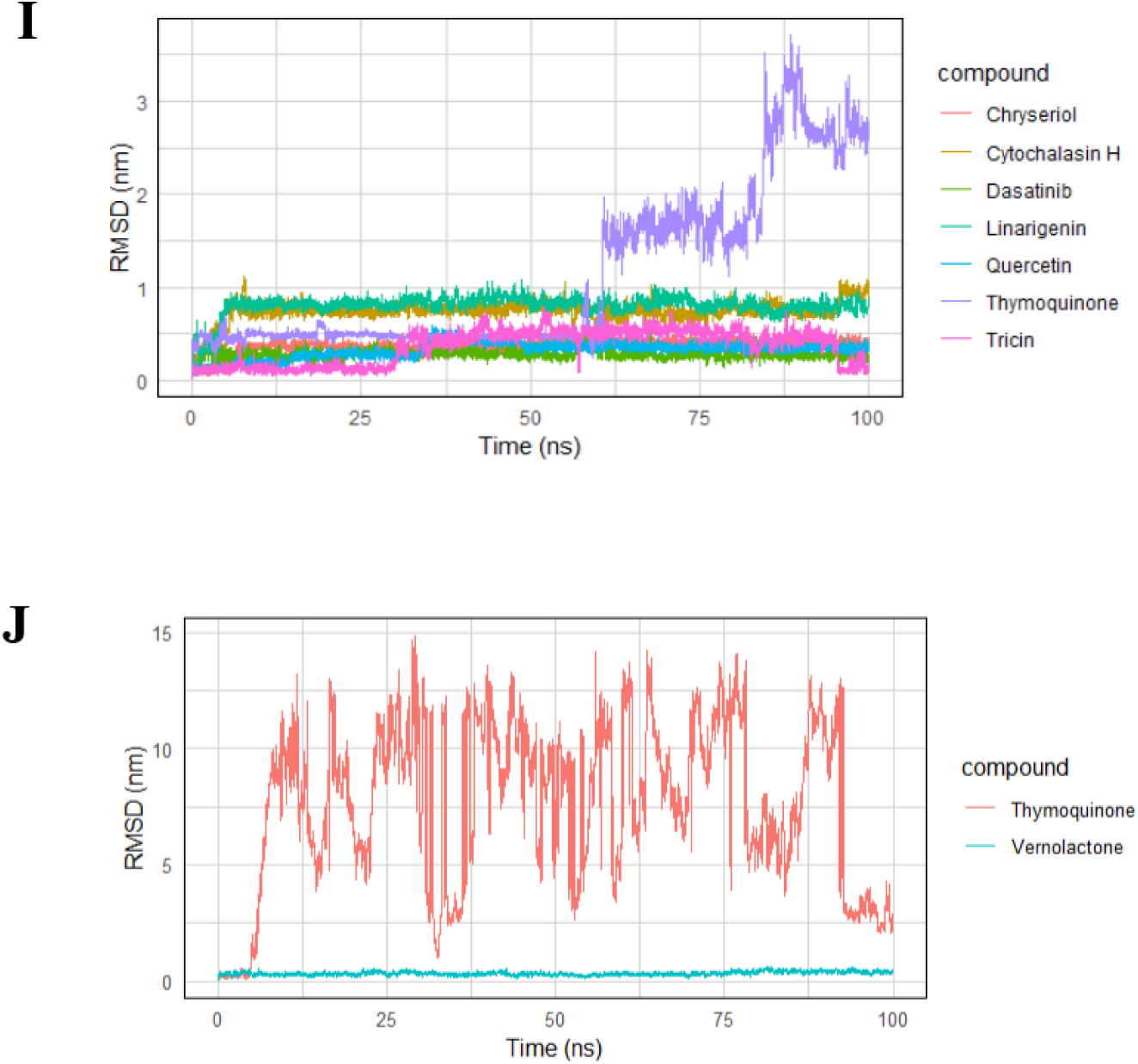
The analysis of the root mean square deviation (RMSD) of the target proteins and their respective compounds: A: AKT1 complex, B: CTNNB1 complex, C: CDK4 complex, D: CDK6 complex, E: JAK1 complex, F: JAK2 complex, G: MAPK3 complex, H: PIK3CA complex, I: SRC complex, J: STAT3 complex

**Fig 14.**
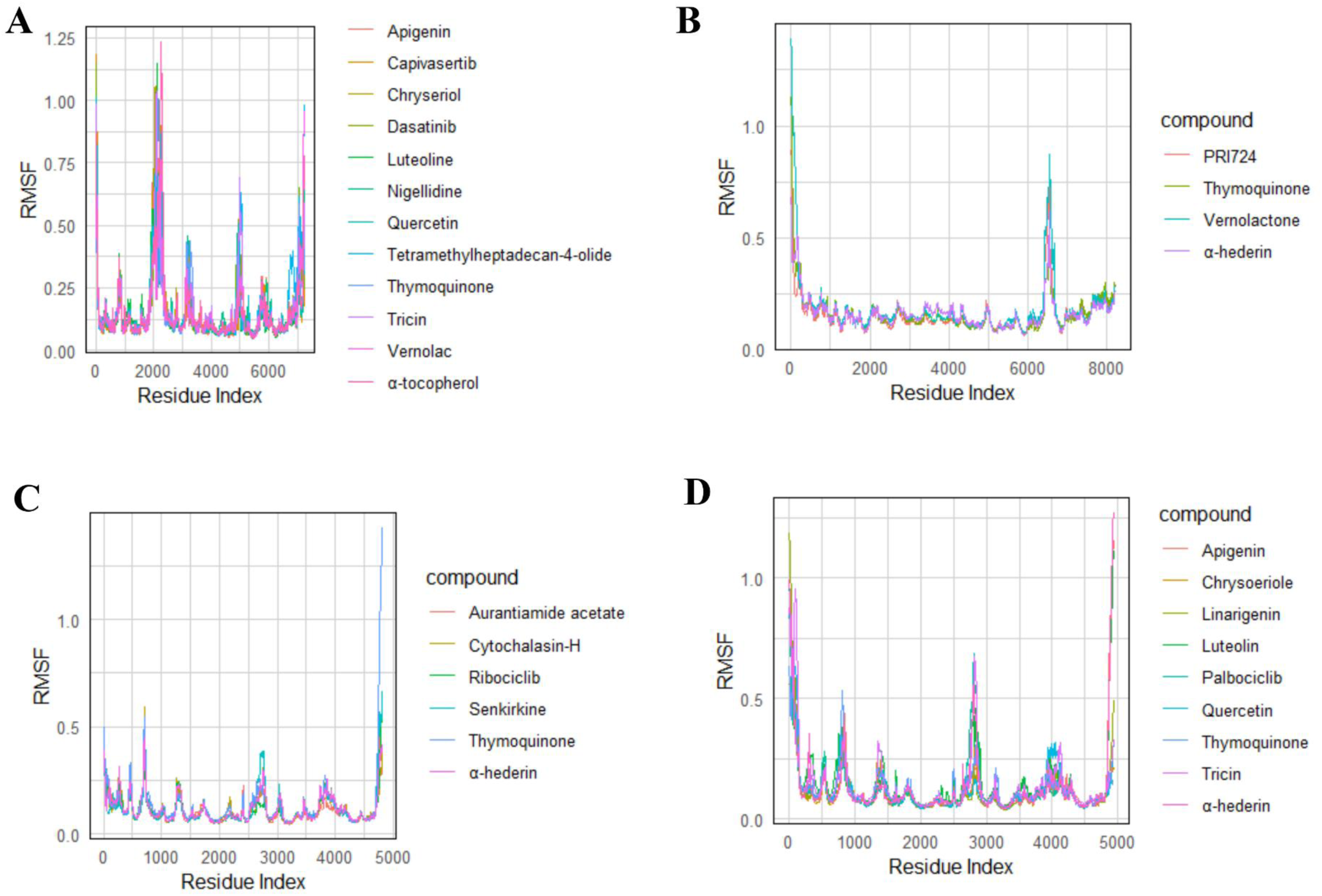
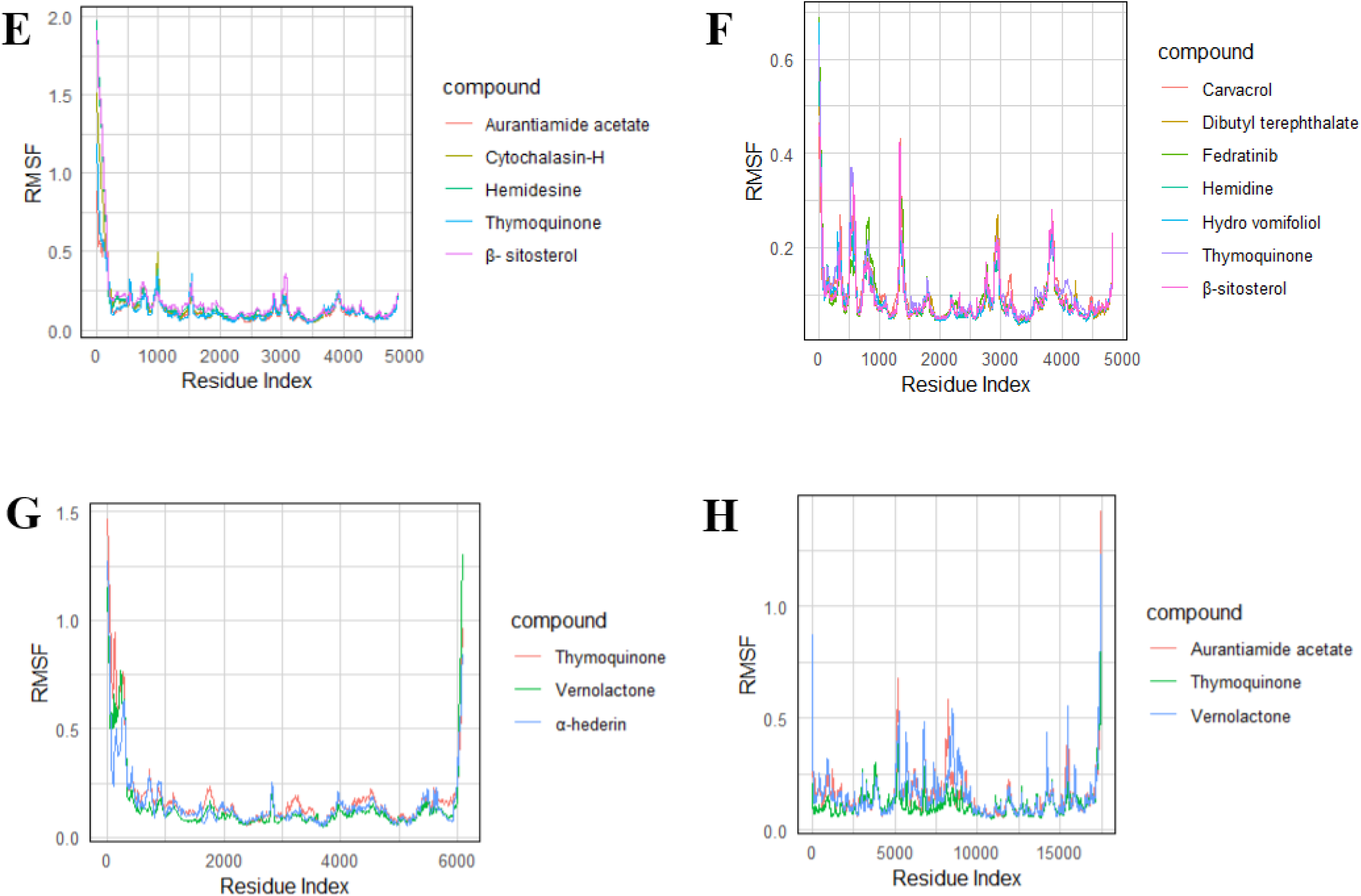
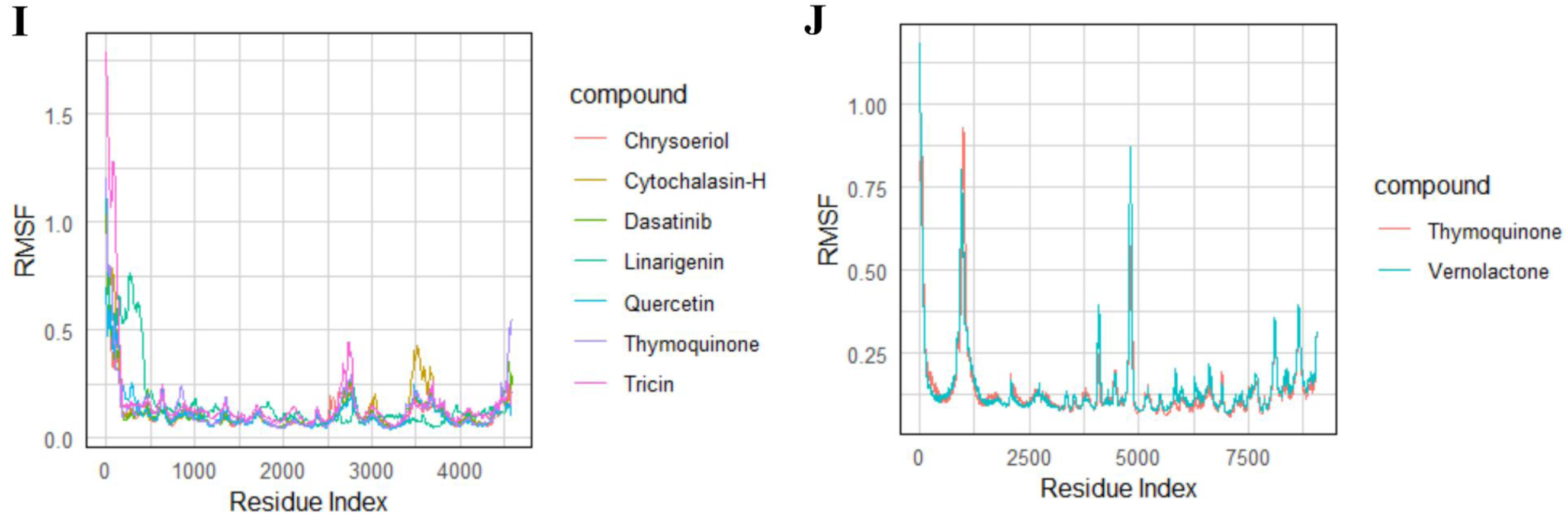
The analysis of the root mean square fluctuation (RMSF) of the target proteins and their respective compounds; A: AKT1 complex, B: CTNNB1 complex, C: CDK4 complex, D: CDK6 complex, E: JAK1 complex, F: JAK2 complex, G: MAPK3 complex, H: PIK3CA, complex, I: SRC complex, J: STAT3 complex

For the AKT1 target protein, a broad spectrum of stability profiles was observed. Quercetin (0.17 ± 0.05 nm) stands out as the most stable binder, with the lowest mean RMSD. This low deviation signifies a highly stable binding and minimal conformational fluctuations within the active site. The reference compounds, Dasatinib (0.21 ± 0.04 nm) and Capivasertib (0.24 ± 0.03 nm), and the compound Vernolactone (0.21 ± 0.05 nm) also demonstrate high stability. Tricin (0.25 ± 0.13 nm), Chryseriol (0.33 ± 0.10 nm), 4,8,12,16-Tetramethylheptadecan-4-olide (0.35 ± 0.05 nm), and Luteolin (0.38 ± 0.07 nm) represent moderately stable binding, showing mean RMSD values in the middle range. In contrast, Thymoquinone (0.43 ± 0.20 nm), Alpha tocopherol (0.47 ± 0.11 nm), Nigellidine (0.50 ± 0.05 nm), and Apigenin (0.53 ± 0.18 nm) displayed the highest RMSD values and significant fluctuations, indicating the least stable and most dynamic interactions.

In the β-catenin system, α-Hederin (0.33 ± 0.07 nm) is the most stable compound, with the lowest mean RMSD, indicating a highly consistent binding pose. The reference control, PRI724 (0.86 ± 0.17 nm), and Vernolactone (0.88 ± 0.13 nm) show moderate stability. In contrast, Thymoquinone (5.08 ± 2.29 nm) is the least stable, with the highest mean RMSD and a very large standard deviation, suggesting significant movement and flexibility within the binding site.

For the CDK4 protein, Cytochalasin H (0.29 ± 0.07 nm) is the most stable compound, maintaining a highly consistent and rigid binding pose. α-Hederin (0.46 ± 0.10 nm) and Senkirkine (0.57 ± 0.12 nm) show moderate stability. The reference control, Ribociclib (1.02 ± 0.17 nm), along with Aurantiamide Acetate (1.18 ± 0.27 nm), are considerably less stable. Thymoquinone (1.36 ± 1.41 nm) is the least stable, with the highest mean RMSD and the largest standard deviation, suggesting significant movement and flexibility.

Within the CDK6 system, Chryseriol (0.13 ± 0.03 nm) is the most stable compound, indicating it maintains a highly consistent and rigid binding pose. Apigenin (0.20 ± 0.03 nm), Quercetin (0.31 ± 0.05 nm), and Linarigenin (0.30 ± 0.07 nm) also demonstrate high stability. The reference control, Palcociclib (0.48 ± 0.05 nm), and Tricin (0.49 ± 0.07 nm) show slightly less stability. α-Hederin (1.08 ± 0.29 nm) and Luteolin (1.15 ± 0.62 nm) are notably less stable, while Thymoquinone (2.49 ± 1.29 nm) is the least stable, with the highest mean RMSD and a large standard deviation.

For the JAK1 system, a unique stability profile is observed. Thymoquinone (10.10 ± 1.48 nm) is the most stable compound, with the lowest mean RMSD, despite its high standard deviation, which suggests significant conformational fluctuation. In contrast, Hemidescine (10.92 ± 0.14 nm) is the least stable with the highest mean RMSD. Other compounds, including Hemidine (10.19 ± 0.11 nm), Cytochalasin H (10.26 ± 0.07 nm), Aurantiamide acetate (10.69 ± 0.16 nm), and β-sitosterol (10.81 ± 0.27 nm), fall between these two extremes.

In analyzing the stability of the JAK2 protein, Fedratinib (0.28 ± 0.06 nm) serves as the reference control and demonstrates the highest stability. All other compounds, including Carvacrol (0.42 ± 0.06 nm), Dibutyl terephthalate (0.43 ± 0.15 nm), Thymoquinone (0.38 ± 0.04 nm), Hemidine (0.48 ± 0.10 nm), Hydro vomifoliol (0.55 ± 0.12 nm) and β-sitosterol (0.58 ± 0.11 nm) exhibit higher mean RMSD values, indicating they are less stable than the reference control.

The analysis for the MAPK3 system reveals Vernolactone (0.48 ± 0.07 nm) maintains a lower mean RMSD suggesting that it is the most stable compound maintaining a structural conformational stability. Meanwhile, Thymoquinone has a mean RMSD of 2.99 ±1.75 nm. This value suggests a significant degree of conformational movement, indicating that the compound is not highly stable in its binding to the MAPK3 protein.

For the STAT3 protein, Vernolactone (0.34 ± 0.08 nm) is the more stable compound, as indicated by its lower mean RMSD, suggesting it maintains a more consistent and rigid binding pose. Thymoquinone (7.69 ± 3.54 nm) is the least stable with a significantly higher mean RMSD and a large standard deviation, which suggests substantial movement and flexibility.

In the PIK3CA system, Vernolactone (0.40 ± 0.07 nm) is the most stable compound, indicating it maintains a more consistent and rigid binding pose. Aurantiamide acetate (0.50 ± 0.16 nm) is moderately stable. In contrast, Thymoquinone (0.79 ± 1.84 nm) is the least stable with the highest mean RMSD and a large standard deviation, suggesting significant movement and flexibility within the binding site.

For the SRC protein, Dasatinib (0.28 ± 0.05 nm) serves as the reference control and is the most stable compound. The other compounds are all less stable, as evidenced by their higher mean RMSD values. Quercetin (0.32 ± 0.09 nm), Tricin (0.36 ± 0.19 nm), and Chryseriol (0.37 ± 0.07 nm) are relatively stable binders. In contrast, Cytochalasin H (0.74 ± 0.12 nm), Linarigenin (0.80 ± 0.11 nm), and especially Thymoquinone (1.11 ± 0.87 nm) are considerably less stable, with Thymoquinone having the highest mean RMSD and a very high standard deviation, indicating the most flexible binding pose.

RMSF (Root Mean Square Fluctuation) is a key metric in molecular dynamics simulations that measures the average deviation of an atom or residue from its reference position, providing insight into the flexibility of a protein-ligand complex. RMSF analysis was employed to evaluate the flexibility of amino acid residues and ligand atoms during the simulation [26,27]. A lower RMSF value indicates a more rigid and stable complex, suggesting a stronger binding interaction, while a higher RMSF points to a more flexible and less stable complex.

Across the provided datasets, the RMSF analysis reveals a range of stability among the protein-ligand pairs. For instance, the JAK2 protein demonstrates the most stable interactions overall, with its ligands having the lowest mean RMSF values in the dataset. Dibutyl terephthalate and Hemidine stand out with exceptionally low RMSF values of (0.090 ± 0.054) nm and (0.090 ± 0.058) nm, respectively, indicating highly rigid complexes. In contrast, the most flexible interactions were observed for JAK1 with β-sitosterol (0.195 ± 0.255 nm) and MAPK3 with Thymoquinone (0.192 ± 0.188 nm). The large standard deviations for these compounds suggest significant fluctuations and less-defined binding poses. The data for AKT1, CTNNB1, CDK4, CDK6, PIK3CA, and SRC fall within this range, each showing a clear hierarchy from most stable to least stable ligand based on their mean RMSF values

### *In vitro* antiproliferative activity

The antiproliferative effects of Vernolac on three cancer cell lines (MCF-7, Caco-2, and NTERA-2 cl.D1) and a normal cell line (MCF-10A) were evaluated using the SRB assay following 24 h and 48 h post-incubation. As Table 8 shows, a dose-dependent reduction in cell viability was observed across the tested cancer cell lines. The IC_50_ values demonstrated increased antiproliferative activity with prolonged exposure (Fig 15). Importantly, the non-tumorigenic MCF-10A cells demonstrated higher IC_50_ values at 24 h and 48 h, suggesting a considerably lower antiproliferative effect of Vernolac on normal cells compared to cancer cells. These findings suggest a potential selective antiproliferative effect of Vernolac toward cancer cells.

**Fig 15.**
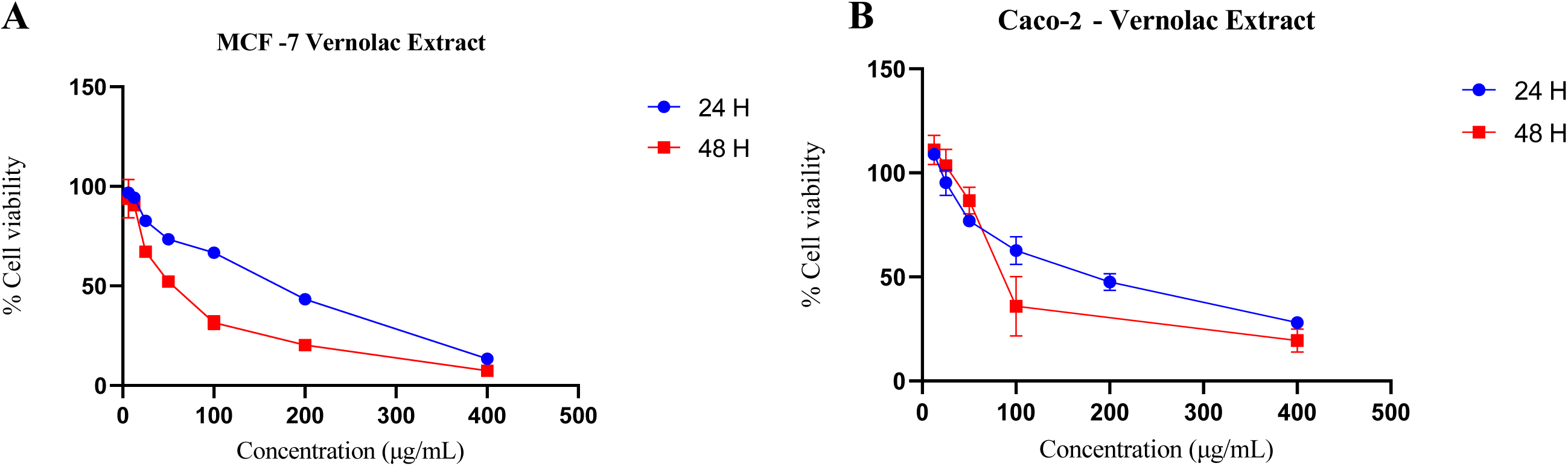
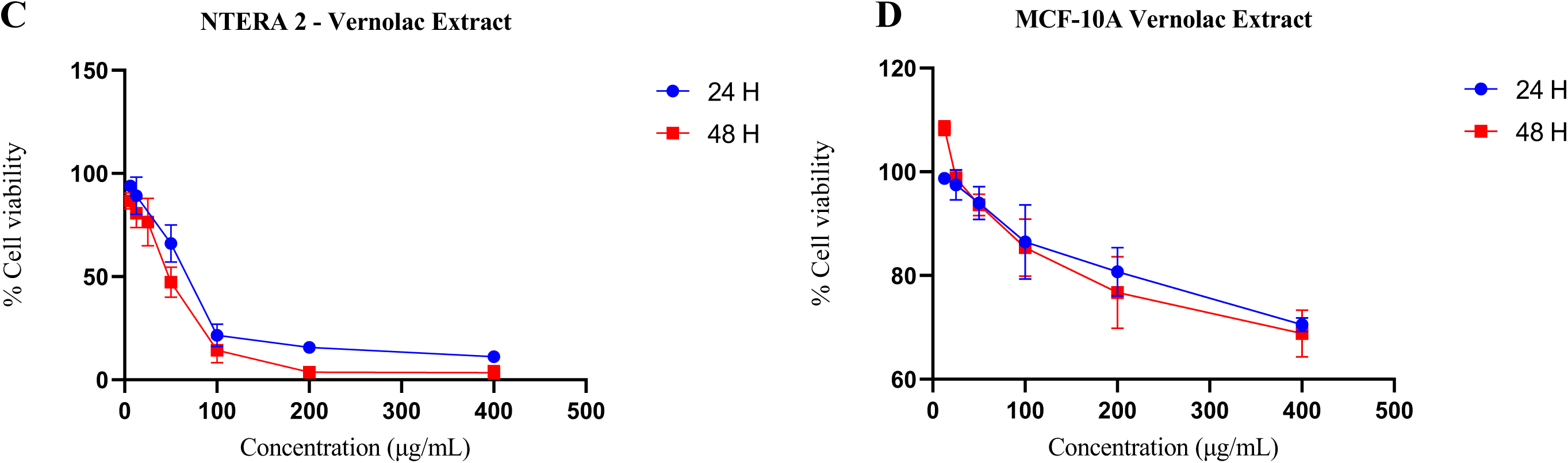
Cell viability (%) of MCF-7, Caco-2, NTERA-2cl.D1, and MCF-10A cells following treatment with Vernolac, assessed by SRB assay at 24 h and 48 h post-incubation. A: MCF-7, B: Caco-2, C: NTERA-2cl.D1, D: MCF-10A

**Table 8.**
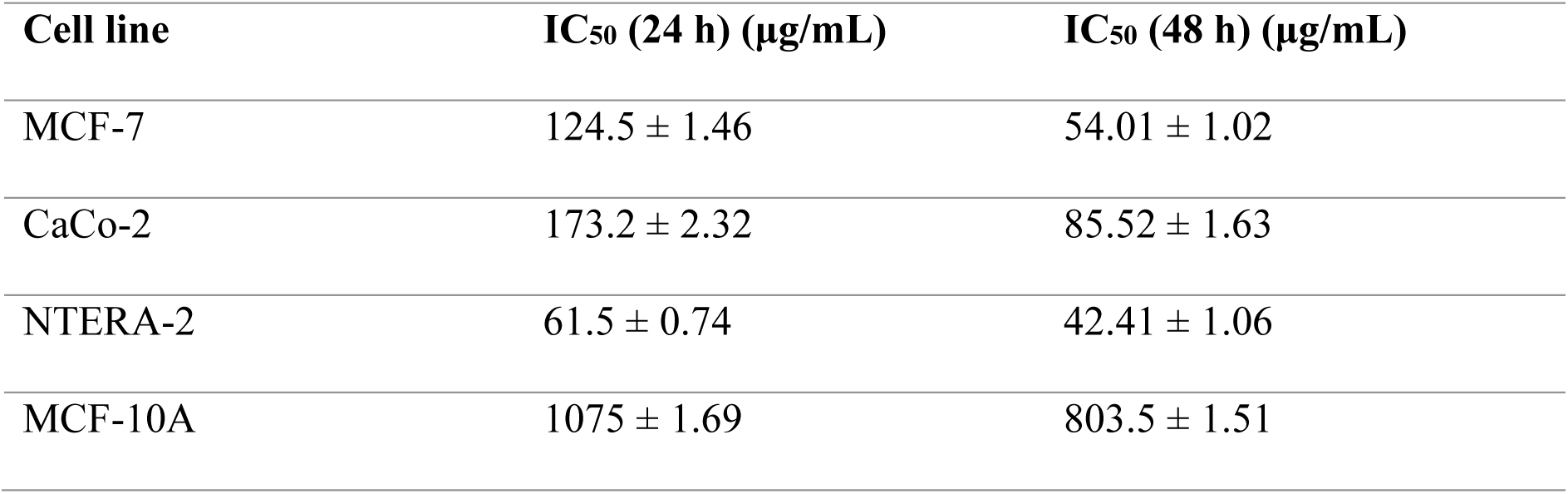
The IC_50_ values (µg/mL) of Vernolac in MCF-7, Caco-2, NTERA 2, and MCF-10A cells at 24 h and 48 h incubation periods, as determined by SRB assay.

| Cell line | IC<sub>50</sub> (24 h) (µg/mL) | IC<sub>50</sub> (48 h) (µg/mL) |
| --- | --- | --- |
| MCF-7 | 124.5 ± 1.46 | 54.01 ± 1.02 |
| CaCo-2 | 173.2 ± 2.32 | 85.52 ± 1.63 |
| NTERA-2 | 61.5 ± 0.74 | 42.41 ± 1.06 |
| MCF-10A | 1075 ± 1.69 | 803.5 ± 1.51 |

## Discussion

Network pharmacology is an emerging paradigm in drug discovery and development, offering a systematic approach to explore complex interactions between compounds, molecular targets, and disease pathways [18,19]. In contrast to the conventional “one drug-one target” model, network pharmacology acknowledges the multifactorial nature of diseases such as cancer. This is particularly beneficial for investigating the mechanisms of action of polyherbal formulations that act through multiple targets and pathways simultaneously. The present study employed a network pharmacology-based approach to explore the potential mechanisms of action of Vernolac, a polyherbal nutraceutical composed of five traditionally used medicinal plants in Sri Lanka. The formula-herb-compound-target-disease network, cluster analysis of the core PPI network, hub node analysis, and topology analysis revealed a set of key hub proteins, including AKT1, EGFR, STAT3, CTNNB1, BCL2, JUN, SRC, MAPK3, and JAK1. These proteins are commonly dysregulated in various types of cancer [28,29]. KEGG pathway enrichment analysis of Vernolac revealed that these targets are significantly enriched in several pathways associated with prostate cancer, glioma, non-small lung carcinoma (NSCLC), endometrial cancer, bladder cancer, pancreatic cancer, renal cell carcinoma, and myeloid leukemias (acute and chronic myeloid leukemias). Results of the present study highlight the broad-spectrum potential of Vernolac against multiple cancer types. Literature reports that several major phytochemicals found in Vernolac have been previously experimentally validated for their modulatory effects on these hub proteins.

Vernolactone, a sesquiterpene lactone isolated from *V. zeylanica*, mediated significant cytotoxic effects in MDA-MB-231 breast cancer cells, with minimal effects on normal mammary epithelial cells (MCF-10A). Moreover, this study reports that Vernolactone exerts pro-apoptotic effects by upregulating *p53* and *Bax*, downregulating *survivin* and Heat Shock Protein (HSP) complex-related genes [30]. Abeysinghe et al. report that vernolactone mediates its antiproliferative activity via apoptosis and autophagy in cancer stem-like cells (NTERA-2), with minimal antiproliferative effects in noncancerous peripheral blood mononuclear cells (PBMC) [31]. As evident from the gene expression results of the same study, vernolactone significantly downregulates the expression of PI3K, AKT, and mTOR, demonstrating its potential to target CSCs, inhibiting the PI3K/Akt/mTOR pathway [31]. Molecular docking and dynamics simulation analyses of the present study identified β-catenin (CTNNB1) as a novel target of vernolactone. To our knowledge, this is the first report to suggest a direct interaction between vernolactone and β-catenin, a key player in the Wnt/β-catenin signaling pathway. This novel finding broadens the mechanistic understanding of the anticancer potential of vernolactone, particularly targeting CSCs.

Thymoquinone, a phytochemical found in *N. sativa*, exhibits anticancer and chemopreventive effects across a broad range of cancer types, targeting multiple molecular pathways [32]. According to previous *in vitro* and *in vivo* studies, its mechanism of action mainly involves the induction of apoptosis via reactive oxygen species (ROS) generation [33], modulating pro- and anti-apoptotic proteins such as Bax, Bcl-2, and caspases, and inhibiting key signaling pathways including PI3K/Akt, and JAK/STAT [32].

Carvacrol, a monoterpenoid phenol abundant in *N. sativa* [34], exhibits broad-spectrum anticancer effects by modulating multiple signaling pathways that align closely with the signaling pathways and hub nodes identified in the present study. According to the literature, its anticancer mechanisms involve the modulation of Bax, Bcl-2, cytochrome c, CDK4/6, p53, caspases (3/6/7/8/9), and cyclins (A/B/D1) [34,35]. Carvacrol also suppresses key signaling pathways including PI3K/Akt, MAPKs (p-ERK, p-JNK, p-38), and ROS generation [34–38]. Previous studies conducted on breast cancer cells (MDA-MB-231 and MCF-7) demonstrate that carvacrol mediates cell growth inhibition and apoptosis by upregulating Bax and PI3K/p-AKT and TRPM7 suppression [38–40]. Similarly, a plethora of *in vitro* cancer cell models including cervical cancer (HeLa cells), choriocarcinoma (JAR and JEG3 cells), lung cancer (A549 and H460), colorectal cancer (Caco-2, HT-29, LoVo cells), prostate cancer (PC-3 cells), melanoma (A375 cells), gastric adenocarcinoma (AGS cells), glioblastoma (U87 cells), and liver cancer (HepG2 cells) have been tested with carvacrol and has strongly validated its anticancer potential [35,36,41–43]. Importantly, carvacrol downregulates STAT3, JAK2 and MMP2 [43,44], three important targets identified by the present network pharmacology study. These findings are further supported by *in vivo* studies demonstrating carvacrol’s ability to suppress tumor growth [44,45], highlighting its potential contribution to the multi-targeted anticancer effects of Vernolac.

Another prominent phytochemical present in Vernolac is α-hederin, a triterpenoid found in *N. sativa*. Previous studies show that α-hederin induces ROS-dependent activation of AMPK/mTOR signaling pathway in colorectal cancer cells (HCT116 and HCT8), leading to both apoptosis and autophagy [46]. It also disrupts mitochondrial membrane potential, elevates Bax/Bcl-2 ratio, activates caspases (3/9) via ROS-mediated mitochondrial apoptosis in gastric (HGC27/DPP), and breast cancer (MCF-7 and MDA-MB-231) models [47,48]. Moreover, α-hederin reduces p-AKT, p-PI3K, and p-mTOR levels in oral cancer cells (SCC-25), thereby suppressing the PI3K/Akt/mTOR pathway [49]. According to literature, α-hederin exhibits a broad-spectrum anticancer potential against numerous in vitro and in vivo cancer models, including ovarian, lung, colorectal, esophageal, and liver cancer [50]. A recent study reports that α-hederin is a potent inhibitor of Wnt/β-catenin pathway genes (cyclin D1 and CD44), and induces apoptosis in breast CSCs [51]. Cyclin D1 binds with CDK4 and CDK6, forming an active complex that promotes cell cycle progression [52]. Therefore, inhibition of cyclin D1 may reduce CDK4 activity by limiting the formation of the cyclin D1-CDK4 complex. However, to the best of our knowledge, no prior studies have reported a direct interaction between α-hederin and CDK4. Findings of the present study suggest that α-hederin directly targets CDK44, highlighting a novel mechanism of action that may contribute to its anticancer effects, independent of its modulation of Cyclin D1.

Quercetin is a flavonoid abundant in *H. indicus* and *S. glabra* with well-established anticancer mechanisms through numerous signaling pathways, including Wnt/β-catenin, MAPK/ERK1/2, p53, JAK/STAT, AMPKα/ASK1/p38, RAGE/PI3K/AKT/mTOR axis, and NF-κB [53]. As evident in the literature, quercetin exhibits ROS-mediated anticancer activity against CSCs [54]. Quercetin also alters Numbl and Notch levels in pancreatic ductal adenocarcinoma cells, inhibiting the self-renewal and aggressiveness of CSCs [55]. According to Seo et al., quercetin induces caspase-dependent extrinsic apoptosis in HER2-overexpressing BT-474 breast cancer cells by inhibiting STAT3 signaling. Their study further highlights the ability of quercetin to prevent or treat HER2-overexpressing breast cancer [56]. Quercetin has demonstrated broad-spectrum anticancer activity against a variety of cancer types, including lung, oral, breast, ovarian, colon, and osteosarcoma.

Among the key phytochemicals of Vernolac, nigellidine is an indazole-type alkaloid [57]. Although experimental validation studying the mechanisms of action of the pure compound, particularly in cancer treatment, is currently limited, an *in silico* study highlights the highest binding effectiveness between nigellidine and HER2 [58]. The results of the present study revealed a strong binding affinity towards AKT1, a serine/threonine kinase that plays a pivotal role in cell proliferation, survival, and apoptosis [59]. To the best of our knowledge, this is the first study to reveal a direct interaction between nigellidine and AKT1, particularly in the context of cancer. The molecular docking results demonstrated a strong binding within the AKT1 active site, and molecular dynamics simulations further confirmed the stability of the protein-ligand complex. This novel finding suggests that nigellidine may exert its anticancer effects through modulations of the PI3K/Akt/mTOR pathway. A previous molecular docking study reports that nigellidine binds strongly to HDAC1, indicating its potential to inhibit histone deacetylase activity [60]. HDAC1 is involved in maintaining the survival of CML cells [61]. Importantly, the CML pathway was identified as significantly enriched in our KEGG pathway analysis, and HDAC1 was identified as a potential target of nigellidine. Therefore, results of the present study further support this mechanistic rationale for nigellidine-mediated epigenetic interferences in CML.

In the present study, network pharmacology integrated with molecular docking and dynamics simulations revealed several novel compound-target interactions relevant to cancer therapy. The present study reports for the first time a strong binding affinity of 4,8,12,16-tetramethylheptadecan-4-olide and tricin to AKT1, a key cancer-related target. Tricin-AKT1 interaction has been previously studied in the context of cardiovascular and neuropathology, including premature ventricular beats [62] and cerebral ischemia/reperfusion injury [63]. However, the relevance of tricin-AKT1 interaction in oncogenic signaling is newly identified herein. Previous studies have reported the anticancer activity of cytochalasin H in lung cancer cells, through modulation of targets such as Bcl-xL, Bcl-2, caspase 3, Bax, p53, and signaling pathways including PI3K/Akt/P70S6K and ERK1/2 [64,65]. However, our study is the first to report CDK4 as a novel molecular target of cytochalasin H, suggesting an additional mechanism by which it may exert antiproliferative effects. These findings expand the understanding of Vernolac’s mechanism of action while suggesting new avenues for targeting key cancer-related proteins.

A polyherbal decoction comprising *N. sativa* seeds, *H. indicus* roots, and *S. glabra* rhizomes in equal proportions has been traditionally utilized for many years by a family of Ayurvedic physicians in Sri Lanka to treat cancers, reflecting its long-standing ethnopharmacological application in cancer therapy [15]. According to the literature, this decoction offers significant protection against chemically induced liver cancer in rats without notable toxicity [16,17]. The decoction has been previously demonstrated to have cytotoxic effects against human hepatoma (HepG2) cells [13,14]. Moreover, the anticancer activity of this decoction has been reported to be mediated through multiple mechanisms, including protection against oxidative damage (12), anti-inflammatory activity [11], and upregulation and downregulation of antiapoptotic and proapoptotic genes [15].

Although chemo- and radiotherapy remain standard treatment modalities in cancer management, their effectiveness is often compromised due to drug resistance, tumor radio-resistance, and severe side effects. In this context, phytochemicals have emerged as potent chemo- and radiosensitizers [66]. According to the literature, phytochemicals such as thymoquinone, quercetin, and lupeol have demonstrated strong chemosensitizing and radiosensitizing effects by modulating key molecular targets, including STAT3, AKT1, TP53, HIF-1α, and PTGS2 [66–68]. These phytochemicals exhibit their chemosensitizing effects via various signaling pathways such as JAK/STAT, NF-kB, Wnt, PI3K/AKT, and EGFR [66]. Phytochemicals such as quercetin have been demonstrated to radiosensitize cancer cells through the expression of p53 and Bax, downregulation of BCL2, and induction of endoplasmic reticulum stress [67].

Chemo and radiotherapy are also associated with various side effects, severely affecting the quality of life of cancer patients. Quercetin, apigenin, ferulic acid, vitamin E, luteolin, and thymoquinone have been reported to demonstrate potent chemoprotective and radioprotective properties [69–71]. They exert this protectivity via a broad range of mechanisms, including antioxidation, anti-inflammation, myeloprotection, antimutagenesis, and immunomodulation [69]. The present network pharmacology-based study also revealed PTGS2, BCL2, PI3K, IL-6, TNF-α, AKT1, and VEGF as key targets within pathways associated with chemoprotection and radioprotection, further supporting the therapeutic relevance of Vernolac’s phytochemicals in alleviating treatment-induced toxicity. Moreover, this dual benefit was further supported by the present *in vitro* cytotoxicity studies. Vernolac demonstrated significant cytotoxicity against breast cancer, colorectal adenocarcinoma, and testicular embryonal carcinoma cells with minimal toxicity in non-tumorigenic mammary epithelial cells. Furthermore, the aforementioned decoction has previously demonstrated protection against bleomycin-induced cytogenetic damage in human peripheral blood lymphocytes (PBLs), indicating significant radioprotective capacity [10]. This suggests a distinct clinical advantage due to its potential to protect normal cells from radiation-induced damage. Notably, Vernolac contains the same three medicinal plants along with two additional components, offering enhanced therapeutic effects.

These findings highlight that Vernolac not only exerts anticancer effects and sensitizes cancer cells to therapy but also improves the quality of life in patients by ameliorating cancer therapy-induced toxicity.

Although the present network pharmacology-based study provides valuable insights into Vernolac’s therapeutic use as a nutraceutical for cancer patients, several limitations remain. The predicted anticancer mechanism of action, chemoradiosensitization effects, chemoprotection, and radioprotection require further experimental validation. Importantly, the *in vivo* metabolic transformation of the phytochemicals was not considered in the present study, which may influence their biological activity and bioavailability. Moreover, novel compound-target interactions identified by the present study require *in vitro* and *in vivo* experimental validation.

## Conclusion

The present study highlights the multi-component, multi-target, and multi-pathway mechanisms underlying the anticancer and chemoradioprotective potential of Vernolac. These findings support the therapeutic relevance of Vernolac, suggesting its potential as an adjunct to conventional cancer treatment.

## Supporting information

**S1 Table. Results of the CytoNCA analysis of 137 core target proteins.**

**S2 Table. Results of the MCODE analysis of the core protein-protein interaction network S3 Table. Protein-ligand interaction data of the molecular docking study**

## References

1. Bray F, Laversanne M, Hyuna, Phd S, Ferlay J, Siegel RL, et al. Global cancer statistics 2022: GLOBOCAN estimates of incidence and mortality worldwide for 36 cancers in 185 countries. A Cancer J Clin [Internet]. 2024 [cited 2024 Aug 19];74(3):229–63. Available from: https://acsjournals.onlinelibrary.wiley.com/doi/10.3322/caac.21834

2. Wu Z, Xia F, Lin R. Global burden of cancer and associated risk factors in 204 countries and territories, 1980–2021: a systematic analysis for the GBD 2021. J Hematol Oncol [Internet]. 2024;17(1):119. Available from: 10.1186/s13045-024-01640-8

3. Zafar A, Khatoon S, Khan MJ, Abu J, Naeem A. Advancements and limitations in traditional anti-cancer therapies: a comprehensive review of surgery, chemotherapy, radiation therapy, and hormonal therapy. Discov Oncol. 2025 Apr;16(1):607.

4. Garg P, Malhotra J, Kulkarni P, Horne D, Salgia R, Singhal SS. Emerging Therapeutic Strategies to Overcome Drug Resistance in Cancer Cells. Cancers (Basel). 2024 Jul;16(13):2478.

5. Manzari-Tavakoli A, Babajani A, Tavakoli MM, Safaeinejad F, Jafari A. Integrating natural compounds and nanoparticle-based drug delivery systems: A novel strategy for enhanced efficacy and selectivity in cancer therapy. Cancer Med [Internet]. 2024 Mar 1;13(5):e7010. Available from: 10.1002/cam4.7010

6. Shakya AK, Naik RR. The Chemotherapeutic Potentials of Compounds Isolated from the Plant, Marine, Fungus, and Microorganism: Their Mechanism of Action and Prospects. J Trop Med. 2022;2022:5919453.

7. Pant S, Joshi S. Emerging insights of nutraceuticals as anticancer. Int J Plant Based Pharm. 2023 Dec 15;3:176–82.

8. Di Napoli R, Balzano N, Mascolo A, Cimmino C, Vitiello A, Zovi A, et al. What Is the Role of Nutraceutical Products in Cancer Patients? A Systematic Review of Randomized Clinical Trials. Nutrients. 2023 Jul;15(14):3249.

9. Afkar A, Afolabi B, Piyathilake P, Perera D, Senathilake K, Wijerathne S, et al. Anticancer Therapeutic Potential of Natural Products. J Desk Res Rev Anal. 2024 Dec 30;2:67–91.

10. Galhena BP, Samarakoon SSR, Thabrew MI, Paul SFD, Perumal V, Mani C. Protective effect of a polyherbal aqueous extract comprised of Nigella sativa (seeds), Hemidesmus indicus (roots), and Smilax glabra (rhizome) on bleomycin induced cytogenetic damage in human lymphocytes. Biomed Res Int. 2017;2017(1):1856713.

11. Galhena PB, Samarakoon SR, Thabrew MI, Weerasinghe GAK, Thammitiyagodage MG, Ratnasooriya WD, et al. Anti-Inflammatory Activity Is a Possible Mechanism by Which the Polyherbal Formulation Comprised of Nigella sativa (Seeds), Hemidesmus indicus (Root), and Smilax glabra (Rhizome) Mediates Its Antihepatocarcinogenic Effects. Evidence-Based Complement Altern Med. 2012;2012(1):108626.

12. Galhena P, Thabrew I, Tammitiyagodage MG, Rachel VAH. Anti-hepatocarcinogenic Ayurvedic herbal remedy reduces the extent of diethylnitrosamine-induced oxidative stress in rats. Pharmacogn Mag. 2009;5(17):19–27.

13. Samarakoon SR, Thabrew I, Galhena PB, De Silva D, Tennekoon KH. A comparison of the cytotoxic potential of standardized aqueous and ethanolic extracts of a polyherbal mixture comprised of Nigella sativa (seeds), Hemidesmus indicus (roots) and Smilax glabra (rhizome). Pharmacognosy Res. 2010;2(6):335–42.

14. Thabrew MI, Mitry RR, Morsy MA, Hughes RD. Cytotoxic effects of a decoction of Nigella sativa, Hemidesmus indicus and Smilax glabra on human hepatoma HepG2 cells. Life Sci. 2005 Aug;77(12):1319–30.

15. Samarakoon SR, Thabrew I, Galhena PB, Tennekoon KH. Modulation of apoptosis in human hepatocellular carcinoma (HepG2 cells) by a standardized herbal decoction of Nigella sativa seeds, Hemidesmus indicus roots and Smilax glabra rhizomes with anti-hepatocarcinogenic effects. BMC Complement Altern Med [Internet]. 2012;12(1):25. Available from: 10.1186/1472-6882-12-25

16. Iddamaldeniya SS, Thabrew MI, Wickramasinghe SMDN, Ratnatunge N, Thammitiyagodage MG. A long-term investigation of the anti-hepatocarcinogenic potential of an indigenous medicine comprised of Nigella sativa, Hemidesmus indicus and Smilax glabra. J Carcinog. 2006 May;5:11.

17. Iddamaldeniya SS, Wickramasinghe N, Thabrew I, Ratnatunge N, Thammitiyagodage MG. Protection against diethylnitrosoamine-induced hepatocarcinogenesis by an indigenous medicine comprised of Nigella sativa, Hemidesmus indicus and Smilax glabra: a preliminary study. J Carcinog. 2003 Oct;2(1):6.

18. Zhao L, Zhang H, Li N, Chen J, Xu H, Wang Y, et al. Network pharmacology, a promising approach to reveal the pharmacology mechanism of Chinese medicine formula. J Ethnopharmacol. 2023 Jun;309:116306.

19. Liu M, Liao H, Peng Q, Huang J, Liu W, Dai M, et al. Comprehensive network pharmacology and experimentation to unveil the therapeutic efficacy and mechanisms of gypenoside LI in anaplastic thyroid cancer. BMC Cancer. 2025 May;25(1):870.

20. Selick HE, Beresford AP, Tarbit MH. The emerging importance of predictive ADME simulation in drug discovery. Drug Discov Today. 2002 Jan;7(2):109–16.

21. Tang Y, Li M, Wang J, Pan Y, Wu FX. CytoNCA: a cytoscape plugin for centrality analysis and evaluation of protein interaction networks. Biosystems. 2015 Jan;127:67– 72.

22. Kesarwani K, Gupta R. Bioavailability enhancers of herbal origin: An overview. Asian Pac J Trop Biomed [Internet]. 2013;3(4):253–66. Available from: https://www.sciencedirect.com/science/article/pii/S222116911360060X

23. Ge SX, Jung D, Yao R. ShinyGO: a graphical gene-set enrichment tool for animals and plants. Bioinformatics [Internet]. 2020 Apr 15;36(8):2628–9. Available from: 10.1093/bioinformatics/btz931

24. Rajagopalan U, Samarakoon S, Tennekoon K, Malavige G, de Silva ED. Screening of five Sri Lankan endemic plants for anti-cancer effects on breast cancer stem cells isolated from MCF-7 and MDA-MB-231 cell lines. Trop J Pharm Res. 2018 Sep 1;17:1825–32.

25. Barabási AL, Gulbahce N, Loscalzo J. Network medicine: a network-based approach to human disease. Nat Rev Genet. 2011 Jan;12(1):56–68.

26. Enayatkhani M, Salimi M, Azadmanesh K, Teimoori-Toolabi L. In-silico identification of new inhibitors for Low-density lipoprotein receptor-related protein6 (LRP6). J Biomol Struct Dyn [Internet]. 2022 Jun 21;40(10):4440–50. Available from: 10.1080/07391102.2020.1857843

27. Wang C, He F, Sun K, Guo K, Lu S, Wu T, et al. Identification and characterization of 7-azaindole derivatives as inhibitors of the SARS-CoV-2 spike-hACE2 protein interaction. Int J Biol Macromol [Internet]. 2023;244:125182. Available from: https://www.sciencedirect.com/science/article/pii/S0141813023020767

28. White BD, Chien AJ, Dawson DW. Dysregulation of Wnt/β-catenin signaling in gastrointestinal cancers. Gastroenterology. 2012 Feb;142(2):219–32.

29. Hahn WC, Bader JS, Braun TP, Califano A, Clemons PA, Druker BJ, et al. An expanded universe of cancer targets. Cell. 2021 Mar;184(5):1142–55.

30. Mendis AS, Thabrew I, Ediriweera MK, Samarakoon SR, Tennekoon KH, Adhikari A, et al. Isolation of a New Sesquiterpene Lactone From Vernonia Zeylanica (L) Less and its Anti-Proliferative Effects in Breast Cancer Cell Lines. Anticancer Agents Med Chem [Internet]. 2019;19(3):410–24. Available from: https://pubmed.ncbi.nlm.nih.gov/30488799/

31. Abeysinghe NK, Thabrew I, Samarakoon SR, Ediriweera MK, Tennekoon KH, Pathiranage VPC, et al. Vernolactone Promotes Apoptosis and Autophagy in Human Teratocarcinomal (NTERA-2) Cancer Stem-Like Cells. Stem Cells Int [Internet]. 2019;2019:6907893. Available from: https://pubmed.ncbi.nlm.nih.gov/31949439/

32. Asaduzzaman Khan M, Tania M, Fu S, Fu J. Thymoquinone, as an anticancer molecule: from basic research to clinical investigation. Oncotarget. 2017 Aug;8(31):51907–19.

33. Yu SM, Kim SJ. The thymoquinone-induced production of reactive oxygen species promotes dedifferentiation through the ERK pathway and inflammation through the p38 and PI3K pathways in rabbit articular chondrocytes. Int J Mol Med. 2015 Feb;35(2):325–32.

34. Sharifi-Rad M, Varoni EM, Iriti M, Martorell M, Setzer WN, del Mar Contreras M, et al. Carvacrol and human health: A comprehensive review. Phyther Res [Internet]. 2018 Sep 1;32(9):1675–87. Available from: 10.1002/ptr.6103

35. Sampaio LA, Pina LTS, Serafini MR, Tavares DDS, Guimarães AG. Antitumor Effects of Carvacrol and Thymol: A Systematic Review. Front Pharmacol. 2021;12:702487.

36. Li L, He L, Wu Y, Zhang Y. Carvacrol affects breast cancer cells through TRPM7 mediated cell cycle regulation. Life Sci [Internet]. 2021;266:118894. Available from: https://www.sciencedirect.com/science/article/pii/S0024320520316477

37. Lim W, Ham J, Bazer FW, Song G. Carvacrol induces mitochondria-mediated apoptosis via disruption of calcium homeostasis in human choriocarcinoma cells. J Cell Physiol. 2019 Feb;234(2):1803–15.

38. Khan I, Bahuguna A, Kumar P, Bajpai VK, Kang SC. In vitro and in vivo antitumor potential of carvacrol nanoemulsion against human lung adenocarcinoma A549 cells via mitochondrial mediated apoptosis. Sci Rep [Internet]. 2018;8(1):144. Available from: 10.1038/s41598-017-18644-9

39. Mari A, Mani G, Nagabhishek SN, Balaraman G, Subramanian N, Mirza FB, et al. Carvacrol Promotes Cell Cycle Arrest and Apoptosis through PI3K/AKT Signaling Pathway in MCF-7 Breast Cancer Cells. Chin J Integr Med. 2021 Sep;27(9):680–7.

40. Arunasree KM. Anti-proliferative effects of carvacrol on a human metastatic breast cancer cell line, MDA-MB 231. Phytomedicine. 2010 Jul;17(8–9):581–8.

41. Heidarian E, Keloushadi M. Antiproliferative and Anti-invasion Effects of Carvacrol on PC3 Human Prostate Cancer Cells through Reducing pSTAT3, pAKT, and pERK1/2 Signaling Proteins. Int J Prev Med. 2019;10:156.

42. Fan K, Li X, Cao Y, Qi H, Li L, Zhang Q, et al. Carvacrol inhibits proliferation and induces apoptosis in human colon cancer cells. Anticancer Drugs. 2015 Sep;26(8):813–23.

43. Khan F, Khan I, Farooqui A, Ansari IA. Carvacrol Induces Reactive Oxygen Species (ROS)-mediated Apoptosis Along with Cell Cycle Arrest at G(0)/G(1) in Human Prostate Cancer Cells. Nutr Cancer. 2017 Oct;69(7):1075–87.

44. Jayakumar S, Madankumar A, Asokkumar S, Raghunandhakumar S, Gokula dhas K, Kamaraj S, et al. Potential preventive effect of carvacrol against diethylnitrosamine-induced hepatocellular carcinoma in rats. Mol Cell Biochem [Internet]. 2012;360(1):51–60. Available from: 10.1007/s11010-011-1043-7

45. Karkabounas S, Kostoula OK, Daskalou T, Veltsistas P, Karamouzis M, Zelovitis I, et al. Anticarcinogenic and antiplatelet effects of carvacrol. Exp Oncol. 2006 Jun;28(2):121–5.

46. Sun J, Feng Y, Wang Y, Ji Q, Cai G, Shi L, et al. α-hederin induces autophagic cell death in colorectal cancer cells through reactive oxygen species dependent AMPK/mTOR signaling pathway activation. Int J Oncol [Internet]. 2019;54(5):1601–12. Available from: 10.3892/ijo.2019.4757

47. Cheng L, Xia TS, Wang YF, Zhou W, Liang XQ, Xue JQ, et al. The anticancer effect and mechanism of α-hederin on breast cancer cells. Int J Oncol [Internet]. 2014;45(2):757–63. Available from: 10.3892/ijo.2014.2449

48. 48. Liu Y, Lei H, Ma J, Deng H, He P, Dong W. α-Hederin Increases The Apoptosis Of Cisplatin-Resistant Gastric Cancer Cells By Activating Mitochondrial Pathway In Vivo And Vitro. Onco Targets Ther. 2019;12:8737–50.

49. Wang H, Wu B, Wang H. Alpha-hederin induces the apoptosis of oral cancer SCC-25 cells by regulating PI3K/Akt/mTOR signaling pathway. Electron J Biotechnol [Internet]. 2019;38:27–31. Available from: https://www.sciencedirect.com/science/article/pii/S0717345819300016

50. Belmehdi O, Taha D, Abrini J, Ming LC, Khalid A, Abdalla AN, et al. Anticancer properties and mechanism insights of α-hederin. Biomed Pharmacother [Internet]. 2023;165:115205. Available from: https://www.sciencedirect.com/science/article/pii/S0753332223009964

51. Saliu TP, Seneviratne NN, Faizan M, Rajagopalan U, Perera DC, Adhikari A, et al. In silico identification and in vitro validation of alpha-hederin as a potent inhibitor of Wnt/β-catenin signaling pathway in breast cancer stem cells. Silico Pharmacol [Internet]. 2024;12(1):31. Available from: 10.1007/s40203-024-00199-z

52. Alao JP. The regulation of cyclin D1 degradation: roles in cancer development and the potential for therapeutic invention. Mol Cancer [Internet]. 2007;6(1):24. Available from: 10.1186/1476-4598-6-24

53. Asgharian P, Tazekand AP, Hosseini K, Forouhandeh H, Ghasemnejad T, Ranjbar M, et al. Potential mechanisms of quercetin in cancer prevention: focus on cellular and molecular targets. Cancer Cell Int [Internet]. 2022;22(1):257. Available from: 10.1186/s12935-022-02677-w

54. Biswas P, Dey D, Biswas PK, Rahaman TI, Saha S, Parvez A, et al. A Comprehensive Analysis and Anti-Cancer Activities of Quercetin in ROS-Mediated Cancer and Cancer Stem Cells. Int J Mol Sci. 2022 Oct;23(19).

55. MacKenzie TN, Mujumdar N, Banerjee S, Sangwan V, Sarver A, Vickers S, et al. Triptolide induces the expression of miR-142-3p: a negative regulator of heat shock protein 70 and pancreatic cancer cell proliferation. Mol Cancer Ther. 2013 Jul;12(7):1266–75.

56. Seo HS, Ku JM, Choi HS, Choi YK, Woo JK, Kim M, et al. Quercetin induces caspase-dependent extrinsic apoptosis through inhibition of signal transducer and activator of transcription 3 signaling in HER2-overexpressing BT-474 breast cancer cells. Oncol Rep. 2016 Jul;36(1):31–42.

57. Atta-ur-Rahman, Malik S, Hasan SS, Choudhary MI, Ni CZ, Clardy J. Nigellidine — A new indazole alkaloid from the seeds of Nigella sativa. Tetrahedron Lett [Internet]. 1995;36(12):1993–6. Available from: https://www.sciencedirect.com/science/article/pii/0040403995002104

58. Bilal A, Tanvir F, Ahmad S, Azam AR, Qasim M, Zafar H. Therapeutical Evaluation of Bioactive Compounds of Nigella Sativa for Her2-Positive Breast Cancer Treatment. J Popul Ther Clin. 2024;31(09):3149–64.

59. Chen L, Kang QH, Chen Y, Zhang YH, Li Q, Xie SQ, et al. Distinct roles of Akt1 in regulating proliferation, migration and invasion in HepG2 and HCT 116 cells. Oncol Rep [Internet]. 2014;31(2):737–44. Available from: 10.3892/or.2013.2879

60. Anwar MJ, Alenezi SK, Azam F, Mahmood D, Imam F, Alharbi KS. Nigella sativa oil alleviates doxorubicin-induced cardiomyopathy and neurobehavioral changes in mice: In vivo: and: in-silico: study. Asian Pac J Trop Biomed [Internet]. 2022;12(7):312–22. Available from: https://journals.lww.com/aptb/fulltext/2022/12070/nigella_sativa_oil_alleviates_doxor ubicin_induced.4.aspx

61. Lernoux M, Schnekenburger M, Dicato M, Diederich M. Epigenetic mechanisms underlying the therapeutic effects of HDAC inhibitors in chronic myeloid leukemia. Biochem Pharmacol [Internet]. 2020;173:113698. Available from: https://www.sciencedirect.com/science/article/pii/S0006295219303971

62. Shuyuan L, Haoyu C. Mechanism of Nardostachyos Radix et Rhizoma–Salidroside in the treatment of premature ventricular beats based on network pharmacology and molecular docking. Sci Rep [Internet]. 2023;13(1):20741. Available from: 10.1038/s41598-023-48277-0

63. Liu Ying, Qu Xiaoning, Yan Mengjun, Li Dalei, Zou Rong. Tricin attenuates cerebral ischemia/reperfusion injury through inhibiting nerve cell autophagy, apoptosis and inflammation by regulating the PI3K/Akt pathway. Hum Exp Toxicol [Internet]. 2022 Jan 1;41:09603271221125928. Available from: 10.1177/09603271221125928

64. Ma Y, Wu X, Xiu Z, Liu X, Huang B, Hu L, et al. Cytochalasin H isolated from mangrove-derived endophytic fungus induces apoptosis and inhibits migration in lung cancer cells. Oncol Rep [Internet]. 2018;39(6):2899–905. Available from: 10.3892/or.2018.6347

65. Ma Y, Xiu Z, Zhou Z, Huang B, Liu J, Wu X, et al. Cytochalasin H Inhibits Angiogenesis via the Suppression of HIF-1α Protein Accumulation and VEGF Expression through PI3K/AKT/P70S6K and ERK1/2 Signaling Pathways in Non-Small Cell Lung Cancer Cells. J Cancer. 2019;10(9):1997–2005.

66. Nisar S, Masoodi T, Prabhu KS, Kuttikrishnan S, Zarif L, Khatoon S, et al. Natural products as chemo-radiation therapy sensitizers in cancers. Biomed Pharmacother [Internet]. 2022;154:113610. Available from: https://www.sciencedirect.com/science/article/pii/S0753332222009994

67. Gong C, Yang Z, Zhang L, Wang Y, Gong W, Liu Y. Quercetin suppresses DNA double-strand break repair and enhances the radiosensitivity of human ovarian cancer cells via p53-dependent endoplasmic reticulum stress pathway. Onco Targets Ther. 2017;17–27.

68. Zhang L, Bai Y, Yang Y. Thymoquinone chemosensitizes colon cancer cells through inhibition of NF-κB. Oncol Lett [Internet]. 2016;12(4):2840–5. Available from: 10.3892/ol.2016.4971

69. Liu YQ, Wang XL, He DH, Cheng YX. Protection against chemotherapy- and radiotherapy-induced side effects: A review based on the mechanisms and therapeutic opportunities of phytochemicals. Phytomedicine [Internet]. 2021;80:153402. Available from: https://www.sciencedirect.com/science/article/pii/S0944711320302336

70. Jain A, Madu CO, Lu Y. Phytochemicals in Chemoprevention: A Cost-Effective Complementary Approach. J Cancer. 2021;12(12):3686–700.

71. Zhang Y, Huang Y, Li Z, Wu H, Zou B, Xu Y. Exploring Natural Products as Radioprotective Agents for Cancer Therapy: Mechanisms, Challenges, and Opportunities. Cancers (Basel). 2023 Jul;15(14):3585.

