## Supplementary material for "A network pharmacology-based approach and molecular docking study to explore the therapeutic potential of a nutraceutical formula (Vernolac) in the treatment of cancer": S3 Table

| **Target protein** | **Bioactive compound** | **Interacting residues** | **Category** |
| --- | --- | --- | --- |
| **AKT1** | Capivesartib (Reference compound) | THR: A:82, VAL: A:271, TYR: A:272, ARG: A:273, ASP: A:274 | Hydrogen bonds |
|  |  | TRP: A:80, LEU: A:264, LYS: A:268, VAL: A:270 | Hydrophobic interactions |
|  |  | ASP: A:292 | Electrostatic interactions |
|  | Dasatinib (Reference compound) | ASN: A:53, THR: A:81, THR: A:82, SER: A:205, LYS: A:268 | Hydrogen bonds |
|  |  | ALA: A:58, GLN: A:79, TRP: A:80, LEU: A:210, TYR: A:263, LEU: A:264, VAL: A:270 | Hydrophobic interactions |
|  |  | ASP: A:292 | Electrostatic interactions |
|  | Nigellidine | VAL: A:270, TRP: A:80, LYS: A:268, LEU: A:210, LEU: A:264 | Hydrophobic interactions |
|  |  | TYR: A:272 | Unfavorable interactions |
|  | Vernolactone | ASN: A:54, GLN: A:79, TYR: A:272, ARG: A:273, TYR: A:326 | Hydrogen bonds |
|  |  | LEU: A:264, VAL: A:270 | Hydrophobic interactions |
|  | Thymoquinone | TRP: A:80, SER: A:205 | Hydrogen bonds |
|  |  | TRP: A:80, LEU: A:210, LEU: A:264, LYS: A:268 | Hydrophobic interactions |
|  | Apigenin | ASN: A: 54, GLN: A: 79, SER: A: 205 | Hydrogen bonds |
|  |  | VAL: A: 270, TRP: A: 80, LYS: A: 268 | Hydrophobic interactions |
|  | Luteoline | ASN: A: 54, ILE: A: 290, THR: A: 211 | Hydrogen bonds |
|  |  | VAL: A: 270, LEU: A: 264, TRP: A: 80, LEU: A: 210 | Hydrophobic interactions |
|  |  | ASP: A: 292 | Electrostatic interactions |
|  | Tetramethylheptadecan-4-olide | THR: A: 82 | Hydrogen bond |
|  |  | VAL: A: 270, TRP: A: 80, LYS: A: 268, TYR: A: 263, LEU: A: 264, LEU: A: 210 | Hydrophobic interactions |
|  | α- tocopherol | LEU: A: 210, LEU: A: 264, TYR: A: 263, LYS: A: 268, TRP: A: 80, VAL: A: 270, ILE: A: 84, TYR: A: 272 | Hydrophobic interactions |
|  | Chryseriol | SER: A: 205, THR: A: 211, ILE: A: 290, GLN: A: 79, ASN: A: 54 | Hydrogen bonds |
|  |  | LEU: A: 210, LEU: A: 264, TRP: A: 80, VAL: A: 270 | Hydrophobic interactions |
|  |  | ASP: A: 292 | Electrostatic interactions |
|  | Quercetin | ASN: A: 54, GLN: A: 79, VAL: A: 271, THR: A: 211, ILE: A: 290 | Hydrogen bonds |
|  |  | VAL: A: 270, TRP: A: 80, LEU: A: 264, LEU: A: 210 | Hydrophobic interactions |
|  |  | ASP: A: 292 | Electrostatic interactions |
|  | Tricin | SER: A: 205, THR: A: 211 | Hydrogen bonds |
|  |  | VAL: A: 270, TRP: A: 80, TYR: A: 272, LEU: A: 264, LEU: A: 210 | Hydrophobic interactions |
|  |  | ASP: A: 292 | Electrostatic Interactions |
| **CTNNB1** | PRI724 (Reference compound) | CYS: A:429, ASN: A:430, LYS: A:435, ARG: A:474, LYS: A:508 | Hydrogen bonds |
|  |  | VAL: A:564 | Hydrophobic interactions |
|  |  | ARG: A:469 | Electrostatic interactions |
|  | Alpha-Hederin | ASP: A: 390, ARG: A: 386, ASN: A: 387 | Hydrogen bonds |
|  |  | HIS: A: 503 | Hydrophobic interactions |
|  |  | ASN: A: 387 | Unfavorable bonds |
|  | Thymoquinone | ASN: A: 426 | Hydrogen bonds |
|  |  | PRO: A: 463, CYS: A: 466 | Hydrophobic interactions |
|  |  | CYS: A: 466 | Electrostatic interactions |
|  | Vernolactone | ARG: A:469, ARG: A:474, LYS: A:508, GLY: A:512 | Hydrogen bonds |
|  |  | CYS: A:429, HIS: A:470 | Hydrophobic interactions |
| **PIK3CA** | Thymoquinone | ASP: A:933 | Hydrogen bonds |
|  |  | ILE: A:800, TYR: A:836, ILE: A:848, MET: A:922, ILE: A:932 | Hydrophobic interactions |
|  |  | TYR: A:836 | Unfavorable bond |
|  | Aurantiamide acetate | SER: A: 774, SER: A: 919 | Hydrogen bonds |
|  |  | MET: A: 772, TRP: A: 780, VAL: A: 850, MET: A: 922, ILE: A: 932, ILE: A: 848 | Hydrophobic interactions |
|  | Vernolactone | SER: A: 773, SER: A: 774, LYS: A: 802, VAL: A: 851 | Hydrogen bonds |
|  |  | MET: A: 922 | Hydrophobic interactions |
| **STAT3** | Vernolactone | CYS: A:251, ARG: A:325, GLN: A:326, PRO: A:333 | Hydrogen bonds |
|  |  | ALA: A:250, PRO: A:256, ARG: A:325 | Hydrophobic interactions |
|  | Thymoquinone | SER: A: 613, GLU: A: 612 | Hydrogen bonds |
|  |  | PRO: A: 639 | Hydrophobic interactions |
|  |  | ARG: A: 609 | Electrostatic interactions |
| **MAPK3** | α-Hederin | GLU: A:50, ASP: A:128, SER: A:170, ASN: A:171 | Hydrogen bonds |
|  |  | ALA: A:52, TYR: A:53, LYS: A:71, CYS: A:183 | Hydrophobic interactions |
|  |  | ASP: A:128 | Unfavorable bond |
|  | Vernolactone | SER: A: 170, GLU: A: 50, LYS: A: 131, GLY: A: 49, ASN: A: 171 | Hydrogen bonds |
|  |  | VAL: A: 56, CYS: A: 183, ALA: A: 69, ILE: A: 48, LEU: A: 173, LEU: A: 124 | Hydrophobic interactions |
|  | Thymoquinone | GLY: A: 186 | Hydrogen bonds |
|  |  | TYR: A: 81, ARG: A: 84, ILE: A: 73 | Hydrophobic interactions |
|  |  | THR: A: 85 | Electrostatic interactions |
| **CDK4** | Ribociclib (Reference compound) | ASP: A:163 | Hydrogen bonds |
|  |  | ILE: A:17, VAL: A:25, ALA: A:38, LYS: A:40, PHE: A:98, LEU: A:152, ALA: A:162 | Hydrophobic interactions |
|  |  | ASP: A:104 | Electrostatic interactions |
|  | α-Hederin | VAL: A:142, ARG: A:144 | Hydrogen bonds |
|  |  | ALA: A:21, LEU: A:64, ILE: A:141, VAL: A:142, ALA: A:167, VAL: A:181, TYR: A:196 | Hydrophobic interactions |
|  |  | ALA: A:167 | Unfavorable bond |
|  | Cytochalasin H | ARG: A:186 | Hydrogen bonds |
|  |  | GLN: A:173, ALA: A:175, VAL: A:181, LEU: A:183, VAL: A:190, LEU: A:191 | Hydrophobic interactions |
|  |  | MET: A:174 | Electrostatic interactions |
|  | Aurantiamide acetate | ALA: A:175, PRO: A:178, ARG: A:186 | Hydrogen bonds |
|  |  | ALA: A:21, VAL: A:181, LEU: A:183, ARG: A:186, GLU: A:224 | Hydrophobic interactions |
|  | Senkirkine | ALA: A:21, SER: A:171 | Hydrogen bonds |
|  |  | ALA: A:21, TYR: A:22, ARG: A:144, PRO: A:178 | Hydrophobic interactions |
|  | Thymoquinone | ALA: A: 175, PRO: A: 178 | Hydrogen bonds |
|  |  | TYR: A: 170, ALA: A: 175, PRO: A: 178 | Hydrophobic interactions |
| **CDK6** | Palcociclib (Reference compound) | LYS: A:43, ASP: A:102, ASP: A:104, GLN: A:149 | Hydrogen bonds |
|  |  | ILE: A:19, VAL: A:27, VAL: A:77, PHE: A:98, GLN: A:103, LEU: A:152, ALA: A:162, PHE: A:172 | Hydrophobic interactions |
|  |  | ASP: A:163 | Electrostatic interactions |
|  | Apigenin | HIS: A:100, VAL: A:101, ASP: A:163 | Hydrogen bonds |
|  |  | ILE: A:19, ALA: A:41, VAL: A:77, LEU: A:152, ALA: A:162 | Hydrophobic interactions |
|  | Chryseriol | HIS: A:100, VAL: A:101, ASP: A:163 | Hydrogen bonds |
|  |  | ILE: A:19, ALA: A:41, VAL: A:77, LEU: A:152, ALA: A:162 | Hydrophobic interactions |
|  | Linarigenin | VAL: A:101, ASP: A:163 | Hydrogen bonds |
|  |  | ILE: A:19, ALA: A:41, VAL: A:77, LEU: A:152, ALA: A:162 | Hydrophobic interactions |
|  | 5,7-Dihydroxy-3',4'-dimethoxyflavone | ILE: A:19, GLU: A:99, VAL: A:101, ASP: A:102, ASP: A:163 | Hydrogen bonds |
|  |  | ILE: A:19, ALA: A:41, VAL: A:77, LEU: A:152, ALA: A:162, PHE: A:172 | Hydrophobic interactions |
|  | Quercetin | HIS: A:100, ASP: A:102 | Hydrogen bonds |
|  |  | ILE: A:19, VAL: A:27, ALA: A:41, PHE: A:98, LEU: A:152 | Hydrophobic interactions |
|  | α- hederin | ASP: A: 102, HIS: A: 100, ILE: A: 19 | Hydrogen bonds |
|  |  | LYS: A: 111 | Hydrophobic interactions |
|  |  | VAL: A: 101 | Unfavorable bonds |
|  | Thymoquinone | LEU: A: 152, VAL: A: 101, PHE: A: 98, ALA: A: 162, VAL: A: 77, ALA: A: 41, VAL: A: 27 | Hydrophobic interactions |
|  | Tricin | ASP: A: 104, ASP: A: 102 | Hydrogen bonds |
|  |  | ILE: A: 19, PHE: A: 172, VAL: A: 27, ALA: A: 41, LEU: A: 152, ALA: A: 162, VAL: A: 77 | Hydrophobic interactions |
|  |  | ASP: A: 104 | Electrostatic interactions |
| **JAK1** | Cytochalasin H | ASP: A: 1021, ASN: A: 1008 | Hydrogen bonds |
|  |  | LEU: A: 881, LEU: A: 1010, ALA: A: 906, VAL: A: 889 | Hydrophobic interactions |
|  | Hemidine | ASN: A:1008, ASP: A:1039 | Hydrogen bonds |
|  |  | PHE: A:886, VAL: A:889, HIS: A:918, LEU: A:1024, | Hydrophobic interactions |
|  |  | ARG: A:1041  ASP: A:1039 | Unfavorable bond |
|  | Aurantiamide acetate | ARG: A: 1007, HIS: A: 885, PHE: A: 886 | Hydrogen bonds |
|  |  | GLU: A: 883, LEU: A: 922 | Hydrophobic interactions |
|  |  | GLY: A: 884, ASP: A: 1042 | Electrostatic interactions |
|  | Hemidesine | HIS: A: 918, ASN: A: 1008, ARG: A: 1007, GLY: A: 962 | Hydrogen bonds |
|  |  | PHE: A: 886, ALA: A: 906, LEU: A: 1010, VAL: A: 938, VAL: A: 889, LEU: A: 1024 | Hydrophobic interactions |
|  | β-Sitosterol | VAL: A: 889, LEU: A: 881, LEU: A: 1010, MET: A: 956, ALA: A: 906 | Hydrophobic interactions |
|  |  | PHE: A: 886 | Unfavorable interactions |
|  | Thymoquinone | LEU: A: 959 | Hydrogen bonds |
|  |  | ALA: A 906, MET: A: 956, VAL: A: 938, VAL: A: 889, LEU: A: 1010, LEU: A: 881 | Hydrophobic interactions |
| **JAK2** | Fedratinib (Reference compound) | LEU: A:855, GLY: A:856, GLU: A:930, LEU: A:932, SER: A:936 | Hydrogen bonds |
|  |  | VAL: A:863, ALA: A:880, VAL: A:911, MET: A:929, ARG: A:980, LEU: A:983 | Hydrophobic interactions |
|  |  | ASP: A:939 | Electrostatic interactions |
|  | Hemidine | SER: A:936, ASP: A:939 | Hydrogen bonds |
|  |  | LEU: A:855, PHE: A:860, VAL: A:863, LEU: A:983, LEU: A:997 | Hydrophobic interactions |
|  |  | ASN: A:859 | Unfavorable bond |
|  | Hydro vomifoliol | GLY: A:993, ASP: A:994 | Hydrogen bonds |
|  |  | LEU: A:855, VAL: A:863, ALA: A:880, LEU: A:983 | Hydrophobic interactions |
|  | Carvacrol | LEU: A:932 | Hydrogen bonds |
|  |  | LEU: A:855, VAL: A:863, ALA: A:880, MET: A:929, LEU: A:983 | Hydrophobic interactions |
|  | Dibutyl terephthalate | LEU: A:855, VAL: A:863, TYR: A:931, LEU: A:983 | Hydrophobic interactions |
|  | Thymoquinone | LEU: A:855, VAL: A:863, ALA: A:880, TYR: A:931, LEU: A:932, LEU: A:983 | Hydrophobic interactions |
|  | β-Sitosterol | GLU: A: 1015 | Hydrogen bonds |
|  |  | LEU: A: 997, VAL: A: 863, LEU: A: 983, LEU: A: 855 | Hydrophobic interactions |
| **SRC** | Dasatinib (Reference compound) | LEU: A:276, THR: A:341, MET: A:344 | Hydrogen bonds |
|  |  | VAL: A:284, ALA: A:296, ILE: A:339, TYR: A:343, LEU: A:396 | Hydrophobic interactions |
|  |  | LYS: A:298, ASP: A:351 | Electrostatic interactions |
|  | Chryseriol | MET: A:344, ASP: A:407 | Hydrogen bonds |
|  |  | VAL: A:284, ALA: A:296, LYS: A:298, VAL: A:326, THR: A:341, LEU: A:396, ALA: A:406, PHE: A:408 | Hydrophobic interactions |
|  | Linarigenin | LYS: A:298 | Hydrogen bonds |
|  |  | VAL: A:284, MET: A:317, VAL: A:326, ILE: A:339, THR: A:341, LEU: A:396, ALA: A:406, ASP: A:407, PHE: A:408, LEU: A:410 | Hydrophobic interactions |
|  | Quercetin | THR: A:341 | Hydrogen bonds |
|  |  | VAL: A:284, LYS: A:298, VAL: A:326, ILE: A:339, THR: A:341, LEU: A:396, ALA: A:406, ASP: A:407 | Hydrophobic interactions |
|  | Tricin | LEU: A:276, GLY: A:277, LYS: A:298, GLU: A:342, MET: A:344, ASP: A:351 | Hydrogen bonds |
|  |  | VAL: A:284, ALA: A:296, LEU: A:396 | Hydrophobic interactions |
|  | Cytochalasin H | LYS: A:298, ARG: A:391, ALA: A:393 | Hydrogen bonds |
|  |  | PHE: A:281, VAL: A:284, LEU: A:300, LEU: A:350, ALA: A:393, ILE: A:414 | Hydrophobic interactions |
|  | Thymoquinone | VAL: A: 284, LEU: A: 396, ALA: A: 296, ALA: A: 406, ILE: A: 339 | Hydrogen bonds |
|  |  | LYS: A: 298 | Hydrophobic interactions |

**The bond analysis of the docked compounds in relation to their respective target proteins**
